# Toward Stable Replication of Genomic Information in Pools of RNA Molecules

**DOI:** 10.1101/2024.07.17.603910

**Authors:** Ludwig Burger, Ulrich Gerland

## Abstract

The transition from prebiotic chemistry to living systems requires the emergence of a scheme for enzyme-free genetic replication. Here, we analyze a recently proposed prebiotic replication scenario, the so-called Virtual Circular Genome (VCG) [Zhou et al., RNA 27, 1-11 (2021)]: Replication takes place in a pool of oligomers, where each oligomer contains a subsequence of a circular genome, such that the oligomers encode the full genome collectively. While the sequence of the circular genome may be reconstructed based on long oligomers, monomers and short oligomers merely act as replication feedstock. We observe a competition between the predominantly error-free ligation of a feedstock molecule to a long oligomer and the predominantly erroneous ligation of two long oligomers. Increasing the length of long oligomers and reducing their concentration decreases the fraction of erroneous ligations, enabling high-fidelity replication in the VCG. Alternatively, the formation of erroneous products can be suppressed if each ligation involves at least one monomer, while ligations between two long oligomers are effectively prevented. This kinetic discrimination (favoring monomer incorporation over oligomer–oligomer ligation) may be an intrinsic property of the activation chemistry, or can be externally imposed by selectively activating only monomers in the pool. Surprisingly, under these conditions, shorter oligomers are extended by monomers more quickly than long oligomers, a phenomenon which has already been observed experimentally [Ding et al., JACS 145, 7504-7515 (2023)]. Our work provides a theoretical explanation for this behavior, and predicts its dependence on system parameters such as the concentration of long oligomers. Taken together, the VCG constitutes a promising scenario of prebiotic information replication: It could mitigate challenges of in non-enzymatic copying via template-directed polymerization, such as short lengths of copied products and high error rates.

## Introduction

In order to delineate possible pathways toward the emergence of life, it is necessary to understand how a chemical reaction network capable of storing and replicating genetic information might arise from prebiotic chemistry. RNA is commonly assumed to play a central role on this path, as it can store information in its sequence and catalyze its own replication (***Joyce, 1989***; ***Robertson and Joyce, 2012***; ***Higgs and Lehman, 2015***). While ribozymes capable of replicating strands of their own length have been demonstrated in the laboratory (***Attwater et al., 2013***), it remains elusive how enzyme-free self-replication might have worked before the emergence of such complex ribozymes.

One possible mechanism is template-directed primer extension (***Kervio et al., 2016***; ***Walton and Szostak, 2016***; ***Ding et al., 2021***; ***Leveau et al., 2022***; ***Welsch et al., 2023***): In this process, a primer hybridizes to a template and is extended by short oligonucleotides, thereby forming a (complementary) copy of the template strand. Considerable progress has been made in optimizing template-directed primer extension, but challenges remain: (i) The produced copies are likely to be incomplete. So far, at most 12 nt have been successfully added to an existing primer (***Leveau et al., 2022***). Moreover, as the pool of primer strands needs to emerge via random polymerization, the primer is likely to hybridize to the template at various positions, and not only to its 3^*′*^-end, leaving part of the 3^*′*^-end of the template uncopied (***Szostak, 2011***). (ii) Errors in enzyme-free copying are frequent due to the limited thermodynamic discrimination between correct Watson-Crick pairing and mismatches (***Kervio et al., 2010***; ***Leu et al., 2011, 2013***). While some activation chemistries (relying on bridged dinucleotides) have been shown to exhibit improved fidelity (***Duzdevich et al., 2021***), the error probability still constrains the length of the genome that can be reliably replicated.

The issue of insufficient thermodynamic discrimination can, in principle, be mitigated by making use of kinetic stalling after the incorporation of a mismatch (***Rajamani et al., 2010***; ***Leu et al., 2013***). By introducing a competition between the reduced polymerization rate and a characteristic timescale of the non-equilibrium environment, it is possible to filter correct sequences from incorrect ones (***Göppel et al., 2021***). To address the challenge of incomplete copies, Zhou *et al*. propose an new scenario of replication, the so-called Virtual Circular Genome (VCG) (***Zhou et al., 2021***). In this scenario, genetic information is stored in a pool of oligomers that are shorter than the circular genome they collectively encode: Each oligomer bears a subsequence of the circular genome, such that the collection of all oligomers encodes the full circular genome virtually. Within the pool, each oligomer can act as template or primer (***Zhou et al., 2021***). The oligomers hybridize to each other and form complexes that allow for templated ligation of two oligomers, or for the extension of an oligomer by a monomer. Because the sequences of the ligated strands and the template are part of the genome, most of the products should also retain the sequence of the genome. That way, long oligomers encoding the circular genome can be produced at the expense of short oligomers (***Zhou et al., 2021***). The long strands, in turn, can assemble into catalytically active ribozymes. With a continuous influx of short oligomers, the VCG might allow for continuous replication of the virtually encoded circular genome. Importantly, replication in the VCG is expected to avoid the issue of incomplete copies. Since the genome is circular, it does not matter which part of the genome an oligomer encodes, as long as the sequence is compatible with the genome sequence. An additional feature of the VCG scenario is that replication should be achievable without the need of adding many nucleotides to a primer: Provided the concentration of oligomers decreases exponentially with their length, the concentration of each oligomer in the pool can be doubled by extending each oligomer only by a few nucleotides (***Zhou et al., 2021***). The extension of an oligomer by a few nucleotides in a VCG pool has already been demonstrated experimentally (***Ding et al., 2023***).

A recent computational study points out that the VCG scenario is prone to loss of genetic information via “sequence scrambling” (***Chamanian and Higgs, 2022***): If the genome contains identical sequence motifs at multiple different loci, replication in the VCG will mix the sequences of these loci, thus destroying the initially defined genome. It is currently unclear, which conditions could prevent this genome instability of VCGs, such that their genetic information is retained. Length distribution, sequence composition, oligonucleotide concentration and environmental conditions, such as temperature, all affect the stability of complexes and thus the replication dynamics of the VCG pool. Here, we characterize the replication fidelity and yield of VCG pools using a kinetic model, which explicitly incorporates association and dissociation of RNA strands as well as templated ligation. We study a broad spectrum of prebiotically plausible and experimentally accessible oligomer pools, from pools containing only monomers and long oligomers of a single length to pools including a range of long oligomers with uniform or exponential concentration profile. The length of the included oligomers as well as their concentration are free parameters of our model.

We find that, regardless of the pool composition, three competing types of template-directed ligation reactions emerge: (i) ligations between two short oligomers (or monomers), producing products too short to specify a unique genomic locus, (ii) ligations between a short and a long oligomer, typically generating longer products compatible with the genome sequence, and (iii) ligations between two long oligomers, which often yield sequences incompatible with the genome. These erroneous ligation of type (iii) are a key driver of sequence scrambling, as they covalently link oligomers originating from non-adjacent genomic loci, effectively “mixing” distant regions of the genome. Fidelity is primarily determined by the competition between the correct extension of a long oligomer and the erroneous ligation of two long oligomers. The likelihood of the latter can be reduced by decreasing the relative abundance of long oligomers, even though this increases the frequency of unproductive ligations between short oligomers. As a result, fidelity can be improved at the cost of reduced yield. The efficiency, meaning the yield attainable at a fixed high fidelity, thus depends on the length distribution of the oligomers in the pool.

Alternatively, the issue of erroneous ligations is mitigated if ligations of long oligomers are kinetically suppressed, such that each ligation incorporates only one monomer at a time, as in the experimental study by Ding *et al*. (***Ding et al., 2023***). In this case, the VCG concentration can be chosen arbitrarily large without compromising fidelity. Interestingly, our model predicts an unexpected feature: In the limit of high VCG concentrations, short oligomers are more likely to be extended than long oligomers, even though, intuitively, complexes containing longer oligomers are expected to be more stable and thus more productive. The same behavior was indeed observed experimentally (***Ding et al., 2023***). We provide an explanation for this feature, and discuss its dependence on system parameters such as the length and the concentration of long oligomers in the pool.

### Model

In the VCG scenario, a circular genome is stored in a pool of oligomers, with each oligomer shorter than the genome it helps encode. Each oligomer bears a subsequence of the circular genome, such that, collectively, the oligomers encode the full genome (Fig. 1A). As the spontaneous emergence of such VCG pools is expected to be rare (***Chamanian and Higgs, 2022***), our study focuses on the conditions under which an existing VCG pool can reliably replicate. We therefore begin with a known genome and an associated VCG pool, without addressing the question of origin. To set up our model of VCG dynamics, we specify (i) the circular genome used, (ii) the procedure by which the genome is mapped to a set of oligomers, and (iii) the chemical reactions governing the system’s evolution.

**Figure 1.**
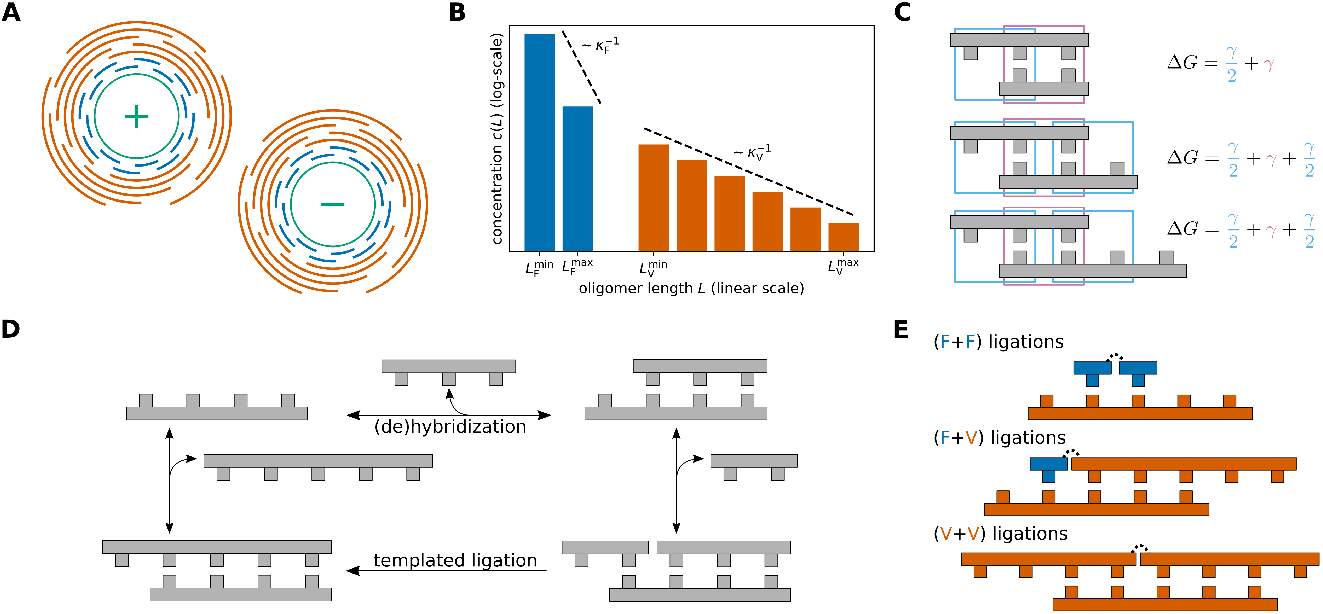
Model. **(A)** In the Virtual Circular Genome (VCG) scenario, a circular genome (depicted in green) as well as its sequence complement are encoded in a pool of oligomers (depicted in blue and orange). Collectively, the pool of oligomers encodes the whole sequence of the circular genome. Depending on their length, two types of oligomers can be distinguished: Long VCG oligomers specify a unique locus on the genome, while feedstock molecules (monomers or short oligomers) are too short to do so. **(B)** The length-distribution of oligomers included in the VCG pool is assumed to be exponential. The concentration of feedstock and VCG oligomers as well as their respective length scales of exponential decay 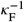 and 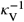 can be varied independently. The set of included oligomer lengths can be restricted via 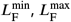 and 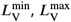. **(C)** The hybridization energy of complexes is computed using a simplified nearest-neighbor model: Each full block comprised of two base pairs (depicted in pink) contributes *γ*, while dangling end blocks (depicted in blue) contribute *γ*/2. **(D)** Oligomers form complexes via hybridization reactions, or dehybridize from an existing complex. The ratio of hybridization and dehybridization rate is governed by the hybridization energy (Eq. 3). If two oligomers are adjacent to each other in a complex, they can undergo templated ligation. **(E)** Based on the length of the reacting oligomers, we distinguish three types of templated ligation: Ligation of two feedstock molecules (F+F), ligation of a feedstock molecule to a VCG oligomer (F+V) and ligation of two VCG oligomers (V+V).

### Circular Genomes

For a given genome length, *L*_*G*_, there are 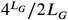 distinct circular genomes (the factor 1/2*L*_*G*_ accounts for the freedom to select the starting position and to choose the Watson or the Crick strand as reference sequence). A key property of the genome is its unique motif length, *L*_*U*_, defined as the shortest length such that all possible subsequences of length *L* ≥ *L*_*U*_ appear at most once in the genome. In other words, all subsequences of length *L* ≥ *L*_*U*_ specify a unique locus on the genome. In addition, each circular genome has another length scale, corresponding to the maximal motif length, up to which all possible motifs are contained in the genome. We refer to this length as the exhaustive coverage length, *L*_*E*_. We typically analyze unbiased genomes in which all possible subsequences of length *L* ≤ *L*_*E*_ are contained at equal frequency.

### Construction of VCG Pools

To specify a VCG pool that encodes a genomic sequence, one must select which subsequences are included in the pool at which concentrations. We consider unbiased pools, where the concentration of subsequences, *c(L)*, depends only on their length, *L*, i.e., all subsequences of a given length are included at equal concentration. We refer to the length-dependent concentration profile as the length distribution of the pool. Depending on their length, oligomers fall into two categories (Fig. 1B): (i) short feedstock molecules (monomers and oligomers) and (ii) long VCG oligomers. Feedstock oligomers are oligomers that are shorter than the unique motif length *L*_*U*_. Since their sequence appears multiple times on the genome, they do not encode a specific position on the genome. Thus, they serve as feedstock for the elongation of VCG oligomers rather than as information storage. Conversely, VCG oligomers, which are longer than the unique motif length *L*_*U*_, have a unique locus on the circular genome. Collectively, the VCG oligomers enable the reconstruction of the full genome.

The full length distribution, *c(L)*, can be decomposed into the contributions of feedstock and VCG oligomers,

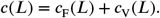

We assume that both *c*_F_ and *c*_V_ follow an exponential length distribution. In our model, the concentration of VCG oligomers can be varied independently of the concentration of feedstock, and the length scales for the exponential decay (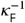 vs. 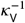) may differ between feedstock and VCG oligomers. Additionally, we can restrict the set of oligomer lengths included in the pool by setting minimal and maximal lengths for feedstock and VCG oligomers individually,

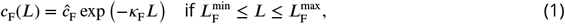

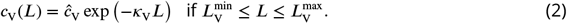

For any other oligomer length, the concentrations equal zero. This parametrization includes uniform length distributions as a special case (*κ*_F_ = 0 and *κ*_V_ = 0), and also allows for concentration profiles that are peaked. Peaked length distributions can emerge from the interplay of templated ligation, (de)hybridization and outflux in open systems (***Rosenberger et al., 2021***). We define the total concentration of feedstock, 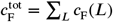, as well as the total concentration of VCG oligomers, 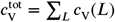. Their ratio will turn out to be an important determinant of the VCG dynamics.

### (De)hybridization Kinetics

Oligomers can hybridize to each other to form double-stranded complexes, or dehybridize from an existing complex. For simplicity, we do not include self-folding within a strand, which is a reasonable assumption for short oligomers. The stability of a complex is determined by its hybridization energy, with lower hybridization energy indicating greater stability. We use a simplified nearest-neighbor energy model to compute the hybridization energy (***Rosenberger et al., 2021***; ***Göppel et al., 2022***; ***Laurent et al., 2024***): The total energy equals the sum of the energy contributions of all nearest-neighbor blocks in a given complex (Fig. 1C). The energy contribution associated with a block of two Watson-Crick base pairs (matches) is denoted *γ* < 0, and dangling end blocks involving one Watson-Crick pair and one free base contribute *γ*/2. Nearest-neighbor blocks with mismatches increase the hybridization energy by *γ*_*MM*_ > 0 per block, thus reducing the stability of the complex. The rate constants of hybridization and dehybridization are related via

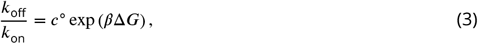

where *c*^°^ = 1 M is the standard concentration, and *ΔG* is the free energy of hybridization. The association rate constant *k*_on_ is proportional to the encounter rate constant, *k*_enc_ = 1/*(c*^°^*t*_0_). The encounter timescale *t*_0_ serves as the elementary time unit of the kinetic model, with all reaction timescales measured relative to it.

### Templated Ligation

Two oligomers A and B that are hybridized adjacently to each other on a third oligomer C can produce a new oligomer A-B via templated ligation (Fig. 1D). Depending on the length of A and B, we distinguish three types of ligation reactions (Fig. 1E): (i) F+F ligations, in which two feedstock molecules ligate, (ii) F+V ligations, where a VCG oligomer is extended by a feedstock molecules, and (iii) V+V ligations involving two VCG oligomers. The formation of a covalent bond via templated ligation is not spontaneous, but requires the presence of an activation reaction. Usually, these reactions add a leaving group to the 5^*′*^-end of the nucleotide, which is cleaved during bond formation (***Kervio et al., 2016***; ***Walton and Szostak, 2016***). We assume that the concentration of activating agent is sufficiently high for the activation to be far quicker than the formation of the covalent bond, such that activation and covalent bond formation can be treated as a single effective reaction. When not otherwise stated, we assume that all possible templated ligation reactions occur with the same rate constant *k*_*lig*_.

### Observables

Templated ligation in the pool forms longer oligomers at the expense of shorter oligomers and monomers. While the product of an F+V ligation (or V+V ligation) is always a VCG oligomer, F+F ligations can produce feedstock or VCG oligomers. In both cases, a produced VCG oligomer can be correct (compatible with the genome) or incorrect (incompatible). We quantify these processes by measuring extension fluxes in units of nucleotides ligated to an existing strand (counting the length of the shorter ligated strand as the number of incorporated bases). In particular, we define the fidelity *f* as the extension flux resulting in correct VCG oligomers relative to the flux resulting in any VCG oligomer,

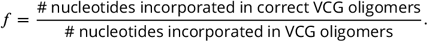

In addition, we introduce the yield *y* as the proportion of total extension flux that produces VCG oligomers,

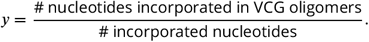

Efficient replication of the VCG requires both, high fidelity and high yield. Hence, we introduce the efficiency of replication *η* as the product of fidelity and yield,

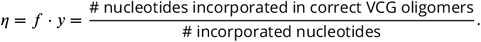

Moreover, we define the ligation share *s* of a ligation type, which allows us to discern the contributions of different types of templated ligations (F+F, F+V, V+V) to fidelity, yield, and efficiency,

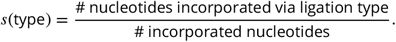

## Results

### Replication efficiency reaches a maximum at intermediate concentrations of VCG oligomers

We begin our analysis of the dynamics of VCG pools with an exemplary genome of length *L*_*G*_ = 16 nt,

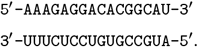

This genome contains all possible monomers and dimers with equal frequency, ensuring that all motifs up to *L*_*E*_ = 2 nt are represented. Identifying a unique address on this genome requires at least three nucleotides. Therefore, the unique motif length is *L*_*U*_ = 3 nt, and VCG oligomers need to be at least 3 nt long. Further below, we also explore genomes of different lengths *L*_*G*_, as well as varying characteristic sequence length scales *L*_*E*_ and *L*_*U*_ (method for genome construction detailed in Supplementary Section S1).

Based on the genome, we construct the initial oligomer pool. As a first step, we focus on a simple scenario in which the pool contains only monomers (serving as feedstock) and VCG oligomers of a single, defined length. The VCG pools are evolved in time using a stochastic simulation based on the Gillespie algorithm (***Rosenberger et al., 2021***; ***Göppel et al., 2022***; ***Laurent et al., 2024***). Since the Gillespie algorithm operates on the level of counts of molecules instead of concentrations, we must assign a volume to each system (in the range 1 µm^3^ to 10 000 µm^3^, see Supplementary Material Section S2). Besides the volume, we also need to choose the reaction rate constants appropriately: (i) The association time *t*_0_ is the fundamental time unit in our kinetic model, and all other times are expressed relative to *t*_0_. Experimentally determined association rate constants are typically around 10^6^ −10^7^ M^−1^ s^−1^ (***Wetmur and Davidson, 1968***; ***Braunlin and Bloomfield, 1991***; ***Ashwood et al., 2023***; ***Todisco et al., 2024a***). For the purpose of estimating absolute timescales, we assume a constant association timescale of *t*_0_ ≈ 1 µ*s* in the following. (ii) The timescale of dehybridization is computed via Eq. (3) using the energy contribution *γ =* −2.5 *k*_*B*_*T* for a matching nearest-neighbor block and *γ*_MM_ = 25.0 *k*_*B*_*T* in case of mismatches. The high energy penalty of nearest-neighbor blocks involving mismatches, *γ*_MM_, is chosen to suppress the formation of mismatches, while the value of *γ* roughly matches the average energy of all matching nearest-neighbor blocks given by the Turner energy model of RNA hybridization (***Mathews et al., 2004***). (iii) For templated ligation, we select a reaction rate constant of 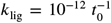. This choice of *k*_*lig*_ is consistent with ligation rates measured in enzyme-free template-directed primer extension experiments, which range from around 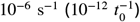 (***Sosson et al., 2019***; ***Leveau et al., 2022***) to roughly 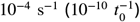 (***Walton and Szostak, 2017***), depending on the underlying activation chemistry. This indicates that, for sufficiently short oligomers (up to about 11 nt long), hybridization and dehybridization occur much faster than ligation, so binding equilibrates before ligation takes place.

Based on the ligation events observed in the simulation, we compute the observables introduced above (Model section). Due to the small ligation rate constant, it is computationally unfeasible to simulate the time evolution for more than a few ligation time units. Consequently, ligation events are scarce, which leads to high variances in the computed observables. We mitigate this issue by calculating the observables based on the concentration of complexes that are in a productive configuration, even if they do not undergo templated ligation within the time window of the simulation (Supplementary Material Section S2).

From the ligation events, we compute the total flux of oligomer formation. The observable *y* (yield) quantifies the fraction of this total flux directed to producing VCG oligomers. Fig. 2B shows how the yield depends on the concentration of VCG oligomers, 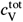, at a fixed total monomer concentration, 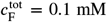. Here, the data points (with error bars) represent the simulation results with the different colors corresponding to different choices of initial VCG oligomer length, *L*_V_. We observe that the yield increases monotonically with the concentration of VCG oligomers, and approaches 100% for high 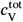. The concentration at which the pool reaches a yield of 50% depends on the oligomer length *L*_V_: Shorter VCG oligomers require higher concentrations for high yield. The concentration dependence of the yield can be rationalized by the types of templated ligations that are involved. For low VCG concentration, most templated ligation reactions are dimerizations (1+1) with the VCG oligomers merely acting as templates (Fig. 2C). As dimers are shorter than the unique motif length *L*_*U*_, their formation does not contribute to the yield, which explains the low yield in the limit of small 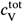. Conversely, for high VCG concentration, most of the templated ligations are F+V or V+V ligations, which produce oligomers of length *L* ≥ *L*_*U*_, implying high yield.

**Figure 2.**
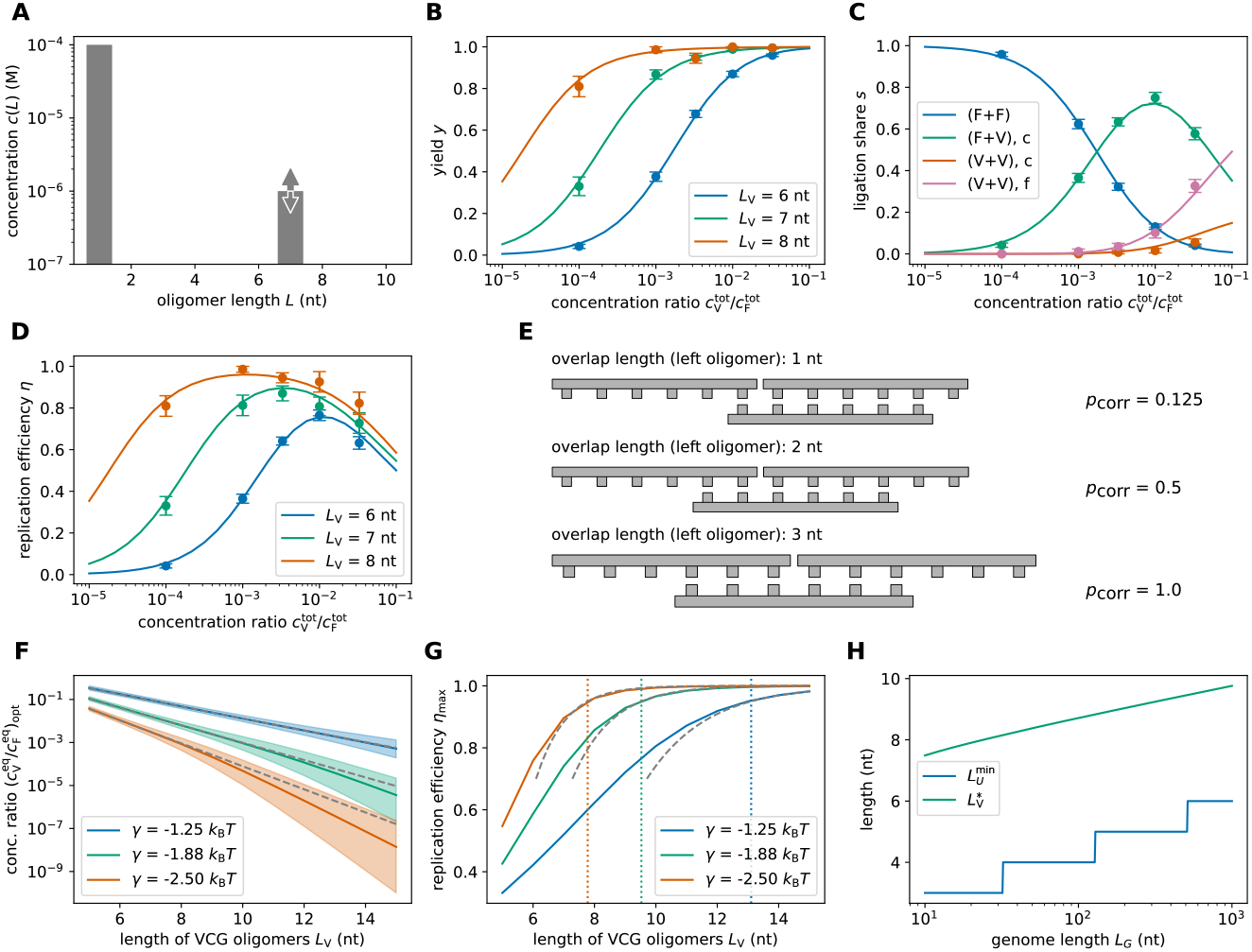
Replication performance of VCG pools containing VCG oligomers of a single length (single-length VCG pools). **(A)** The pool contains a fixed concentration of monomers, 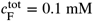, as well as VCG oligomers of a single length, *L*_V_, at variable concentration 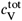 (the VCG oligomers cover all possible subsequences of the genome and its complement at equal concentration). **(B)** The yield increases as a function of 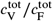, because dimerizations become increasingly unlikely for high VCG concentrations. **(C)** The ligation share of different ligation types depends on the total VCG concentration: In the low concentration limit, dimerization (F+F) dominates; for intermediate concentrations, F+V ligations reach their maximum, while, for high concentrations, a substantial fraction of reactions are V+V ligations. The panel depicts the behavior for *L*_V_ = 6 nt. **(D)** Replication efficiency is limited by the small yield for small 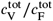. In the limit of high 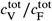, replication efficiency decreases due to the growing number of error-prone V+V ligations. Maximal replication efficiency is reached at intermediate VCG concentration. **(E)** V+V ligations are prone to the formation of incorrect products due to the short overlap between educt strand and template. In general, the probability of correct product formation, *p*_corr_, depends on the choice of circular genome and as well as its mapping to the VCG pool. The probabilities listed here refer to a VCG pool with *L*_*G*_ = 16 nt, *L*_*E*_ = 2 nt and *L*_*U*_ = 3 nt. **(F)** The optimal equilibrium concentration ratio of free VCG strands to free feedstock strands, which maximizes replication efficiency, decays as a function of length (continuous line). The analytical scaling law (dashed line, Eq. (4)) captures this behavior. The window of close-to-optimal replication, within which efficiency deviates no more than 1% from its optimum (shaded areas), increases with *L*_V_, facilitating reliable replication without fine-tuning to match the optimal concentration ratio. **(G)** Maximal replication efficiency, which is attained at the optimal VCG concentration depicted in panel E, increases as a function of *L*_V_ and approaches a plateau of 100%. For high efficiency, Eq. (5) provides a good approximation of the length-dependence of *η*_max_ (dashed lines). The oligomer length at which replication efficiency equals 95% is determined using Eq. (5) (vertical dotted lines). **(H)** The unique motif length, 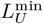, increases logarithmically with the length of the genome, *L*_*G*_. The length of VCG oligomers, *L*_V_, at which the optimal replication efficiency reaches 95% (computed using Eq. (5)) exhibits the same logarithmic dependence on *L*_*G*_.

Fig. 2C also shows that the relative contribution of V+V ligations increases with increasing 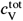, with a large fraction of them producing incorrect products (denoted V+V,f). This reduces the fidelity of replication, *f*, and causes a trade-off between fidelity and yield, which causes replication efficiency, *η = f* · *y*, to reach a maximum at intermediate VCG concentrations(Fig. 2D). Using hexamers as an example, Fig. 2E illustrates why V+V ligations are prone to forming incorrect oligomers: In order to ensure that two oligomers only ligate if their sequences are adjacent to each other in the true circular genome, both oligomers need to share an overlap of at least *L*_*U*_ nucleotides with the template oligomer. Otherwise, the hybridization region is too short to identify the locus of the oligomer uniquely, and two oligomers from non-adjacent loci might ligate. The probability of forming incorrect products is a consequence of the combinatorics of possible subsequences in the VCG. Specifically, for a genome with *L*_*G*_ *=* 16 nt, there are 32 different hexamers. For example, if the left educt hexamer only has one nucleotide of overlap with the template, there are 1/4 · 32 = 8 possible educt oligomers. However, only one out of those 8 hexamers is the correct partner for the right educt hexamer, implying an error probability of 1 − 1/8 = 7/8 (first example in Fig. 2E).

Characterizing replication efficiency via full simulations is computationally expensive: Depending on parameters, obtaining a single data point in Fig. 2B–D can require hundreds of simulations, each lasting several days. To explore a broad parameter space more easily, we introduce an approximate adiabatic method that (i) assumes ligation is much slower than any hybridization or dehybridization event, and (ii) relies on a coarse-grained sequence-independent representation of oligomers. Details are provided in Supplementary Material Sections S3 and S4. In brief, because ligation is rare, we first compute the equilibrium distribution of free and bound oligomers. Oligomers of the same length share a common concentration, and complex concentrations are determined via the mass action law using length-dependent dissociation constants. Combining the mass action law with a mass conservation constraint for each oligomer length allows us to compute the equilibrium concentrations of free VCG and feedstock strands, 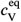 and 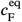, given total concentrations 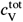 and 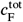. With these equilibrium values, we determine the concentrations of productive complexes and thus obtain the desired ligation-based observables without running full stochastic simulations.

The results of the adiabatic approach agree well with the simulation data (Fig. 2B-D), supporting that the replication efficiency depends non-monotonously on the concentration of VCG oligomers, with a maximum at intermediate concentration. While the available simulation data only allows for a qualitative characterization of this trend, the adiabatic approach enables a quantitative analysis. For instance, we use the adiabatic approach to determine the equilibrium concentration ratio at which replication efficiency is maximal as a function of the VCG oligomer length *L*_V_ (solid line in Fig. 2F). As expected from the qualitative trend observed in the simulation, pools containing longer oligomers reach their maximum for lower concentration of VCG oligomers. The shaded area indicates the range of VCG concentrations within which the pool’s efficiency deviates by no more than one percent from its optimum. We observe that this range of close-to-optimal VCG concentrations increases with *L*_V_. Thus, pools containing longer oligomers require less fine-tuning of the VCG concentration for replication with high efficiency.

In addition to the numerical results, we utilize the adiabatic approach to study the optimal equilibrium VCG concentration analytically (Supplementary Material Section S6). We find that, for any choice of *L*_V_, replication efficiency reaches its maximum when the fractions of dimerization (1+1) reactions and erroneous V+V ligations are equal (Fig. 2C for *L*_V_ *=* 6 nt). This criterion can be used to derive a scaling law for the optimal equilibrium concentration ratio 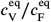 as a function of the oligomer length *L*_V_,

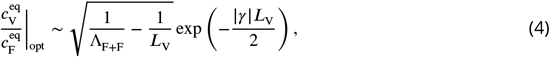

which is shown as a dashed line in Fig. 2F (the length-scale Λ_F+F_ is defined in Supplementary Material Section S6). The optimal equilibrium concentration ratio decreases exponentially with *L*_V_, while the hybridization energy *γ*/2 sets the inverse length-scale of the exponential decay. Analytical estimate and numerical solution agree well, as long as the hybridization is weak and oligomers are sufficiently short. For strong binding and long oligomers, complexes involving more than three strands play a non-negligible role, but such complexes are neglected in the analytical approximation.

Fig. 2G shows how the maximal replication efficiency depends on the VCG oligomer length. Consistent with the qualitative trend observed in Fig. 2D, longer oligomers enable higher maxima in replication efficiency. Regardless of the choice of *γ*, replication efficiency reaches 100% if *L*_V_ is sufficiently high. Starting from Eq. (4), we find the following approximation for the maximal replication efficiency attainable at a given oligomer length (dashed lines in Fig. 2G),

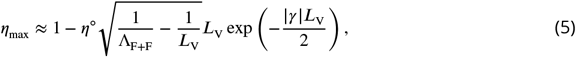

where *η*^°^ is a genome-dependent constant (Supplementary Material Section S7). This equation can provide guidance for the construction of VCG pools with high replication efficiency: Given a target efficiency, the necessary oligomer length and hybridization energy, i.e., temperature, can be calculated. In Fig. 2G, we determine the oligomer length necessary to achieve *η*_max_ *=* 95*%* for varying hybridization energies *γ*. At higher temperature (weaker binding), VCG pools require longer oligomers to replicate with high efficiency.

Eq. (5) is not restricted to the specific example genome of length *L*_*G*_ *=* 16 nt, but applies more generally to genomes of arbitrary length. Any genome of length *L*_*G*_ can contain at most 2 *L*_*G*_ distinct motifs. Consequently, the minimum length required to specify a unique address on the genome equals 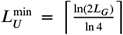. By the same logic, the longest motif length for which all 4^*L*^ possible sequences can be exhaustively represented within the genome is 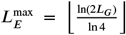.Both of these characteristic lengths scale logarithmically with genome size. For genomes where *L*_*E*_ is taken to be maximal and *L*_*U*_ minimal, we find that the characteristic VCG oligomer length 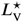 required for replication with high efficiency (*η*_max_ *= 95%*) also scales logarithmically with genome length (Fig. 2H). Across genome lengths, the offset between *L*_*U*_ and 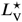 is roughly constant.

In genomes with different choices of *L*_*E*_ and *L*_*U*_ (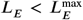 and 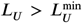), the characteristic VCG oligomer length required for efficient replication, 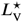, is primarily determined by the unique motif length *L*_*U*_. Specifically, the length of VCG oligomers, *L*_V_, must exceed *L*_*U*_, regardless of the value of *L*_*E*_. This is shown for genomes of length *L*_*G*_ *=* 64 nt in Supplementary Material Section S8, where we generate genomes with specified length scales *L*_*E*_ and *L*_*U*_ using a Metropolis–Hastings algorithm and analyze their replication efficiency. Intuitively, the required VCG oligomer length 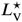 is set by *L*_*U*_, because *L*_*U*_ defines the minimal hybridization region needed to ensure correct ligation via sequence-specific recognition. The precise offset between *L*_*U*_ and 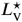 depends on genome-specific features, such as the frequency distribution of motifs with lengths between *L*_*E*_ and *L*_*U*_. If this distribution is nearly uniform (noting that at least one motif must repeat, or else it would constitute a unique address), then 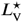 will be close to *L*_*U*_ (Supplementary Material Fig. S9B). In contrast, strongly biased motif distributions require larger 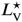 to achieve reliable replication (Supplementary Material Fig. S9A), though even in this case, 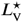 typically exceeds *L*_*U*_ by only a few nucleotides.

### Multi-length VCG pools are dominated by longest oligomers

In the previous section, we characterized the behavior of pools containing VCG oligomers of a single length. We observed that V+V ligations are error-prone due to insufficient overlap between the educt strands and the template, whereas F+V ligations extend the VCG oligomers with high efficiency. The F+V ligations will gradually broaden the length distribution of the VCG pool, raising the question of how this broadening affects the replication behavior. In principle, introducing multiple oligomer lengths into the VCG pool might even improve the fidelity of V+V ligations, since a long VCG oligomer could serve as a template for the correct ligation of two shorter VCG oligomers.

To analyze this question quantitatively, we first consider the simple case of a VCG pool that only contains monomers, tetramers, and octamers. The concentration of monomers is set to *c*(1) = 0.1 mM, while the concentrations of the VCG oligomers are varied independently (Fig. 3A). Replication efficiency reaches its maximum at *c*(8) ≈ 0.1 µM and very low tetramer concentration, *c*(4) ≈ 7.4 pM, effectively resembling a single-length VCG pool containing only octamers. As shown in Fig. 3B, the maximal efficiency is surrounded by a plateau of close-to-optimal efficiency. The octamer concentration can be varied by more than one order of magnitude without significant change in efficiency. Similarly, adding tetramers does not affect efficiency as long as the tetramer concentration does not exceed the octamer concentration. Fig. 3C illustrates that the plateau of close-to-optimal efficiency coincides with the concentration regime where the ligation of a monomer to an octamer with another octamer acting as template, 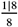, is the dominant ligation reaction. For high tetramer concentration and intermediate octamer concentration, templated ligation of tetramers on octamer templates, 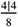, surpasses the contribution of 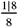 ligations (green shaded area in Fig. 3C). The 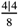 reactions give rise to a ridge of increased efficiency, which, however, is small compared to the plateau of close-to-optimal efficiency (Fig. 3B). Even though reactions of the type 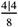 produce correct products in most cases, they compete with error-prone ligations like 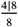 or 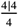, which reduce fidelity. Increasing the binding affinity, i.e., lowering *γ*, enhances the contribution of 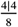 ligations at the expense of 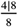 ligations, resulting in an increased local maximum in replication efficiency (Supplementary Material Fig. S11). However, very strong hybridization energy, *γ =* −5.0 *k*_*B*_*T*, is necessary for 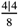 ligations to give rise to similar efficiency as 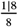 ligations. Thus, in this example, high efficiency replication facilitated by the ligation of short VCG oligomers on long VCG oligomers is in theory possible, but requires unrealistically strong binding affinity. Moreover, this mechanism is only effective if educts and template differ significantly with respect to their length. If the involved VCG oligomers are too similar in size, templated ligation of two short oligomers tends to be error-prone, irrespective of the choice of binding affinity (Supplementary Material Fig. S12 and S13). In that case, F+V ligations remain the most reliable replication mechanism.

**Figure 3.**
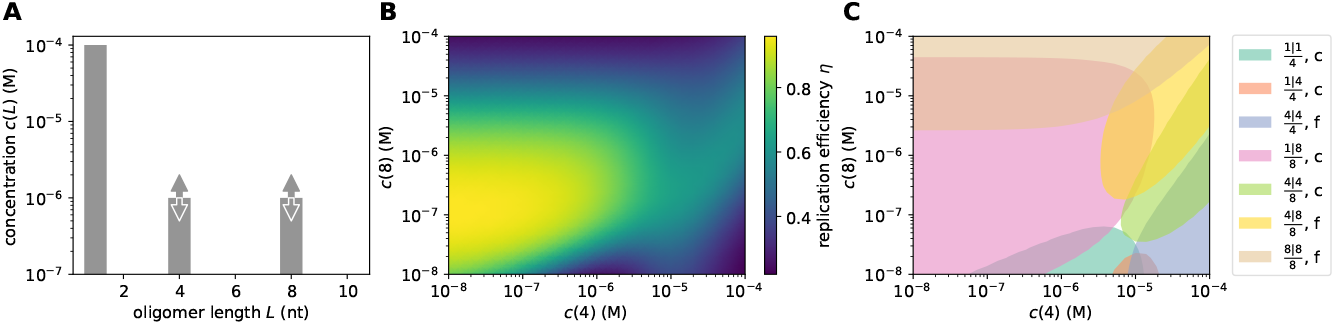
Replication performance of pools containing VCG oligomers of two different lengths. **(A)** The pool contains a fixed concentration of monomers, 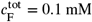, as well as tetramers and octamers at variable concentration. The hybridization energy per nearest-neighbor block is *γ* = −2.5 *k*_B_*T*. **(B)** Replication efficiency reaches its maximum for *c*(8) ≈ 0.1 µM and significantly lower tetramer concentration, *c*(4) ≈ 7.4 pM. Efficiency remains close-to-maximal on a plateau around the maximum spanning almost two orders of magnitude in tetramer and octamer concentration. In addition, efficiency exhibits a ridge of increased efficiency for high tetramer concentration and intermediate octamer concentration. **(C)** Complexes that facilitate templated ligation are grouped by the length of the template and the educts, 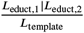. We distinguish complexes producing correct (labeled “c”) and false products (labeled “f”). For each relevant type of complex, we highlight the region in the concentration plane where it contributes most significantly, i.e., at least 20% of the total ligation flux. The plateau of high efficiency is dominated by the ligation of monomers to octamers, whereas the ridge of increased efficiency is due to the correct ligation of two tetramers templated by an octamer.

We observe similar behavior in VCG pools containing a range of oligomer lengths from 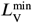 to 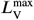, rather than just tetramers and octamers. For a uniform VCG concentration profile (Fig. 4A, dark grey), the maximal replication efficiency (at the optimal VCG concentration) is attained when F+V ligations involving long VCG oligomers dominate the templated ligation (see Fig. 4D, showing the contribution of different ligation types in pools with varying 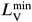 at fixed 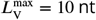, using *γ =* −2.5 *k*_*B*_*T*). Importantly, the maximal replication efficiency is always bounded by the efficiency of the longest VCG oligomer in the pool, independent of the presence or length of shorter oligomers (blue curve in Fig. 4B; identical to the orange curve in Fig. 2G). Including short VCG oligomers has minimal effect on the dominant ligation types and only slightly increases the proportion of unproductive F+F ligations. Consequently, reducing 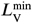 while keeping 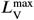 fixed leads to only a modest decline in maximal efficiency. In contrast, decreasing 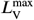 while holding 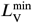 constant causes a substantial reduction in efficiency (Fig. 4B), because the longest oligomer in the pool sets an upper bound on replication efficiency. Moreover, at low 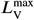, short and long oligomers become more similar in length, giving rise to a spectrum of erroneous V+V ligations that compete with the productive F+V ligations (Fig. 4E).

**Figure 4.**
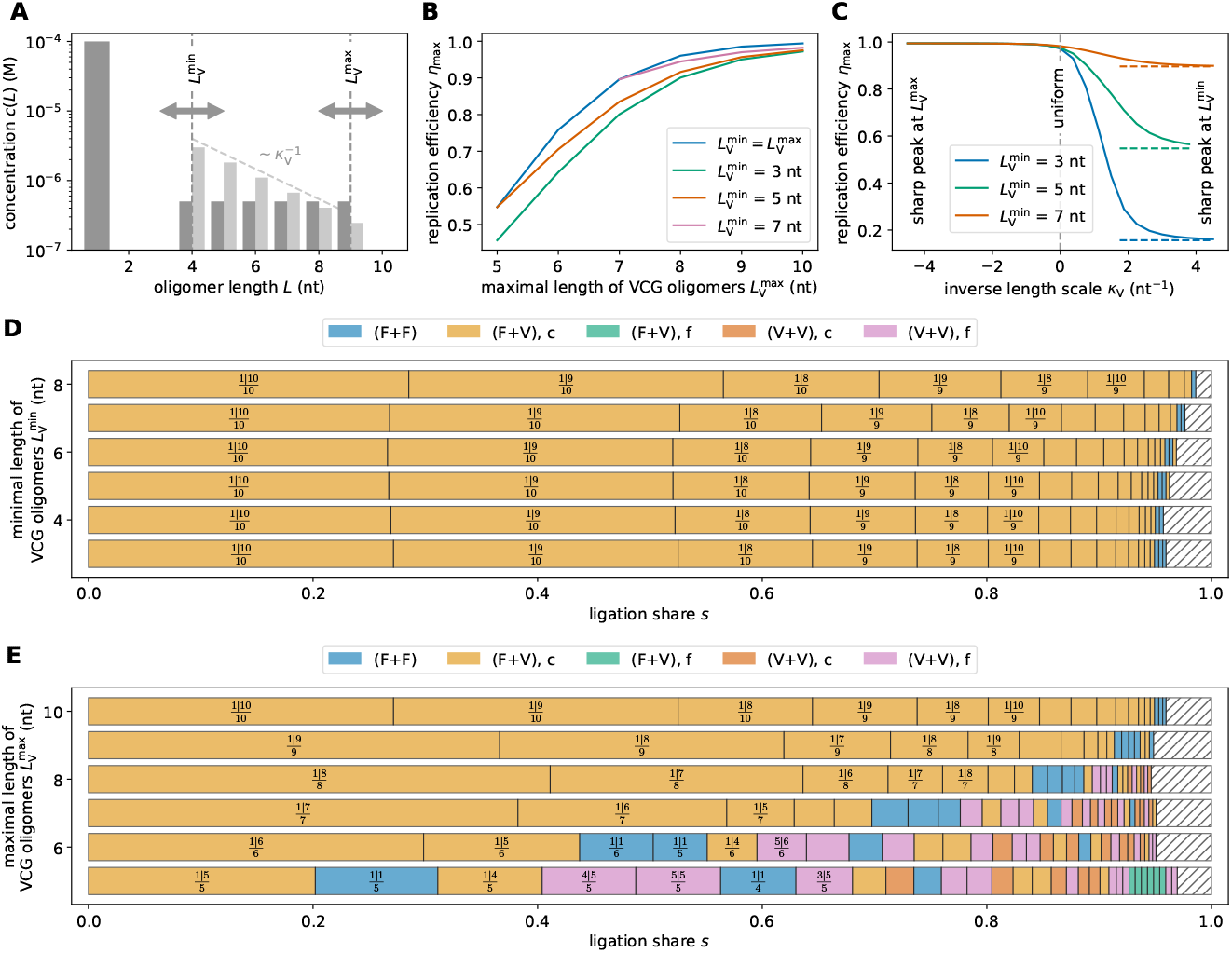
Replication performance of multi-length VCG pools. **(A)** The pool contains a fixed concentration of monomers, 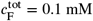, as well as long oligomers in the range 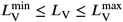 at variable concentration 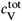. The length dependence of the concentration profile is assumed to be uniform (for panels B, D, and E) or exponential (for panel C); its steepness is set by the parameter *κ*_V_. **(B)** If the length distribution is uniform, reducing 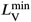 decreases the maximal efficiency, whereas increasing 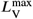 increases it. Pools containing a range of oligomer lengths are always outperformed by single-length VCGs (blue curve). **(C)** Assuming an exponential length distribution of VCG oligomers allows us to tune from a poorly-performing regime (dominated by oligomers of length 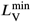) to a well-performing regime (dominated by oligomers of length 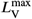). In the limit *κ*_V_ → ∞, *η*_max_ approaches the replication efficiency of single-length pools containing only oligomers of length 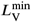 (dashed lines). **(D)** For high 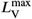, replication is dominated by primer extension of the long oligomers in the VCG (here 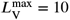). In this limit, addition of shorter oligomers leaves the dominant F+V ligations almost unchanged. **(E)** Reducing 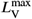 for fixed 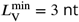 increases the fraction of unproductive (i.e., dimerization) or erroneous ligation reactions.

In a realistic prebiotic scenario, the concentration profile of the VCG pool would not be uniform. Depending on the mechanism producing the pool and its coupling to the non-equilibrium environment, it might have a concentration profile that decreases or increases exponentially with length. We use the parameter *κ*_V_ to control this exponential length dependence (Fig. 4A): For negative *κ*_V_, the concentration increases as a function of length, while exponentially decaying length distributions have positive *κ*_V_. We find that replication efficiency is high if the concentration of long VCG oligomers exceeds or at least matches the concentration of short VCG oligomers (*κ*_V_ ≤ 0 in Fig. 4C). In that case, replication efficiency is dominated by the long oligomers in the pool, since these form the most stable complexes. As the concentration of long oligomers is decreased further (*κ*_V_ > 0), the higher stability of complexes formed by longer oligomers is eventually insufficient to compensate for the reduced concentration of long oligomers. Replication efficiency is then governed by short VCG oligomers, which exhibit lower replication efficiency. In the limit *κ*_V_ → ∞, replication efficiency approaches the replication efficiency of a single-length VCG pool containing only oligomers of length 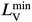 (Fig. 4C and Fig. 2G).

### Adding dinucleotides to the feedstock decreases replication efficiency

So far, we focused on ensembles that contain solely monomers as feedstock. However, examining the influence of dimers on replication in VCG pools is of interest, since dinucleotides have proven to be interesting candidates for enzyme-free RNA copying (***Walton and Szostak, 2016***; ***Sosson et al., 2019***; ***Leveau et al., 2022***). For this reason, we study oligomer pools like those illustrated in Fig. 5A: The ensemble contains monomers, dimers and VCG oligomers of a single length, *L*_V_. As our default scenario, the dimer concentration is set to 10% of the monomer concentration, corresponding to *κ*_F_ = 2.3, but this ratio can be modified by changing *κ*_F_.

**Figure 5.**
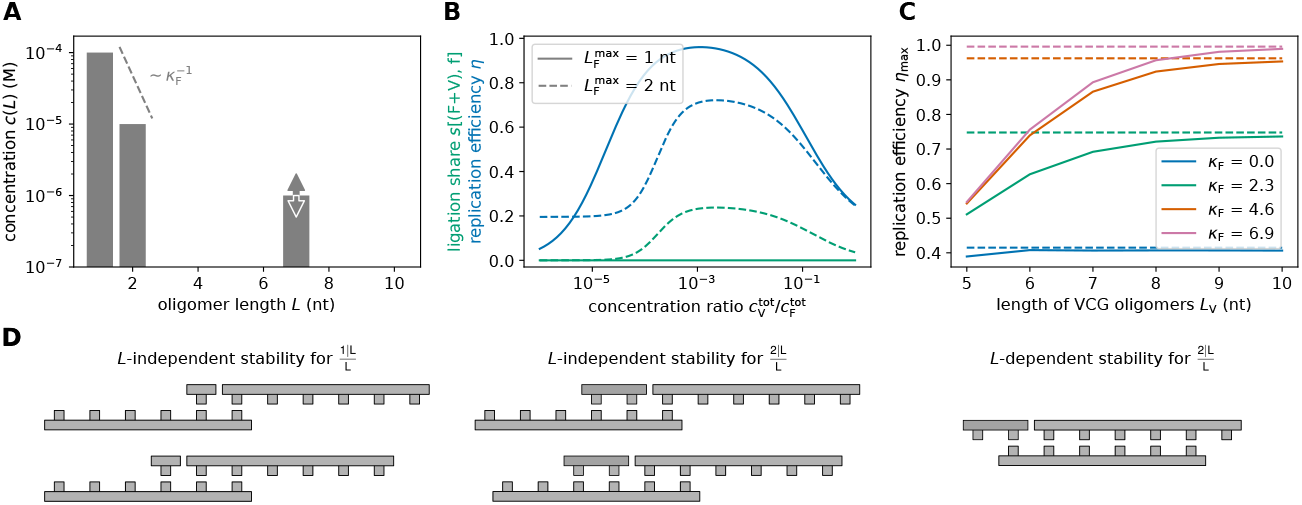
Replication performance of single-length VCG pools containing monomers and dimers as feedstock. **(A)** The pool contains a fixed total concentration of feedstock, 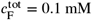, partitioned into monomers and dimers, as well as VCG oligomers of a single length, *L*_V_. The proportion of monomers and dimers can be adjusted via *κ*_F_, and the concentration of the VCG oligomers is a free parameter, 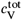. **(B)** Replication efficiency exhibits a maximum at intermediate VCG concentration in systems with (dashed blue curve) and without dimers (solid blue curve). The presence of dimers reduces replication efficiency significantly, as they enhance the ligation share of incorrect F+V ligations (dashed green curve). The panel depicts the behavior for *L*_V_ = 7 nt and *κ*_F_ = 2.3. **(C)** Optimal replication efficiency increases as a function of oligomer length, *L*_V_, and asymptotically approaches a plateau (dashed lines, Eq. (6)). The value of this plateau, 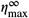, is determined by the competition between correct and false 2+V reactions, both of which grow exponentially with *L*_V_. Thus, 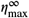 depends on the relative concentration of the dimers in the pool: the more dimers are included, the lower is 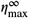. **(D)** Erroneous 1+V ligations are possible if the educt oligomer has a short overlap region with the template. The hybridization energy for such configurations is small, and independent of the length of the VCG oligomers (left). While 2+V ligations may produce incorrect products via the same mechanism (middle), they can also be caused by complexes in which two VCG oligomers hybridize perfectly to each other, but the dimer has a dangling end. The stability of these complexes increases exponentially with oligomer length (right).

Fig. 5B compares the replication efficiency of a pool with 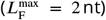 and without dimers 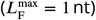. In both cases, pools that are rich in VCG oligomers exhibit low efficiency of replication. As erroneous V+V ligations are the dominant type of reaction in this limit, all pools achieve the same efficiency regardless of the presence of dimers. In contrast, when the pool is rich in feedstock (small 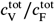), pools with and without dimers behave differently: If only monomers are included, the efficiency approaches zero, as dimerizing monomers do not contribute to the yield, and thus not to the efficiency. However, the presence of dimers enables the ligation of monomers and dimers to form trimers and tetramers, which lead to a non-zero yield. Given the low VCG concentration, the ligations are likely to proceed using dimers as template. As a consequence, educt oligomers can only hybridize to the template with a single-nucleotide-long hybridization region, leading to frequent formation of incorrect products and, consequently, low efficiency.

Replication efficiency reaches its maximum at intermediate VCG concentrations, where replication is dominated by F+V ligations. Notably, the maximal attainable efficiency is significantly lower for pools with dimers than without, as dimers increase the number and the stability of complex configurations that can form incorrect products (Fig. 5C). Without dimers, ligation products are only incorrect if the overlap between the VCG educt oligomer and template is shorter than the unique motif length, *L*_*U*_. With dimers, however, dangling end dimers can cause incorrect products even in case of long overlap of educt oligomer and template (right column in Fig. 5C). The stability of the latter complexes depends on the length of the VCG oligomers, *L*_V_, whereas the stability of complexes facilitating incorrect monomer addition is independent of oligomer length (Fig. 5C).

In the presence of dimers, the length-dependent stability of complexes allowing for correct and incorrect F+V ligations causes a competition, which sets an upper bound on the efficiency of replication (Supplementary Material Section S10),

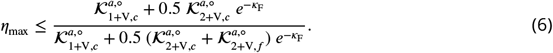

Here, we introduced effective association constants 𝒦^*a*^, which depend differently on the VCG oligomer length, *L*_V_. While the effective association constant 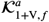 of complexes enabling incorrect 1+V ligations is length-independent, the effective association constant for incorrect 2+V ligations, 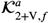, scales exponentially with *L*_V_,

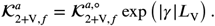

The effective association constants for correct 1+V and 2+V ligations also scale exponentially with the oligomer length (Supplementary Material Fig. S14).

In systems without dimers, i.e., 𝒦_F_ → ∞, *η*_max_ approaches 100%, which is consistent with the behavior observed in the previous sections. Conversely, in systems containing dimers, the maximal efficiency remains at a value below 100%, which depends on the concentration of dimers in the feedstock. Fig. 5C shows the dependence of maximal replication efficiency on the length of VCG oligomers in pools containing monomers, dimers and VCG oligomers. As *L*_V_ increases, *η*_max_ converges towards the upper bound defined in Eq. (6) (dashed line in Fig. 5C).

### Kinetic suppression of error-prone VCG oligomer ligation

In all scenarios considered so far, the efficiency of replication is limited by a common mechanism, regardless of the specifics of the VCG pool: Ligations involving two oligomers that hybridize to the template over a region shorter than *L*_*U*_ are prone to generate incorrect products. In previous sections, we minimized these erroneous ligations by fine-tuning the concentration and length of VCG oligomers. However, such control may become unnecessary if the typically error-prone ligation of two oligomers (V+V) is kinetically suppressed. Kinetic suppression can be an intrinsic property of the activation chemistry: Templated ligation of two oligomers can be several orders of magnitude slower than the extension of an oligomer by a single monomer (***Prywes et al., 2016***; ***Ding et al., 2021***). As a result, V+V ligations are disfavored purely by their slower kinetics. In addition, it is conceivable that only monomers are chemically activated while longer oligomers remain inactive, which would further reduce the likelihood of erroneous ligations. This scenario has already been explored experimentally (***Ding et al., 2023***). In natural environments, it could occur, for instance, when activated monomers are produced externally and then diffuse into compartments containing the VCG but lacking internal activation pathways (***Toparlak et al., 2023***; ***Kriebisch et al., 2024***).

Within our model, we capture the kinetic suppression by introducing two different rates of ligation, *k*_lig,1_ for ligations involving a monomer and *k*_lig,>1_ for ligations involving no monomer, allowing for kinetic discrimination between these processes. We explore the resulting replication efficiency in the limit of perfect kinetic discrimination (*k*_lig,>1_/*k*_lig,1_ → 0) where only monomers are reactive for ligation. We first consider a pool where the reactive monomers are mixed with VCG oligomers of a single length as well as non-reactive dimers (Fig. 6A). We vary the concentration of VCG oligomers, but keep the feedstock concentrations constant. For small VCG concentrations, we observe low efficiencies (Fig. 6B), as the ligation of two monomers, or one monomer and one dimer, are most likely. Conversely, high 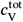 facilitates the formation of complexes in which VCG oligomers are extended by monomers, which implies high efficiency (Fig. 6B). Note that, unlike in systems where all oligomers are reactive, replication efficiency does not decrease for high VCG concentration, as erroneous V+V ligations are impossible. Instead, perfect replication efficiency (100%) is reached for sufficiently high *L*_V_.

**Figure 6.**
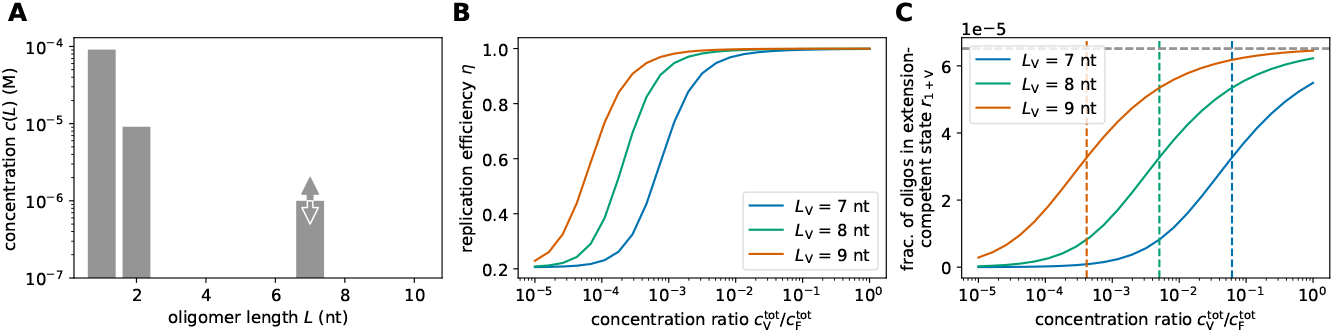
Replication performance of single-length VCG pools with kinetic suppression of ligation between oligomers. **(A)** The pool contains reactive monomers alongside non-reactive dimers and VCG oligomers of a single length. The concentrations of monomers and dimers are fixed, *c*(1) = 0.091 mM and *c*(2) = 9.1 µM, adding up to a total feedstock concentration of 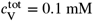, while the concentration of VCG oligomers, 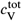, is varied. **(B)** Unlike in pools in which all ligation processes occur, replication efficiency does not decrease at high VCG concentration if ligations not involving monomers are kinetically suppressed. Instead, replication efficiency approaches an asymptotic value of 100%, as erroneous V+V ligations are impossible. **(C)** The fraction of oligomers that are in a monomer-extension-competent state depends on the total concentration of VCG oligomers. At low VCG concentration, most oligomers are single-stranded, and extension of oligomers by monomers is scarce. At high VCG concentration, *r*_1+V_ approaches the asymptotic value 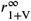 (Eq. (7), grey dashed line). In this limit, almost all oligomers form duplexes, which facilitate monomer addition upon hybridization of a monomer. Thus, the asymptotic fraction of oligomers that gets extended by monomers is not determined by the oligomer length, but by the binding affinity of monomers to existing duplexes. Conversely, the threshold concentration at which 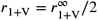 depends on oligomer length (colored dashed lines): Longer oligomers reach higher *r*_1+V_ at lower VCG concentration.

While replication efficiency characterizes the relative amount of nucleotides used for the correct elongation of VCG oligomers, it is also interesting to analyze which fraction of VCG oligomers is in a monomer-extension-competent state,

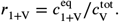

Here, 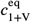 denotes the equilibrium concentration of all complexes enabling the addition of a monomer to any VCG oligomer. We find that *r*_1+V_ depends on the VCG concentration qualitatively in the same way as the efficiency: *r*_1+V_ is small for small VCG concentration, but approaches a value 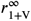 asymptotically for high VCG concentration (Fig. 6C). In this limit, almost all VCG oligomers are part of a duplex. Monomers can bind to these duplexes to form complexes allowing for 1+V ligations. Since almost all VCG oligomers are already part of a duplex, increasing 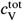 further does not increase the fraction of VCG oligomers that can be extended by monomers. Instead, the asymptotic value is determined by the concentration of monomers and their binding affinity *K*_*d*_ (1) to an existing duplex (Supplementary Material Section S11),

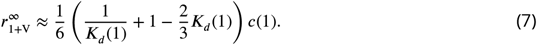

While the asymptotic value 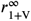 does not depend on the length of the VCG oligomers, the threshold VCG concentration at which 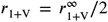 scales exponentially with *L*_V_ (Supplementary Material Section S11),

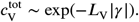

This scaling implies that longer oligomers require exponentially lower VCG concentration to achieve a given ratio *r*_1+V_ (Fig. 6C), as their greater length allows them to form more stable complexes. This observation implies that pools with longer oligomers will always be more productive than pools with shorter oligomers (at equal VCG concentration).

The behavior becomes more complex in pools containing VCG oligomers of multiple lengths, due to the competitive binding within such heterogeneous pools. To illustrate this, we examine an ensemble containing VCG oligomers ranging from 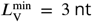 to 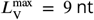, along with the same feedstock as previously (reactive monomers and non-reactive dimers, see Fig. 7A). For simplicity, we assume that the length-distribution of VCG oligomers is uniform. We study the fraction of oligomers in a monomer-extension-competent state as a function of oligomer length,

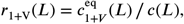

where 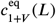 denotes the equilibrium concentration of all complexes that allow monomer-extension of VCG oligomers of length *L*. At low VCG concentration, longer oligomers are more likely to be extended by monomers than shorter ones (Fig. 7B). This behavior is intuitive, as longer oligomers tend to form more stable complexes, which lead to higher productivity. Surprisingly, increasing the VCG concentration reverses the length-dependence of the productivity, such that short oligomers are more likely to be extended by monomers than long ones (note the three crossings of the curves in Fig. 7B). For example, 8-mers are more likely to undergo primer-extension than 9-mers once the VCG concentration exceeds ≈ 0.5 µM. We derived a semi-analytical expression for the threshold VCG concentrations at which oligomers of two different lengths have equal productivity (dashed lines in Fig. 7B, Supplementary Material Sections S12 and S13).

**Figure 7.**
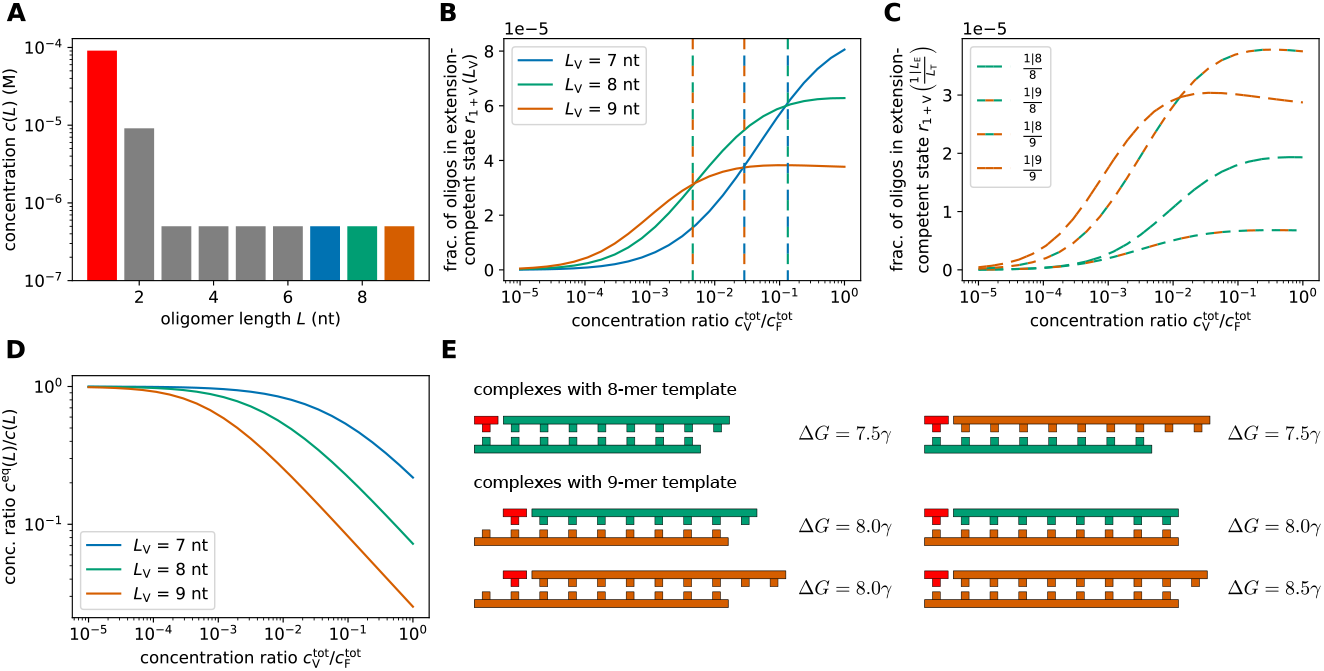
Replication performance of multi-length VCG pools with kinetic suppression of ligation between oligomers. **(A)** The pool contains reactive monomers as well as non-reactive dimers and VCG oligomers. The concentrations of monomers and dimers are fixed, *c*(1) = 0.091 mM and *c*(2) = 9.1 µM, adding up to a total feedstock concentration of 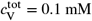, while the total concentration of VCG oligomers, 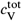, is varied. All VCG oligomers are assumed to have the same concentration. **(B)** At low VCG concentration, long oligomers are more likely in a monomer-extension-competent state than short oligomers, whereas at high VCG concentration, the trend reverses and short oligomers are more likely to be extended by monomers (“inversion of productivity”). The threshold concentration at which a short oligomer starts to outperform a longer oligomer depends on the lengths of the compared oligomers (dashed lines). **(C)** The mechanism underlying inversion of productivity can be understood based on the pair-wise competition of different VCG oligomers, e.g., 8-mers vs. 9-mers. Over the entire range of VCG concentrations, complexes with 8-mer templates have a lower relative equilibrium concentration than complexes with 9-mer templates (bottom two curves vs. top two curves). As the concentration of VCG oligomers is increased, ligations of type 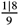 exceed ligations of type 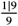, i.e., the extension fraction of 8-mers that are extended by monomers using a 9-mer as a template exceeds the fraction of extended 9-mers. **(D)** The equilibrium concentration of free oligomer decreases with increasing 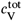. For longer oligomers, the equilibrium fraction of free oligomers is lower, as they can form more stable complexes with longer hybridization sites. **(E)** Complexes in which 8-mers serve as template are less stable than complexes with 9-mer templates, explaining why complexes with 8-mer template are more abundant than complexes with 9-mer template (see panel C). Complexes with 9-mer template have similar stability regardless of the length of the educt oligomer, i.e., 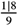 and 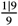 are similarly stable. This similar stability together with the higher concentration of free 8-mers compared to 9-mers (see panel D) is the reason why the fraction of monomer-extended 8-mers exceeds the one of 9-mers (see panel C).

To understand the mechanism underlying the inversion of productivity, we analyze how different complex types contribute to *r*_1+V_*(L)*. We introduce

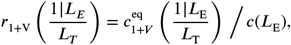

which denotes the fraction of oligomers of length *L*_E_ that are in a monomer-extension-competent complex configuration that uses an oligomer of length *L*_T_ as template. The term 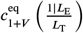 includes the sum over all possible configurations of complexes with the given lengths. Focusing on 8-mers and 9-mers as an example, we observe that complexes utilizing the 9-mer as template are responsible for the inversion of productivity (top two curves in Fig. 7C): As the VCG concentration increases, the fraction of monomer-extendable 8-mers eventually surpasses the fraction of monomer-extendable 9-mers, i.e., 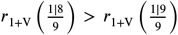 (Fig. 7C). Two factors give rise to this feature: (i) Ternary complexes of type 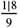 and 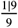 have similar stability (Fig. 7E), and (ii) the equilibrium concentration of free 8-mers is higher than that of 9-mers (Fig. 7D). As a result, 8-mers are more likely than 9-mers to hybridize to an existing duplex 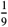, and, given the stability of the complex 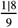, 8-mers remain bound almost as stably as 9-mers. In summary, short oligomers sequester long oligomers as templates to enhance their monomer-extension rate, while long oligomers cannot make use of short oligomers as templates due to the relative instability of the corresponding complexes.

It is noteworthy that Ding *et al*. have already observed the inversion of productivity experimentally (***Ding et al., 2023***). In their study, they included activated monomers, activated imidazolium-bridged dinucleotides and oligomers up to a length of 12 nt, and observed that the primer extension rate for short primers is higher than the extension rate of long primers. Even though our model differs from their setup in some aspects (e.g., different circular genome, no bridged dinucleotides) evaluating our model using parameters similar to those of the experimentally studied system predicts inversion of productivity that qualitatively agrees with the experimental findings (Supplementary Material Section S14). We therefore assume that the mechanism underlying inversion of productivity described here also applies to the experimental observations.

## Discussion and Conclusion

While significant progress has been made in understanding the prebiotic formation of ribonucleotides (***Powner et al., 2009***; ***Kim et al., 2011***; ***Benner et al., 2012***; ***Becker et al., 2016***) and characterizing ribozymes that might play a role in an RNA world (***Mutschler et al., 2015***; ***Attwater et al., 2018***; ***Pressman et al., 2019***; ***Tjhung et al., 2020***), a convincing scenario bridging the gap between prebiotic chemistry and ribozyme-catalyzed replication is still missing. Here, we studied a scenario proposed by Zhou *et al*. (***Zhou et al., 2021***) (the ‘Virtual Circular Genome’, VCG) using theoretical and computational approaches. We analyzed the process whereby template-directed ligation replicates the genomic information that is collectively stored in the VCG oligomers. Our analysis revealed a trade-off between the fidelity and the yield of this process: At low concentration of VCG oligomers, most of the ligations produce oligomers that are too short to specify a unique locus on the genome, resulting in a low yield of replication (Fig. 2B-C). At high VCG concentration, erroneous templated ligations cause sequence scrambling and consequently limit the fidelity of replication (Fig. 2C-D). We considered two solutions to these issues: (i) a VCG pool composition that optimizes its replication behavior within the bounds of the fidelity-yield trade-off, and (ii) breaking the fidelity-yield trade-off given that error-prone ligations can be kinetically suppressed.

The first solution maximizes the yield of replication for fixed fidelity. In pools containing only monomers and VCG oligomers of a single length, replication efficiency can be maximized by increasing the length of VCG oligomers and decreasing their concentration (Fig. 2F-G). This reduces the likelihood of error-prone templated ligation of long oligomers. When the pool contains VCG oligomers of multiple lengths, replication efficiency is typically governed by the longest oligomer in the pool (Fig. 4D). Including dimers as feedstock for the replication increases the error fraction (Fig. 5B-C), as dimers that bind to a template with a dangling end are prone to form an incorrect product (Fig. 5D).

The second solution eliminates the error-prone templated ligation of two VCG oligomers by suppressing them kinetically, e.g., by assuring that only monomers are chemically activated. This enables both fidelity and yield to remain high at high VCG concentrations (Fig. 6B), effectively breaking the fidelity-yield trade-off. Longer VCG oligomers are then more likely to be extended than shorter oligomers at equal concentration, (Fig. 6C). However, this is only true for pools with VCG oligomers of a single length — once multiple VCG oligomer lengths compete with each other, shorter oligomers can be more productive than longer ones (Fig. 7B). This feature, which has also been observed experimentally (***Ding et al., 2023***), is caused by an asymmetry in the productive interaction between short and long oligomers (Fig. 7C): While short oligomers can sequester longer oligomers as templates for their extension by a monomer, short oligomers are unlikely to serve as templates for longer oligomers (Fig. 7D-E).

As we intended to study the pathways responsible for sequence scrambling and to explore possible mitigation strategies, we based our analysis on a coarse-grained model which neglects some experimental details. First, we assumed that a complex instantaneously dehybridizes if it contains a non-complementary base pair, whereas in reality, short duplexes can tolerate a limited number of mismatches (***Todisco et al., 2024a***). While such mismatches can facilitate incorrect hybridization and introduce additional replication errors, we expect this effect to be moderate: Mismatches preferentially occur near the ends of the hybridized region, where their destabilizing effect on binding is weakest (***Todisco et al., 2024a***). However, such terminal mismatches have also been shown to significantly reduce ligation rates (***Rajamani et al., 2010***; ***Leu et al., 2013***), which in turn limits the likelihood of forming incorrect products.

Second, we simplified the hybridization dynamics by assuming that all oligomers bind to each other at equal rates, and that dehybridization rates are determined by the hybridization energy computed via a nearest-neighbor model. However, recent work has shown that hybridization to a gap flanked by two oligomers proceeds more slowly than binding to an unoccupied template. Moreover, the resulting nicked complexes (two oligomers hybridized adjacently on a template) are more stable than predicted by standard nearest-neighbor models due to enhanced stacking interactions at the nick site (***Todisco et al., 2024b***). While this added stability is not expected to affect overall replication efficiency of the VCG (since all productive complexes, correct or incorrect, contain a nick), it can impact the kinetics of the system. In particular, the extended lifetime of such complexes may challenge the adiabatic approximation used in much of our analysis, which assumes ligation is always slower than hybridization and dehybridization.

Third, we do not model the activation chemistry explicitly, but instead assume that all monomers (and, depending on the scenario, also oligomers) are always reactive. As a result, some activated intermediates that are known to form in experiments, such as imidazolium-bridged dinucleotides (***Walton and Szostak, 2016***), are not modeled. Nonetheless, we include aspects of activation chemistry in a coarse-grained manner. Specifically, to capture the experimentally observed difference in reactivity between monomer incorporation and templated ligation of oligomers under aminoimidazolium activation, we introduce two distinct ligation rate constants. With this approach, we describe the experimental setup well enough to qualitatively reproduce features observed in experiments, for example, the preferential extension of shorter oligomers by monomers in pools containing VCG oligomers of varying lengths (***Ding et al., 2023***).

The VCG scenario was proposed to close the gap between prebiotic chemistry and ribozyme-catalyzed replication. To this end, VCG pools need to be capable of replicating (parts of) ribozymes that play a role in the emergence of life. While there are cases of small ribozymes (***Pressman et al., 2019***) or ribozymes with small active sites (e.g., the Hammerhead ribozyme (***Scott et al., 2013***)), ribozymes obtained experimentally via *in vitro* evolution are often more than a hundred nucleotides long (***Johnston et al., 2001***; ***Müller and Bartel, 2008***; ***Wochner et al., 2011***; ***Attwater et al., 2013***). Remarkably, our model suggests that the VCG scenario enables high-fidelity replication of long genomes, even in pools containing relatively short VCG oligomers. For a genome of length *L*_*G*_, a sequence of at least 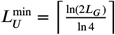 nucleotides is required to uniquely specify a position within the genome. If the oligomers in the pool exceed this length by about three nucleotides, accurate replication becomes feasible (Fig. 2H). For example, genomes of length 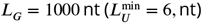 can be replicated in VCG pools containing 10 nt oligomers. However, *L*_*U*_ equals 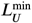 only for a restricted set of genome sequences; more generally, 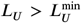. In such cases, reliable replication requires correspondingly longer oligomers. While the precise margin between oligomer length and *L*_*U*_ depends on genome-specific features (particularly the motif distribution at sub-*L*_*U*_ scales), it typically amounts to only a few additional nucleotides (Section S8 in the Supplementary Material).

It is noteworthy that replication in the VCG scenario imposes a selection pressure on prebiotic genomes to reduce their unique motif length, *L*_*U*_. A circular genome requiring many nucleotides to specify a unique locus (high *L*_*U*_) replicates less efficiently than one with a shorter *L*_*U*_, assuming all other properties of the VCG pool — particularly the oligomer length distribution — remain identical. This length distribution arises from the interplay between the chemical kinetics and molecular transport governed by the physical environment. For instance, templated ligation in an open system with continuous oligomer influx and outflux can produce a non-monotonic length distribution, with a concentration peak at a characteristic oligomer length *L*_*c*_, determined by the interplay between dehybridization and outflux (***Rosenberger et al., 2021***). Through this emergent length scale, the environment shapes replication in the VCG scenario. If the environment facilitates long oligomers (*L*_*c*_ *> L*_*U*_), replication proceeds efficiently. Conversely, in environments with a small *L*_*c*_, repeating motifs longer than *L*_*c*_ are selected against. In such cases, mutational errors may replace long repeated motifs with functionally equivalent sequences composed of shorter unique motifs, thereby increasing replication efficiency.

Given the broad range of prebiotically plausible non-equilibrium environments (***Ianeselli et al., 2023***), it is reasonable to expect that some environments provide the required conditions for efficient replication. The constraints formulated in this work can help to guide the search for self-replicating oligomer pools, in the vast space of possible concentration profiles and non-equilibrium environments.

## Acknowledgments

We thank Paul Higgs and members of the Gerland group for stimulating discussions. This work was supported by the Deutsche Forschungsgemeinschaft (DFG, German Research Foundation) via the CRC/TRR 392 Molecular Evolution (Project-ID 521256690), and under Germany’s Excellence Strategy (EXC-2094-390783311, ORIGINS).

## Code Availability

Most of the software used in this study is publicly available at https://github.com/gerland-group/VirtualCircularGenome. This repository includes code for sampling circular genomes using a Metropolis-Hastings-type algorithm, as well as code for the coarse-grained adiabatic approach used in the numerical analysis of replication performance. The full kinetic simulation code is available from the authors upon reasonable request.

## Supplementary Material

### S1. CONSTRUCTING CIRCULAR GENOMES

In the Virtual Circular Genome (VCG) scenario, genomes are encoded in a pool of oligomers. The encoded genomes are assumed to be circular sequences of length *L*_*G*_, containing both the original sequence and its reverse complement. Each genome is characterized by two fundamental length scales that reflect different aspects of motif distribution along the sequence. The minimal unique motif length, *L*_*U*_, is defined as the shortest subsequence length for which all motifs of length *L* ≥ *L*_*U*_ appear at most once in the genome. In contrast, the exhaustive coverage length, *L*_*E*_, denotes the largest motif length for which all 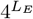 possible motifs are present within the genome. Since only 2*L*_*G*_ distinct motifs can be encoded in a genome (including its complement), it follows that *L*_*E*_ cannot exceed

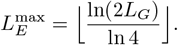

Similarly, for a motif to be uniquely addressable on the genome, its length must be at least

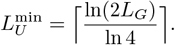

We note that 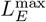 and 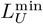 are essentially the same length scale (differing by at most one nucleotide), such that they can be summarized into a single characteristic subsequence lengthscale *L*_*S*_ for convenience.

The characteristic length scales *L*_*E*_ and *L*_*U*_ impose constraints on how motifs are distributed. For example, when *L* = *L*_*U*_, all motifs of length *L* must appear at most once, while at least one motif of length *L* − 1 must occur more than once. To quantify motif distributions, we introduce the motif entropy,

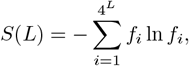

where *f*_*i*_ denotes the relative frequency of motif *i* across the genome and its reverse complement. Motif entropy ranges from zero (a homogeneous sequence with only one motif) to a maximum value that depends on the subsequence length,

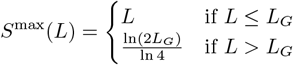

For a motif length *L* to qualify as the unique motif length *L*_*U*_, its entropy must be maximal, *S*(*L*) = *S*^max^(*L*), while *S*(*L* − 1) must be smaller than its respective maximum, *S*(*L* − 1) *< S*^max^(*L* − 1).

The correspondence between characteristic length scales and motif entropy provides a way to construct genome sequences with specified motif characteristics. By treating the entropy function as an effective “Hamiltonian” ℋ, we can generate genome sequences through Metropolis–Hastings sampling. Each update step in the Metropolis-Hastings algorithm involves either a single-nucleotide mutation or a cut-and-paste operation that relocates a segment of the genome to a new position (the cut-and-paste operation is 10 times more likely than the single nucleotide mutation). The acceptance criterion follows the standard Metropolis rule: modifications that reduce the Hamiltonian are always accepted, while increases in energy are accepted with probability exp [−*β*(*E*_old_ − *E*_new_)], where *β*^−1^ is an effective temperature chosen to be small compared to the typical energy to ensure convergence to the minimum. Simulations are either run until a predefined entropy target is reached or until the energy converges to a plateau.

To generate genomes with 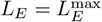 and 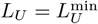, we minimize the Hamiltonian

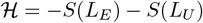

Starting from a random sequence, we perform 10,000 Metropolis–Hastings steps at an inverse temperature *β* = 10^−5^ to construct genome sequences of lengths *L*_*G*_ = 16 nt (Main Text) and *L*_*G*_ = 64 nt (Table S2).

To explore genomes, where 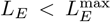 and 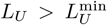, we initialize the algorithm from the maximally diverse genomes 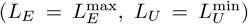 and then reduce entropy across the range *L*_*E*_ *< L < L*_*U*_. This is done by minimizing the Hamiltonian

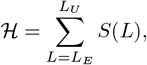

using two different sampling protocols. In the first, the simulation is terminated as soon as the genome reaches the desired values of *L*_*E*_ and *L*_*U*_. The resulting motif distributions on the intermediate length scales (*L*_*E*_ *< L < L*_*U*_) remain close to uniform, with only minor biases sufficient to enforce the length-scale constraints. In the second protocol, entropy minimization continues beyond the point at which the target values are achieved, leading to more strongly biased motif distributions on intermediate length scales. These construction strategies allow us to systematically tune genome complexity and motif structure, enabling controlled investigations of how the characteristic length scales influence replication dynamics (Section S8).

### S2. COMPUTING REPLICATION OBSERVABLES BASED ON THE KINETIC SIMULATION

We simulate the dynamics of VCG pools using a kinetic simulation that is based on the Gillespie algorithm. In the simulation, oligomers can hybridize to each other to form complexes, or dehybridize from an existing complex. Moreover, two oligomers can undergo templated ligation if they are hybridized adjacent to each other on a third oligomer. At each time *t*, the state of the system is determined by a list of all single-stranded oligomers and complexes as well as their respective copy number. We refer to the state of the system at the time *t* as the ensemble of compounds ℰ_*t*_. Given the copy numbers, the rates *r*_*i*_ of all possible chemical reactions *i* ∈ ℐ can be computed. To evolve the system in time, we need to perform two steps: (i) We sample the waiting time until the next reaction, *τ*, from an exponential distribution with mean 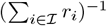, and update the simulation time, *t* → *t* + *τ*. (ii) We pick which reaction to perform by sampling from a categorical distribution. Here, the probability to pick reaction *i* equals 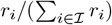. The copy numbers are updated according to the sampled reaction, yielding ℰ_*t*+*τ*_. Steps (i) and (ii) are repeated until the simulation time *t* reaches the desired final time, *t*_final_. A more detailed explanation of the kinetic simulation is presented in [12, 33].

Our goal is to compute observables characterizing replication in the VCG scenario based on the full kinetic simulation. For clarity, we focus on one particular observable (yield) for the derivation. The results for other observables are stated directly, as their derivations follow analogously. Recall the definition of the yield introduced in the main text,

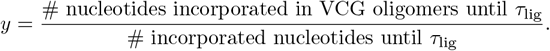

As we are interested in the initial replication performance of the VCG, we compute the yield based on the ligation events that take place until the characteristic timescale of ligations 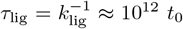. We would like to compute the yield based on the templated ligation events that we observe in the simulation. Unfortunately, for reasonable system parameters, it is impossible to simulate the system long enough to observe sufficiently many ligation events to compute *y* to reasonable accuracy. For example, for a VCG pool containing monomers at a total concentration of 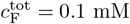 and VCG oligomers of length *L* = 8 at a total concentration of 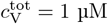, it would take about 1700 hours of simulation time to reach *t* = 5 *·* 10^12^ *t*_0_. Multiple such runs would be needed to estimate the mean and the variance of the observables of interest, rendering this approach unfeasible.

Instead, we compute the replication observables based on the copy number of complexes that could potentially perform a templated ligation, i.e., complexes, in which two strands are hybridized adjacent to each other, such that they could form a covalent bond. It can be shown analytically that the number of potentially productive complexes is a good approximation for the number of incorporated nucleotides. The number of incorporated nucleotides can be computed as the integral over the ligation flux, weighted by the number of nucleotides that are added in each templated ligation reaction,

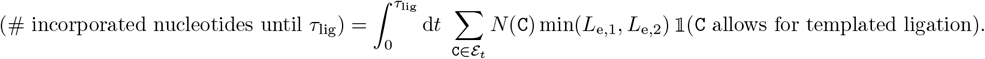

Here, *N*(`C`) denotes the copy number of the complex `C` in the pool ℰ_*t*_. *L*_e,1_ and *L*_e,2_ denote the lengths of the oligomers that undergo ligation, and 𝟙 is an indicator function which enforces that only complexes in a configuration that allows for templated ligation contribute to the reaction flux. As only few ligation events are expected to happen until *τ*_lig_, it is reasonable to assume that the ensembles ℰ_*t*_ do not change significantly during *t* ∈ [0, *τ*_lig_]. Therefore, the integration over time may be interpreted as a multiplication by *τ*_lig_,

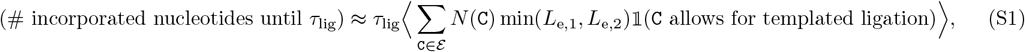

where ⟨… ⟩ denotes the average over realizations of the ensembles ℰ_*t*_ within the time interval *t* ∈ [*τ*_eq_, *τ*_lig_]. Note that, at this point, we made the additional assumption that no templated ligations are taking place between [0, *τ*_eq_]. This assumption is reasonable, as (i) the equilibration process is very short compared to the characteristic timescale of ligation, and (ii) the number of complexes that might allow for templated ligation during equilibration is lower than in the equilibrium (we start the simulation with an ensemble of single-stranded oligomers), implying that the rate of templated ligation is small.

In order to compute the average over different realizations of ensembles ℰ, we need to sample a set of uncorrelated ensembles that have reached the hybridization equilibrium. For this purpose, we run a full kinetic simulation. The simulation starts with a pool containing only single-stranded oligomers, and reaches the (de)hybridization equilibrium after a time *τ*_eq_. We identify this timescale of equilibration by fitting an exponential function to the total hybridization energy of all complexes in the system, Δ*G*_tot_ (Fig. S1A). In the set of ensembles used to evaluate the average in Eq. (S1), we only include ensembles for time *t > τ*_eq_ to ensure that the ensembles have reached (de)hybridization equilibrium. To ensure that the ensembles are uncorrelated, we require that the time between two ensembles that contribute to the average is at least *τ*_corr_. The correlation time, *τ*_corr_, is determined via an exponential fit to the autocorrelation function of Δ*G*_tot_ (Fig. S1B). Besides computing the expectation value (Eq. S1), we are also interested in the “uncertainty” of this expectation value, i.e., in the standard deviation of the sample mean *σ*_⟨*X*⟩_. (We use *X* as a short-hand notation for ∑_C∈*ℰ*_ *N*(`C`) min(*L*_e,1_, *L*_e,2_) 𝟙 (`C` allows for templated ligation).) The standard deviation of the sample mean, *σ*_⟨*X*⟩_, is related to the standard deviation of *X, σ*_*X*_, by the number of samples, *σ*_⟨*X*⟩_ = (*N*_*s*_)^−1*/*2^*σ*_*X*_. Moreover, based on the van-Kampen system size expansion, we expect the standard deviation of *X* to be proportional to *V* ^−1*/*2^. Thus, *σ*_⟨*X*⟩_ ∝ (*N*_*s*_*V*)^−1*/*2^.

Using Eq. (S1) (as well as an analogous expression for the number of nucleotides that are incorporated in VCG oligomers), the yield can be expressed as

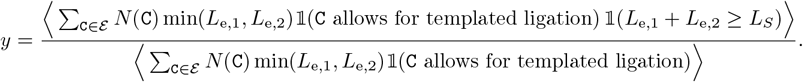

The additional condition 𝟙 (*L*_e,1_ + *L*_e,2_ ≥ *L*_*S*_) in the numerator ensures that the product oligomer is long enough to be counted as a VCG oligomer, i.e., at least *L*_*S*_ nucleotides long. Analogously, the expression for the fidelity of replication reads

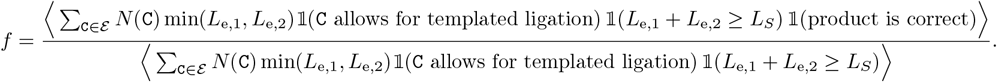

Multiplying fidelity and yield results in the efficiency of replication,

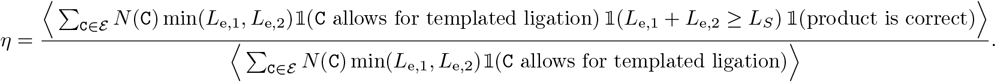

**FIG. S1.**
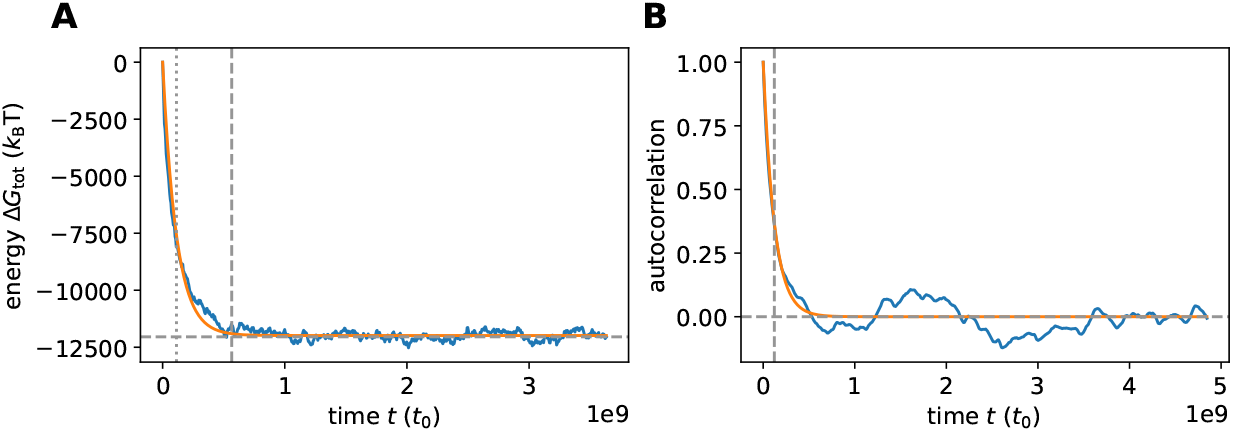
Characteristic timescales in the kinetic simulation. **A** The equilibration timescale is determined based on the total hybridization energy of all strands in the pool, Δ*G*_tot_. By fitting an exponential function to Δ*G*_tot_, we obtain a characteristic timescale, *τ* ^*\**^ (vertical dotted line), which is then used to calculate the equilibration time as *τ*_eq_ = 5*τ* ^*\**^ (vertical dashed line). The horizontal dashed line shows the total hybridization energy expected in (de)hybridization equilibrium according to the coarse-grained adiabatic approach (Sections S3 and S4). **B** The correlation timescale is determined based on the autocorrelation of Δ*G*_tot_. We obtain *τ*_corr_ (vertical dashed line) by fitting an exponential function to the autocorrelation. In both panels, we show simulation data obtained for a VCG pool containing monomers and VCG oligomers with a concentration of 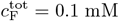 as well as oligomers of length *L* = 8 nt with a concentration of 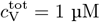.

The ligation share of a particular type of templated ligation *s*(type), that is, the relative contribution of this templated-ligation type to the nucleotide extension flux, can be represented in a similar form as the other observables,

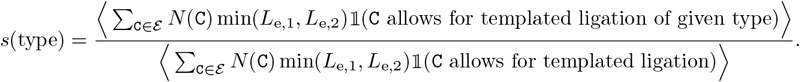

As all observables are expressed as the ratio of two expectation values, 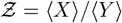, we can compute the uncertainty of the observables via Gaussian error propagation,

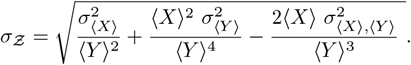

Since the variances, 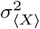 and 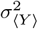, as well as the covariance, 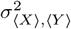, are proportional to (*N*_*s*_*V*)^−1^, the standard deviation of the observable mean, *σ*_*Z*_, scales with the inverse square root of the number of samples and the system volume, i.e., *σ*_*Z*_ ∝ (*N*_*s*_*V*)^−1*/*2^. Therefore, the variance of the computed observable can be reduced by either increasing the system volume or increasing the number of samples used for averaging. Both approaches incur the same computational cost: (i) Increasing the number of samples, *N*_*s*_, requires running the simulation for a longer duration, with the additional runtime scaling linearly with the number of samples. (ii) Similarly, the additional runtime needed due to increased system volume, *V*, also scales linearly with *V* (Fig. S2). One update step in the simulation always takes roughly the same amount of runtime, but the change in simulation time per update step depends on the total rate of all reactions in the system. The total rate is dominated by the association reactions, and their rate is proportional to the volume. Therefore, the change in simulation time per update step is proportional to *V* ^−1^. The runtime, which is necessary to reach the same simulation time in a system with volume *V* as in a system with volume 1, is a factor of *V* longer in the larger system. With this in mind, it makes no difference whether the variance is reduced by increasing the volume or the number of samples. For practical reasons (post-processing of the simulations is then less memory- and time-consuming), we opt to choose moderate number of samples, but slightly higher system volumes to compute the observables of interest. The simulation parameters (length of oligomers, concentrations, hybridization energy, volume, number of samples, characteristic timescales) used to obtain the results presented in the main text (Main Text Fig. 2**Replication performance of VCG pools containing VCG oligomers of a single length (single-length VCG pools). (A)** The pool contains a fixed concentration of monomers, 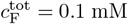, as well as VCG oligomers of a single length, *L*_V_, at variable concentration 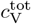 (the VCG oligomers cover all possible subsequences of the genome and its complement at equal concentration). **(B)** The yield increases as a function of 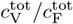, because dimerizations become increasingly unlikely for high VCG concentrations. **(C)** The ligation share of different ligation types depends on the total VCG concentration: In the low concentration limit, dimerization (F+F) dominates; for intermediate concentrations, F+V ligations reach their maximum, while, for high concentrations, a substantial fraction of reactions are V+V ligations. The panel depicts the behavior for *L*_V_ = 6 nt. **(D)** Replication efficiency is limited by the small yield for small 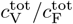. In the limit of high 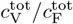, replication efficiency decreases due to the growing number of error-prone V+V ligations. Maximal replication efficiency is reached at intermediate VCG concentration. **(E)** V+V ligations are prone to the formation of incorrect products due to the short overlap between educt strand and template. In general, the probability of correct product formation, *p*_corr_, depends on the choice of circular genome and as well as its mapping to the VCG pool. The probabilities listed here refer to a VCG pool with *L*_*G*_ = 16 nt, *L*_*E*_ = 2 nt and *L*_*U*_ = 3 nt. **(F)** The optimal equilibrium concentration ratio of free VCG strands to free feedstock strands, which maximizes replication efficiency, decays as a function of length (continuous line). The analytical scaling law (dashed line, Eq. (4equation.8)) captures this behavior. The window of close-to-optimal replication, within which efficiency deviates no more than 1% from its optimum (shaded areas), increases with *L*_V_, facilitating reliable replication without fine-tuning to match the optimal concentration ratio. **(G)** Maximal replication efficiency, which is attained at the optimal VCG concentration depicted in panel E, increases as a function of *L*_V_ and approaches a plateau of 100%. For high efficiency, Eq. (5equation.9) provides a good approximation of the length-dependence of *η*_max_ (dashed lines). The oligomer length at which replication efficiency equals 95% is determined using Eq. (5equation.9) (vertical dotted lines). **(H)** The unique motif length, 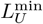, increases logarithmically with the length of the genome, *L*_*G*_. The length of VCG oligomers, *L*_V_, at which the optimal replication efficiency reaches 95% (computed using Eq. (5equation.9)) exhibits the same logarithmic dependence on *L*_*G*_.figure.caption.7) are summarized in Tab. S1.

**FIG. S2.**
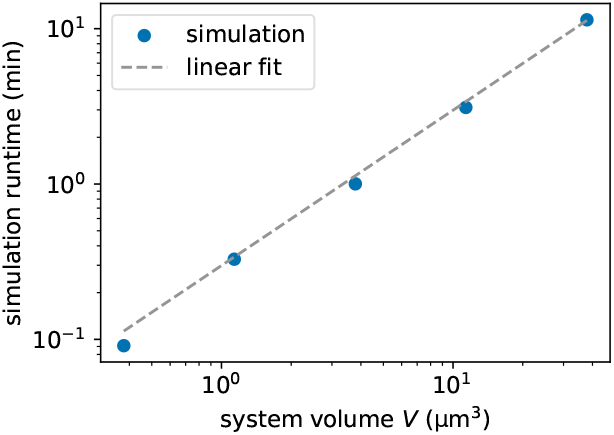
Simulation runtime of the full kinetic simulation for a VCG pool that includes monomers and VCG oligomers of length *L* = 8. The total concentration of feedstock monomers equals 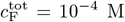, while the total concentration of VCG oligomers equals 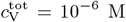. The energy contribution per matching nearest-neighbor block equals *γ* = −2.5 *k*_B_T. The volume of the system is varied, and the time-evolution is simulated until *t* = 5.0 · 10^7^*t*_0_. The runtime of the simulation scales linearly with the volume of the system.

**TABLE S1.**
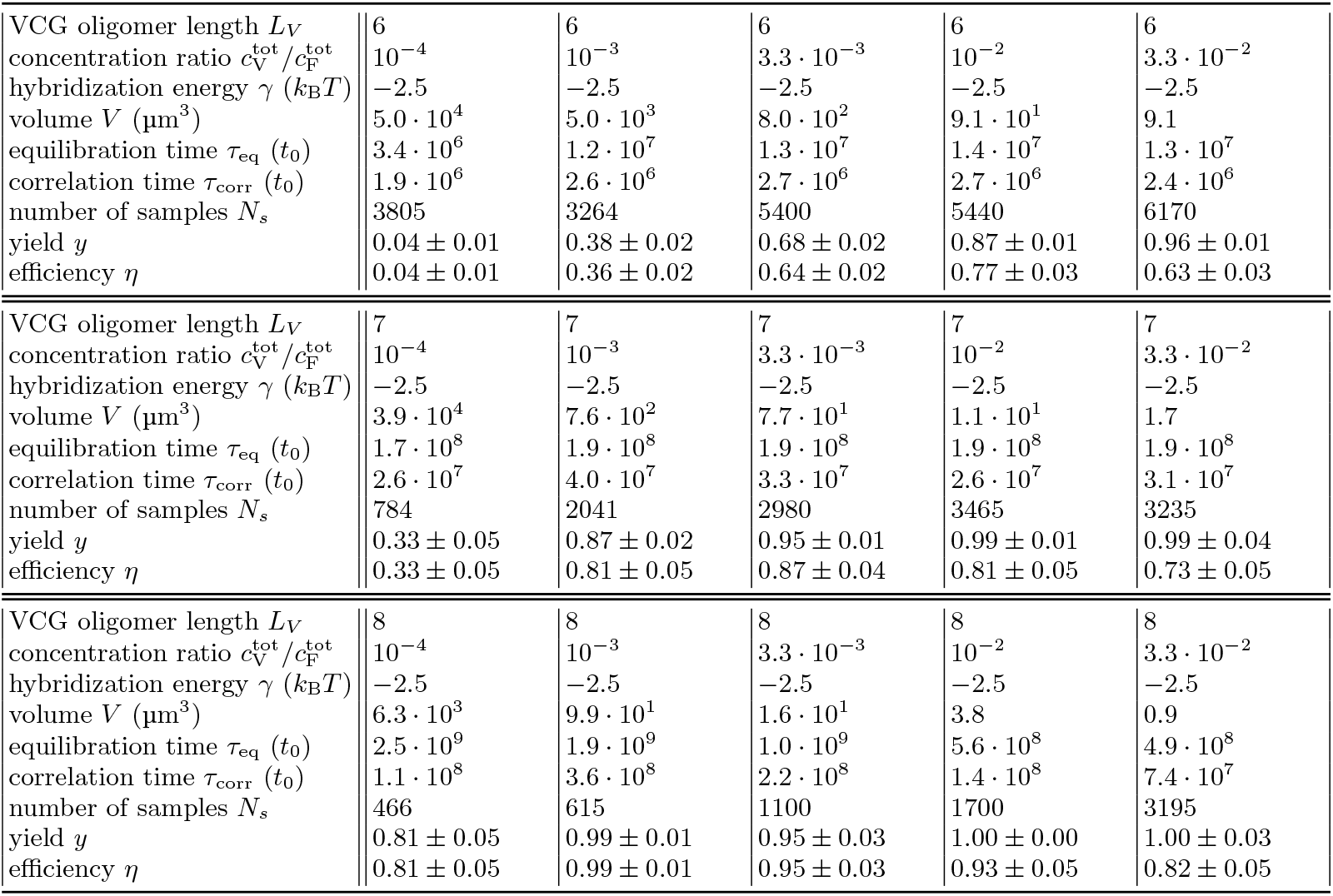
System parameters used to compute the replication observables yield, *y*, and replication efficiency, *η*, based on the kinetic simulation. The computed observables are shown in Fig. 2**Replication performance of VCG pools containing VCG oligomers of a single length (single-length VCG pools). (A)** The pool contains a fixed concentration of monomers, 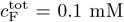, as well as VCG oligomers of a single length, *L*_V_, at variable concentration 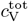(the VCG oligomers cover all possible subsequences of the genome and its complement at equal concentration). **(B)** The yield increases as a function of 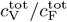, because dimerizations become increasingly unlikely for high VCG concentrations. **(C)** The ligation share of different ligation types depends on the total VCG concentration: In the low concentration limit, dimerization (F+F) dominates; for intermediate concentrations, F+V ligations reach their maximum, while, for high concentrations, a substantial fraction of reactions are V+V ligations. The panel depicts the behavior for *L*_V_ = 6 nt. **(D)** Replication efficiency is limited by the small yield for small 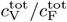. In the limit of high 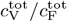, replication efficiency decreases due to the growing number of error-prone V+V ligations. Maximal replication efficiency is reached at intermediate VCG concentration. **(E)** V+V ligations are prone to the formation of incorrect products due to the short overlap between educt strand and template. In general, the probability of correct product formation, *p*_corr_, depends on the choice of circular genome and as well as its mapping to the VCG pool. The probabilities listed here refer to a VCG pool with *L*_*G*_ = 16 nt, *L*_*E*_ = 2 nt and *L*_*U*_ = 3 nt. **(F)** The optimal equilibrium concentration ratio of free VCG strands to free feedstock strands, which maximizes replication efficiency, decays as a function of length (continuous line). The analytical scaling law (dashed line, Eq. (4equation.8)) captures this behavior. The window of close-to-optimal replication, within which efficiency deviates no more than 1% from its optimum (shaded areas), increases with *L*_V_, facilitating reliable replication without fine-tuning to match the optimal concentration ratio. **(G)** Maximal replication efficiency, which is attained at the optimal VCG concentration depicted in panel E, increases as a function of *L*_V_ and approaches a plateau of 100%. For high efficiency, Eq. (5equation.9) provides a good approximation of the length-dependence of *η*_max_ (dashed lines). The oligomer length at which replication efficiency equals 95% is determined using Eq. (5equation.9) (vertical dotted lines). **(H)** The unique motif length, 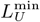, increases logarithmically with the length of the genome, *L*_*G*_. The length of VCG oligomers, *L*_V_, at which the optimal replication efficiency reaches 95% (computed using Eq. (5equation.9)) exhibits the same logarithmic dependence on *L*_*G*_.figure.caption.7 in the main text.

|  |  |  |  |  |  |
| --- | --- | --- | --- | --- | --- |
| VCG oligomer length $L_V$ | 6 | 6 | 6 | 6 | 6 |
| concentration ratio $c_V^{\text{tot}}/c_F^{\text{tot}}$ | $10^{-4}$ | $10^{-3}$ | $3.3 \cdot 10^{-3}$ | $10^{-2}$ | $3.3 \cdot 10^{-2}$ |
| hybridization energy $\gamma$ ($k_B T$) | -2.5 | -2.5 | -2.5 | -2.5 | -2.5 |
| volume $V$ ($\mu\text{m}^3$) | $5.0 \cdot 10^4$ | $5.0 \cdot 10^3$ | $8.0 \cdot 10^2$ | $9.1 \cdot 10^1$ | 9.1 |
| equilibration time $\tau_{\text{eq}}(t_0)$ | $3.4 \cdot 10^6$ | $1.2 \cdot 10^7$ | $1.3 \cdot 10^7$ | $1.4 \cdot 10^7$ | $1.3 \cdot 10^7$ |
| correlation time $\tau_{\text{corr}}(t_0)$ | $1.9 \cdot 10^6$ | $2.6 \cdot 10^6$ | $2.7 \cdot 10^6$ | $2.7 \cdot 10^6$ | $2.4 \cdot 10^6$ |
| number of samples $N_s$ | 3805 | 3264 | 5400 | 5440 | 6170 |
| yield $y$ | $0.04 \pm 0.01$ | $0.38 \pm 0.02$ | $0.68 \pm 0.02$ | $0.87 \pm 0.01$ | $0.96 \pm 0.01$ |
| efficiency $\eta$ | $0.04 \pm 0.01$ | $0.36 \pm 0.02$ | $0.64 \pm 0.02$ | $0.77 \pm 0.03$ | $0.63 \pm 0.03$ |
| VCG oligomer length $L_V$ | 7 | 7 | 7 | 7 | 7 |
| concentration ratio $c_V^{\text{tot}}/c_F^{\text{tot}}$ | $10^{-4}$ | $10^{-3}$ | $3.3 \cdot 10^{-3}$ | $10^{-2}$ | $3.3 \cdot 10^{-2}$ |
| hybridization energy $\gamma$ ($k_B T$) | -2.5 | -2.5 | -2.5 | -2.5 | -2.5 |
| volume $V$ ($\mu\text{m}^3$) | $3.9 \cdot 10^4$ | $7.6 \cdot 10^2$ | $7.7 \cdot 10^1$ | $1.1 \cdot 10^1$ | 1.7 |
| equilibration time $\tau_{\text{eq}}(t_0)$ | $1.7 \cdot 10^8$ | $1.9 \cdot 10^8$ | $1.9 \cdot 10^8$ | $1.9 \cdot 10^8$ | $1.9 \cdot 10^8$ |
| correlation time $\tau_{\text{corr}}(t_0)$ | $2.6 \cdot 10^7$ | $4.0 \cdot 10^7$ | $3.3 \cdot 10^7$ | $2.6 \cdot 10^7$ | $3.1 \cdot 10^7$ |
| number of samples $N_s$ | 784 | 2041 | 2980 | 3465 | 3235 |
| yield $y$ | $0.33 \pm 0.05$ | $0.87 \pm 0.02$ | $0.95 \pm 0.01$ | $0.99 \pm 0.01$ | $0.99 \pm 0.04$ |
| efficiency $\eta$ | $0.33 \pm 0.05$ | $0.81 \pm 0.05$ | $0.87 \pm 0.04$ | $0.81 \pm 0.05$ | $0.73 \pm 0.05$ |

TABLE S1. System parameters used to compute the replication observables yield, $y$ , and replication efficiency, $\eta$ , based on the kinetic simulation.
|  |  |  |  |  |  |
| --- | --- | --- | --- | --- | --- |
| VCG oligomer length $L_V$ | 8 | 8 | 8 | 8 | 8 |
| concentration ratio $c_V^{\text{tot}}/c_F^{\text{tot}}$ | $10^{-4}$ | $10^{-3}$ | $3.3 \cdot 10^{-3}$ | $10^{-2}$ | $3.3 \cdot 10^{-2}$ |
| hybridization energy $\gamma$ ($k_B T$) | -2.5 | -2.5 | -2.5 | -2.5 | -2.5 |
| volume $V$ ($\mu\text{m}^3$) | $6.3 \cdot 10^3$ | $9.9 \cdot 10^1$ | $1.6 \cdot 10^1$ | 3.8 | 0.9 |
| equilibration time $\tau_{\text{eq}}(t_0)$ | $2.5 \cdot 10^9$ | $1.9 \cdot 10^9$ | $1.0 \cdot 10^9$ | $5.6 \cdot 10^8$ | $4.9 \cdot 10^8$ |
| correlation time $\tau_{\text{corr}}(t_0)$ | $1.1 \cdot 10^8$ | $3.6 \cdot 10^8$ | $2.2 \cdot 10^8$ | $1.4 \cdot 10^8$ | $7.4 \cdot 10^7$ |
| number of samples $N_s$ | 466 | 615 | 1100 | 1700 | 3195 |
| yield $y$ | $0.81 \pm 0.05$ | $0.99 \pm 0.01$ | $0.95 \pm 0.03$ | $1.00 \pm 0.00$ | $1.00 \pm 0.03$ |
| efficiency $\eta$ | $0.81 \pm 0.05$ | $0.99 \pm 0.01$ | $0.95 \pm 0.03$ | $0.93 \pm 0.05$ | $0.82 \pm 0.05$ |

### S3. COARSE-GRAINED REPRESENTATION OF COMPLEXES IN THE ADIABATIC APPROACH

It is computationally expensive to evaluate the replication observables via the full kinetic simulation. For this reason, we develop an adiabatic approach, which allows us to compute the replication observables, provided that templated ligation is far slower than (de)hybridization. The adiabatic approach relies on a coarse-grained representation of the oligomers in the pool, which is introduced in this section.

#### Single Strands

In the coarse-grained description, oligomers of identical length are assumed to have equal concentration, irrespective of their sequence. This assumption is justified for two reasons: (i) We initialize the VCG pool without sequence bias, i.e., all oligomers compatible with the genome sequence are included at equal concentration. (ii) Hybridization energy in our simplified energy model (and therefore also the stability of complexes) only depends on the length of the hybridization site, not on its sequence, provided there is no mismatch. Each coarse-grained oligomer is uniquely identified by its length *L*, and it represents a group of oligomers with ℭ (*L*) distinct sequences. We refer to the number ℭ (*L*) as the combinatorial multiplicity of the coarse-grained oligomer. The value of ℭ (*L*) depends on the choice of the encoded genome. By construction (see main text), we assume that all possible oligomer sequences of length *L < L*_*S*_ are included in the genome. For *L* ≥ *L*_*S*_, only a subset of all possible 4^*L*^ sequences is included, but no sequence is repeated multiple times across the genome. Therefore, the combinatorial multiplicity equals

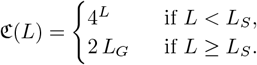

#### Duplexes

**FIG. S3.**
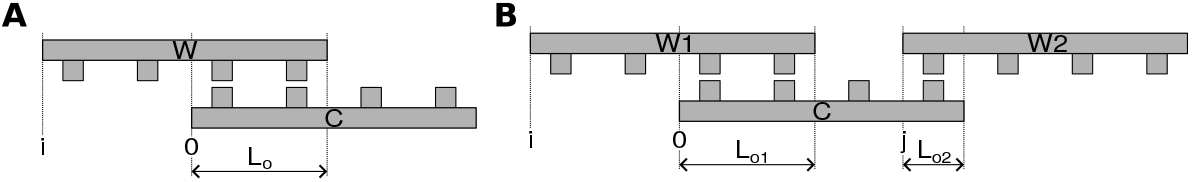
Schematic representation of complexes considered in the adiabatic approach. **A** A duplex is comprised of two strands, which we refer to as `W` (Watson) and `C` (Crick). The relative position of the strands is characterized by alignment index *i*; for the depicted duplex, *i* = −2. The length of the hybridization region is called *L*_o_. **B** A triplex contains three strands. By convention, we denote the two strands that are on the same “side” of the complex as `W1` and `W2`, and the complementary strand as `C`. The alignment indices *i* and *j* denote the positions of `W1` and `W2` relative to `C`. For the depicted triplex, *i* = −2 and *j* = 3. The length of the hybridization regions are called *L*_o,1_ and *L*_o,2_.

Two strands can form a duplex by hybridizing to each other. We refer to the bottom oligomer as ‘Crick” strand `C` and to the top oligomer as “Watson” strand `W`. A duplex is uniquely characterized by the lengths of the oligomers, *L*_C_ and *L*_W_, as well as their relative alignment (Fig. S3A). The alignment index *i* denotes the position of the Watson strand with respect to the Crick strand. As there needs to be at least one nucleotide of overlap between the strands for a duplex to exist, the alignment index needs lie in the interval *i* ∈ [− (*L*_W_ − 1), *L*_C_ − 1]. Using the alignment index, we can also determine if the duplex has a left (or right) dangling end. The corresponding indicator variables are called *d*_*l*_ (or *d*_*r*_),

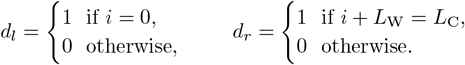

Moreover, the length of the hybridization region *L*_o_ can be computed via

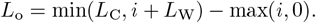

The hybridization energy of the duplex depends on the length of the hybridization region as well as on the existence/absence of dangling ends. For a hybridization site of length *L*_o_, there are *L*_o_ − 1 nearest-neighbor energy blocks each of which contributes *γ* to the energy. Moreover, each dangling end contributes 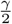 to the energy,

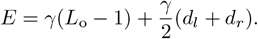

To compute the combinatorial multiplicity for a duplex with fixed *L*_C_, *L*_W_ and alignment index *i*, we need to multiply the combinatorial multiplicity of the Crick strand by the number of possible hybridization partners. We assume that a hybridization partner is possible if its sequence is perfectly complementary to the lower strand within the hybridization region, whereas hybridization partners with mismatches are not accounted for. This is sensible as long as the energetic penalty for mismatches in the full kinetic simulation is sufficiently large to suppress mismatches. The number of possible hybridization partners is determined by the length of the overlap region *L*_o_: If *L*_o_ ≥ *L*_*S*_, the pool contains only one oligomer sequence that can act as hybridization partner by construction of the genome. For shorter hybridization regions, multiple hybridization partners might be possible. Their number is set by the combinatorial multiplicity of the Watson oligomer divided by the combinatorial multiplicity of the hybridization region,

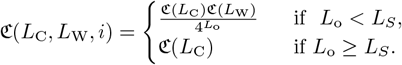

To avoid double-counting, we only account for complexes in which the Crick strand is at least as long as the Watson strand, *L*_C_ ≥ *L*_W_, and multiply ℭ (*L*_W_, *L*_C_, *i*) by 1*/*2 if *L*_W_ = *L*_C_.

#### Ternary Complexes

Ternary complexes, i.e., complexes comprised of three strands, are uniquely characterized by the length of the three oligomers, *L*_C_, *L*_W,1_, *L*_W,2_, as well as their respective alignment (Fig. S3B). The alignment index *i* denotes the position of strand `W1` relative to strand `C`. Analogously, *j* denotes the relative position of `W2` relative to oligomer `C`. Two strands that are hybridized to each other need to have a hybridization region of at least one nucleotide. Moreover, the strands `W1` and `W2` must not occupy the same position on the template strand `C`. Taking both requirements together, the alignment indices fall within the intervals,

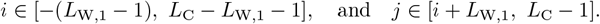

A triplex may have a dangling end not only on its left or right end, but also in the gap between strands `W1` and `W2`. Three boolean variables are necessary to denote the presence/absence of the respective dangling ends,

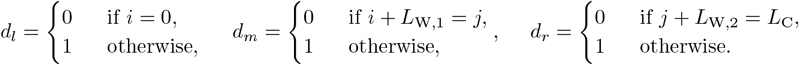

The length of the two hybridization regions are given by

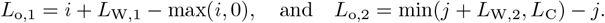

The hybridization energy depends on the length of the overlap regions as well as on the existence of dangling ends: As in the duplex, each overlap region of length *L*_o,1_ (or *L*_o,2_) comprises *L*_o,1_ − 1 (or *L*_o,2_ − 1) nearest neighbor blocks, each of which contributes *γ* to the total energy. Moreover, every dangling end contributes *γ/*2. Note that the presence of a gap between strands `W1` and `W2`, i.e., *d*_*m*_ = 1, implies that there are two dangling ends, one for `W1` and another for `W2`. Gaps in between two complexes contribute *γ/*2 per each dangling end, adding up to *γ*. If there is no gap between the strands, i.e., *d*_*m*_ = 0, there are no dangling end contributions, but a new full nearest neighbor block emerges, which contributes *γ* to the energy. Therefore, the total energy reads,

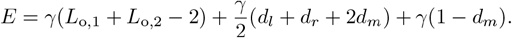

The combinatorial multiplicity of a triplex is computed in the same way as for the duplex: The combinatorial multiplicity of the strand `C` is multiplied by the number of possible hybridization partners `W1` and `W2`. Again, the number of possible partners is set by the length of the hybridization regions,

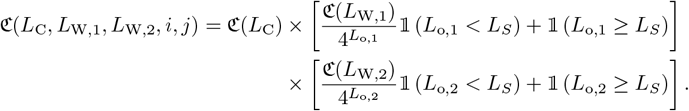

We use 𝟙 to denote the indicator function which returns 1 in case the condition in the bracket is fulfilled and zero otherwise. As all ternary complexes are asymmetric, there is no need to introduce a symmetry correction factor.

#### Quaternary complex

The largest complexes to be accounted for in our coarse-grained adiabatic approach are quaternary complexes, i.e., complexes comprised of four strands. We need to distinguish three types of such complexes: (i) 3-1 quaternary complexes, (ii) left-tilted 2-2 quaternary complexes and (iii) right-tilted 2-2 quaternary complexes. In 3-1 quaternary complexes, three Watson strands are hybridized to one Crick strand (Fig. S4), whereas in 2-2 quaternary complexes, two Watson strands are hybridized to two Crick strands (Fig. S5 and Fig. S6).

##### 3-1 quaternary complexes

**FIG. S4.**
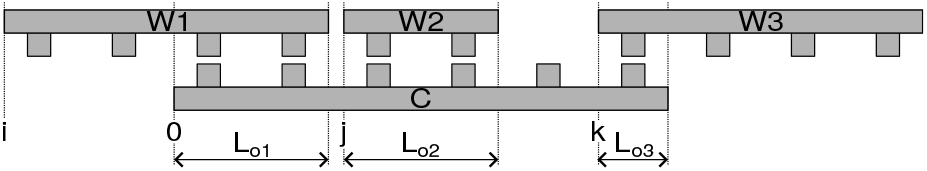
Schematic representation of a 3-1 tetraplex. Three strands (in the following referred to as Watson strands `W1, W2`, and `W3`), hybridize to a single template strand (Crick strand `C`). The positions relative to the left end of the `C` strand are given by the alignment indices *i, j*, and *k*; here, *i* = −2, *j* = 2, *k* = 5. The length of the overlap regions are denoted *L*_o,1_, *L*_o,2_ and *L*_o,3_.

Fig. S4 depicts a typical 3-1 tetraplex. Such a tetraplex is uniquely characterized by the length of its oligomers, *L*_C_, *L*_W,1_, *L*_W,2_, *L*_W,3_, as well as their relative position to each other denoted by the alignment indices *i, j*, and *k*. All positions within the triplex are measured relative to the left end of the `C` strand. Any `W` strand needs to have at least one nucleotide of overlap with the `C` strand, but two `W` strands must never occupy the same position on the `C` strand. Consequently, the alignment indices fall within the intervals,

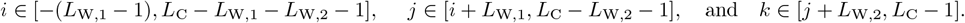

There are two dangling ends (left and right) and potentially two gaps between the `W` strands: one gap between `W1` and `W2` and another one between `W2` and `W3`. The following boolean variables indicate the presence/absence of the respective dangling ends,

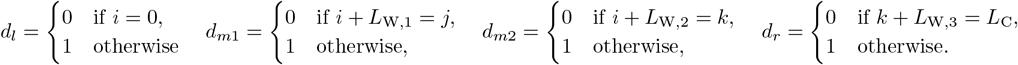

The length of the hybridization regions is given by

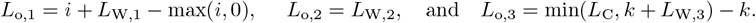

Following the same reasoning as in the case of ternary complexes, the energy equals

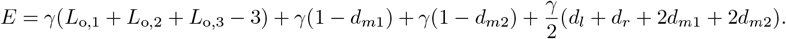

Similarly, the combinatorial multiplicity of 3-1 quaternary complexes is constructed using the same reasoning as in the case of ternary complexes,

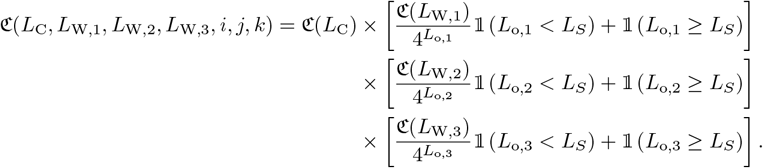

As 3-1 quaternary complexes are not symmetric under rotation, no symmetry correction of the combinatorial multiplicity is necessary.

**FIG. S5.**
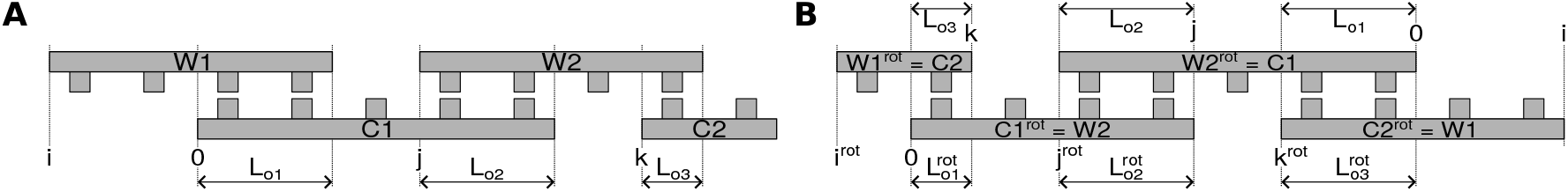
Schematic representation of a left-tilted 2-2 tetraplex. **A** Two Watson strands (`W1` and `W2`) are hybridized to two Crick strands (`C1` and `C2`). Both Watson strands are hybridized to the left Crick strand `C1`, whereas only `W2` is hybridized to the right Crick strand `C2`. The alignment indices *i, j* and *k* denote the position of the strands relative to the left end of `C1`; here, *i* = −2, *j* = 3 and *k* = 6. The length of the hybridization regions are called *L*_o,1_, *L*_o,2_ and *L*_o,3_. **B** Rotating the schematic representation of a left-tilted 2-2 tetraplex by 180^*°*^ produces an alternative representation of the same complex, which is again a left-tilted 2-2 tetraplex. The panel depicts the rotated tetraplex representation (variables with superscript “rot”) as well as the un-rotated representation (variables without superscript). There is a unique linear mapping between un-rotated and rotated representation, e.g., `C2` after rotation always corresponds to `W1` before rotation.

##### Left-Tilted 2-2 quaternary complexes

A 2-2 tetraplex is comprised of two `C` strands and two `W` strands. We call a 2-2 tetraplex left-tilted if strand `W1` is connected to strand `W2` via strand `C1` (Fig. S5A). The lengths of the oligomers are called *L*_W,1_, *L*_W,2_, *L*_C,1_ and *L*_C,2_. The positions of the strands relative to each other are governed by the alignment indices. All positions are measured relative to the position of the left end of strand `C1`. The alignment indices may take on the following values,

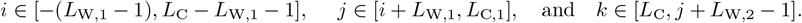

The complex can have dangling ends on the right and on the left end of the complex; the presence of these dangling ends is indicated by the boolean variables *d*_*l*_ and *d*_*r*_. Moreover, two gaps are possible: There might be a gap between strands `W1` and `W2`, or a gap between `C1` and `C2`. The respective boolean variables read

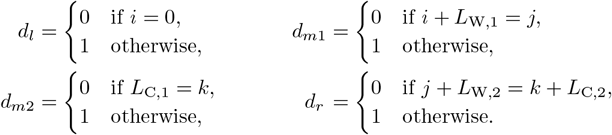

We refer to the hybridization region of strand `W1` and `C1` as overlap region 1, to the hybridization region of strand `W2` and `C1` as overlap region 2 and to the hybridization region of strand `W2` and `C2` as overlap region 3. The length of these hybridization regions is computed via

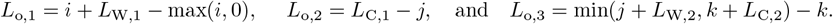

Given the length of the hybridization region as well as the presence/absence of dangling ends, we can compute the hybridization energy,

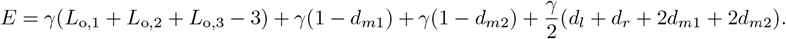

The combinatorial multiplicity of a left-tilted 2-2 tetraplex is constructed using the same reasoning as in the case of a 3-1 tetraplex,

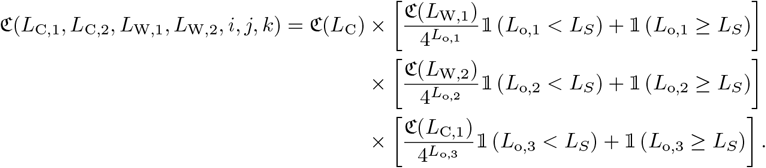

To prevent double-counting the same tetraplex, we include either the complex or its rotated representation in the container of possible complexes, but not both. If the complex is symmetric under rotation, we multiply the combinatorial multiplicity by 1/2. Given a left-tilted 2-2 tetraplex (*L*_C,1_,*L*_C,2_,*L*_W,1_,*L*_W,2_, *i, j, k*), we can compute the corresponding rotated tetraplex 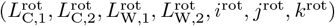 by applying a linear map. The mapping of the oligomers lengths is illustrated in Fig. S5B. We see that strand `C2` after rotation corresponds to strand `W1` before rotation, implying that 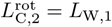. The same reasoning can be applied to all strands leading to the following map,

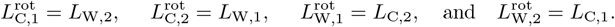

In order to compute the map of the alignment indices under rotation, we need to express the relative positions of the strands with respect to the position of strand `C1` after rotation, which corresponded to `W2` before rotation. For example, *i*^rot^ corresponds to the number of nucleotides by which strand `C2` (before rotation) protrudes beyond strand `W2` (before rotation). Expressed in terms of variables before rotation, this distance may be written as *k* +*L*_C,2_ − *jL*_W,2_. Analogous relations can be derived for all alignment indices,

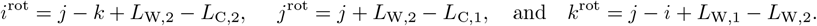

##### Right-tilted 2-2 quaternary complexes

**FIG. S6.**
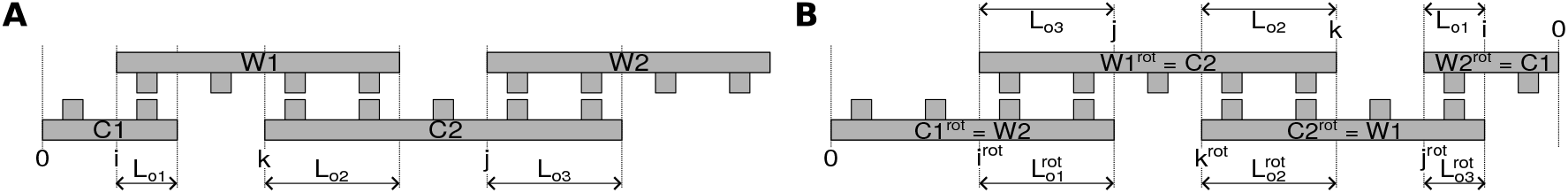
Schematic representation of a right-tilted 2-2 tetraplex. **A** Two Watson strands (`W1` and `W2`) are hybridized to two Crick strands (`C1` and `C2`). Unlike in the left-tilted 2-2 tetraplex, both Watson strands are hybridized to the right Crick strand `C2`, whereas only `W1` is hybridized to the left Crick strand `C1`. The alignment indices *i, j*, and *k* denote the positions of the strands relative to `C1`; here, *i* = 1, *k* = 3, and *j* = 6. The length of the overlap regions are called *L*_o,1_, *L*_o,2_ and *L*_o,3_. **B** Rotating the schematic representation of a right-tilted 2-2 tetraplex by 180^*°*^ produces an alternative representation of the same complex, which is again a right-tilted 2-2 tetraplex. The panel depicts the rotated tetraplex representation (variables with superscript “rot”) as well as the un-rotated representation (variables without superscript). There is a unique linear mapping between un-rotated and rotated representation, e.g., `C2` after rotation always corresponds to `W1` before rotation. The mapping is identical for left- and right-tilted 2-2 quaternary complexes.

A 2-2 tetraplex is called right-tilted if strand `W1` is connected to strand `W2` via strand `C2` (Fig. S6). As in the case of the left-tilted 2-2 tetraplex, the oligomer lengths are again called *L*_W,1_, *L*_W,2_, *L*_C,1_ and *L*_C,2_, but the values of the alignment indices that are possible for the right-tilted tetraplex differ from the ones of the left-tilted tetraplex,

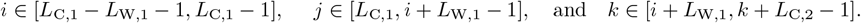

Note that the range of *i* is chosen such that at least one nucleotides of strand `W1` always extends to the right beyond the end of strand `C1`, allowing for a hybridization region between strand `C2` and `W1`. The boolean variables denoting the presence/absence of dangling ends read

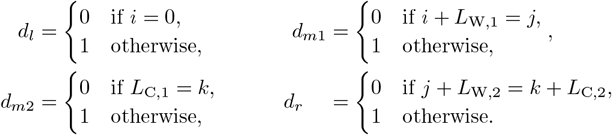

The length of the overlap regions is given by

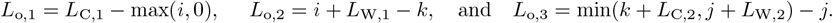

Like in the case of left-tilted 2-2 quaternary complexes, the total hybridization energy is computed via

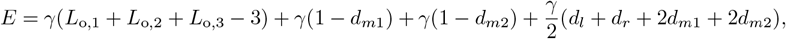

and the combinatorial multiplicity via

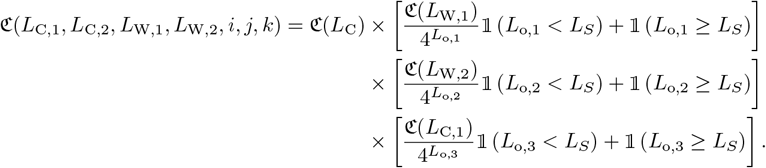

We include either the tetraplex or its rotated representation in the list of possible complexes to avoid double-counting of quaternary complexes. Moreover, the combinatorial multplicity is divided by 2, if the tetraplex is symmetric under rotation. It turns out that the rotation map for the right-tilted 2-2 tetraplex is identical to the one of the left-tilted 2-2 tetraplex,

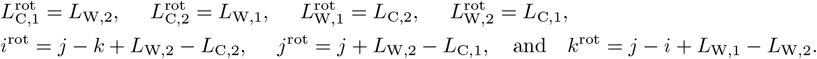

### S4. NUMERICAL SOLUTION OF THE (DE)HYBRIDIZATION EQUILIBRIUM IN THE ADIABATIC APPROACH

Based on the list of complexes constructed in the previous section, we can compute the equilibrium concentration of strands and complexes reached in the (de)hybridization equilibrium. In the following, we denote the concentration of a coarse-grained oligomer with length *L* as *c*(*L*), and the concentration of an oligomer with length *L* and known sequence as *c*_*s*_(*L*). Recall that we assumed that all sequences of a given length that are compatible with the circular genome are equally likely in the pool (Section S3). Thus, the concentration of the coarse-grained oligomer and the concentration of an oligomer with specified sequence are related by the combinatorial multiplicity,

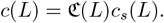

In order to compute the concentration of a complex based on the concentration of single strands, we make use of the law of mass action. The concentration of a specific sequence realization of a complex is computed as the product of concentrations of the strands forming the complex divided by the dissociation constant *K*_*d*_,

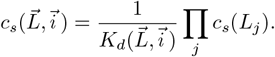

Here, 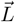 is the vector denoting the lengths of the strands comprising the complex, and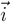 are the alignment indices. The dissociation constant is set by the hybridization energy, Δ*G*, of the complex,

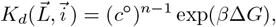

where *c*^°^ = 1 M is the standard concentration. Just as in the case of single strands, the concentration of the sequence-independent coarse-grained complex is related to the concentration of a complex with specific sequence realization via the combinatorial prefactor,

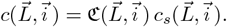

It can be useful to combine the combinatorial multiplicity and the dissociation constant into a single effective association constant,

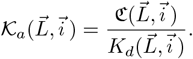

Note that the effective association constant including the combinatorial multiplicity is denoted by 𝒦_*a*_ (in curly font), while the association constant without combinatorial multiplicity is denoted 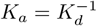 (in regular font).

In the adiabatic approach, we study the behavior of the system on timescales that are long enough for the system to reach the (de)hybridization equilibrium, but too short for templated ligation events to take place. Therefore, the length of the oligomers are expected not to change throughout the equilibration process, and we need to introduce a separate mass conservation law for each coarse-grained oligomer, i.e., each oligomer length, that is included in the pool. For each length, the concentration of single-stranded coarse-grained oligomers of length *L* and the concentration of coarse-grained oligomers of length *L* bound in a complex need to add up to their total concentration *c*^tot^(*L*) set by the initial condition,

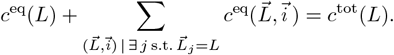

In this equation, 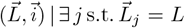 denotes the summation over all complexes that contain at least one strand of length *L*. Note that we referred to the total concentration of oligomers of length *L* as *c*(*L*) in the main text, but for clarity we use *c*^tot^(*L*) in the supplementary material.

Combining the mass conservation requirement with the mass action law gives a set of coupled polynomial equations. The number of equations equals the number of distinct oligomer lengths included in the pool. The polynomial equations are of degree 4, as quaternary complexes are the largest complexes to be accounted for and their concentrations equals the product of the four strands comprising the complex. We determine the equilibrium concentrations by finding the root of this set of fourth-degree polynomial equations using the Levenberg-Marquardt algorithm.

### S5. COMPUTING REPLICATION OBSERVABLES BASED ON THE ADIABATIC APPROACH

**FIG. S7.**
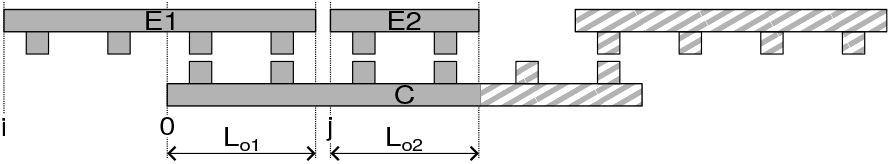
Schematic representation of a complex that allows for templated ligation. The strands `E1` and `E2` are adjacent to each other, such that a covalent bond can form between their ends. The length of the product strand, *L*_P_, is set by the length of the educt strands, *L*_e,1_ and *L*_e,2_. The likelihood for the complex to form a product oligomer whose sequence is compatible with the true circular genome, *p*_corr_, is determined by the length of the educts and the length of their hybridization region with the template. The parts of the complex that are depicted with hatching do not affect *p*_corr_.

As we are not modeling templated ligation events explicitely in the adiabatic approach, we compute replication observables based on the equilibrium concentration of complexes that are in a configuration which allows for a templated ligation reaction to happen. Templated ligations are possible if two strands in the complex are adjacent to each other, i.e., there is no gap in between two oligomers that are hybridized to the same template strand (Fig. S7). Recall that the absence of a gap between two oligomers in the complex implies that the dangling end indicator variable *d*_*m*_ = 0 (Section S3). The length of the product strand `P` is equal to the sum of the lengths of the two educt strands `E1` and `E2`, *L*_P_ = *L*_E,1_ +*L*_E,2_. We can use the information about the length of the product strand to compute the yield of replication. By definition (see main text), the yield equals the fraction of nucleotides used to form VCG oligomers, i.e., strands longer than *L*_*S*_,

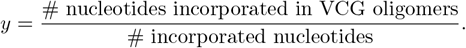

We can express this quantity in terms of the equilibrium concentration of complexes facilitating templated ligation,

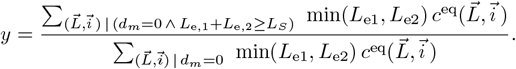

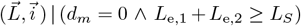 denotes the summation over all complexes, in which (i) the strands `E1` and `E2` are adjacent to each other, i.e., *d*_*m*_ = 0, and (ii) the length of the product *L*_e,1_ + *L*_e,2_ ≥ *L*_*S*_. We multiply the equilibrium concentration by the length of the shorter educt strand to account for the number of incorporated nucleotides in line with the definition of the yield, Eq. (S2a).

In order to compute the fidelity of replication, we need to distinguish between product oligomers sequences that are compatible with the genomes (correct sequences) and sequences that are incompatible with the genome (false sequences). As we do not know about the details of the sequences due to the coarse-grained representation of the complexes, we need to invoke a combinatorial argument to determine the fraction of correct products. To this end, we compare the number of product sequences that might be produced in a complex of given oligomer lengths and relative oligomer position, to the number of correct products. The combinatorial multiplicity of the products that **could** be produced by a complex of given configuration is set by the combinatorial multiplicity of the possible templates, ℭ (*L*_o,1_ + *L*_o,2_), multiplied by the multiplicity of the educt strands hybridizing to the template with given lengths of the hybridization regions, *L*_o,1_ and *L*_o,2_,

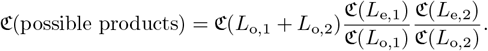

The multiplicity of **correct** products equals the combinatorial multiplicity of strands that have the same length as the product,

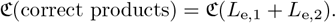

This implies that the probability for a complex of given shape 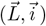 to form a correct product is given by

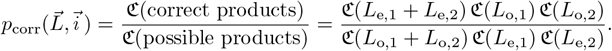

Using this probability, we can compute the fidelity of replication,

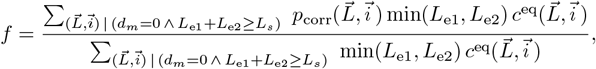

as well as the replication efficiency,

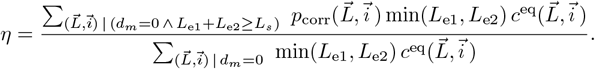

### S6. SCALING LAW OF THE EQUILIBRIUM CONCENTRATION RATIO THAT MAXIMIZES REPLICATION EFFICIENCY

We consider a pool containing monomers and VCG oligomers of a single length. As the feedstock only contains monomers, the total monomer concentration and the total feedstock concentration are identical, 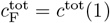. Similarly, the total VCG concentration equals the total concentration of oligomers of length *L*_V_,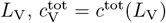. In such a system, the replication efficiency can be expressed as

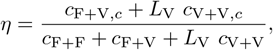

where the concentrations denote the equilibrium concentration of all complexes facilitating the indicated type of templated ligation. Note that we do not include dimerizations, i.e., F+F ligations, in the numerator as they do not contribute to the yield. Moreover, the V+V ligations are multiplied by the length of the oligomers *L*_V_, as each V+V extends an existing oligomer by *L*_V_ nucleotides.

Assuming that complexes comprised of three strands are the dominant type of complexes, we can express the equilibrium concentrations as product of the equilibrium concentrations of the free strands and the effective association constants,

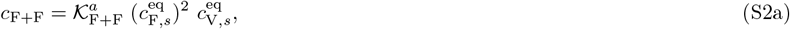

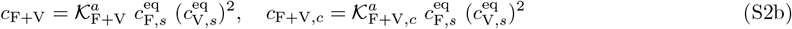

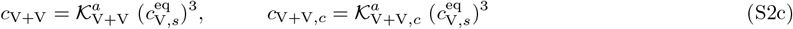

Here, 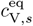 denotes the equilibrium concentration of a VCG oligomer with a specific sequence; the equilibrium concentration of all VCG oligomers for any sequence 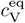 is computed by multiplying with the combinatorial multiplicity, 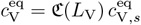. The same holds for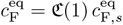. Eq. (S2) allows us to write the replication efficiency as,

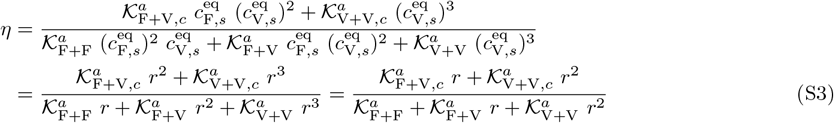

where we introduced the ratio of equilibrium concentrations, 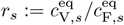. Maximizing Eq. (S3) with respect to *r*_*s*_ yields,

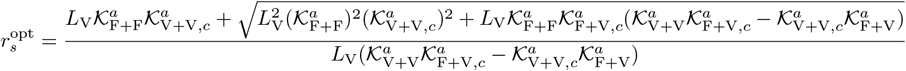

The effective association constant are computed using the combinatorial rules outlined in Section S4. We find that 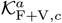 is similar to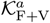, deviating at most by 10%. Moreover,

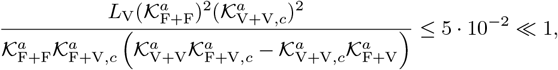

and,

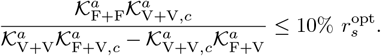

These observations allow us to obtain an significantly simplified approximate expression for 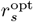,

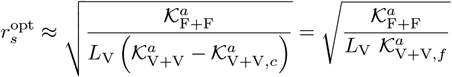

We used the identity 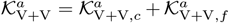 in the last step. To derive the scaling of 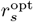 as a function of *L*_V_, we need to analyze how the effective association constants depend on *L*_V_.

1. 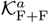 scales linearly with *L*_V_. In F+F ligations, an VCG oligomer acts as template facilitating the ligation of two monomers. Independent of the length of the VCG oligomer, the two monomers always have the same binding affinity to their template. However, the number of possible positions that the two adjacent monomers can have on the template scales linearly in template length, causing the effective association constant to depend linearly on *L*_V_,

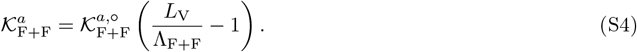
2. 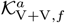scales exponentially with *L*_V_. As illustrated in Fig. 2**Replication performance of VCG pools containing VCG oligomers of a single length (single-length VCG pools). (A)** The pool contains a fixed concentration of monomers, 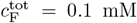, as well as VCG oligomers of a single length, *L*_V_, at variable concentration 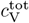(the VCG oligomers cover all possible subsequences of the genome and its complement at equal concentration). **(B)** The yield increases as a function of 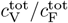, because dimerizations become increasingly unlikely for high VCG concentrations. **(C)** The ligation share of different ligation types depends on the total VCG concentration: In the low concentration limit, dimerization (F+F) dominates; for intermediate concentrations, F+V ligations reach their maximum, while, for high concentrations, a substantial fraction of reactions are V+V ligations. The panel depicts the behavior for *L*_V_ = 6 nt. **(D)** Replication efficiency is limited by the small yield for small 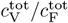. In the limit of high 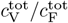, replication efficiency decreases due to the growing number of error-prone V+V ligations. Maximal replication efficiency is reached at intermediate VCG concentration. **(E)** V+V ligations are prone to the formation of incorrect products due to the short overlap between educt strand and template. In general, the probability of correct product formation, *p*_corr_, depends on the choice of circular genome and as well as its mapping to the VCG pool. The probabilities listed here refer to a VCG pool with *L*_*G*_ = 16 nt, *L*_*E*_ = 2 nt and *L*_*U*_ = 3 nt. **(F)** The optimal equilibrium concentration ratio of free VCG strands to free feedstock strands, which maximizes replication efficiency, decays as a function of length (continuous line). The analytical scaling law (dashed line, Eq. (4equation.8)) captures this behavior. The window of close-to-optimal replication, within which efficiency deviates no more than 1% from its optimum (shaded areas), increases with *L*_V_, facilitating reliable replication without fine-tuning to match the optimal concentration ratio. **(G)** Maximal replication efficiency, which is attained at the optimal VCG concentration depicted in panel E, increases as a function of *L*_V_ and approaches a plateau of 100%. For high efficiency, Eq. (5equation.9) provides a good approximation of the length-dependence of *η*_max_ (dashed lines). The oligomer length at which replication efficiency equals 95% is determined using Eq. (5equation.9) (vertical dotted lines). **(H)** The unique motif length, 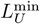, increases logarithmically with the length of the genome, *L*_*G*_. The length of VCG oligomers, *L*_V_, at which the optimal replication efficiency reaches 95% (computed using Eq. (5equation.9)) exhibits the same logarithmic dependence on *L*_*G*_.figure.caption.7E in the main text, the length of the hybridization region equals *L* in complexes facilitating incorrect V+V ligations. This leads to exponential scaling of 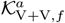 with *L*_V_, as the hybridization energy is proportional to the length of the hybridization site. The number of complexes facilitating incorrect product formation is only a function of the unique subsequence length *L*_*S*_, but does not depend on the oligomer length *L*_V_. Therefore, there is no multiplicative prefactor proportional *L*_V_,

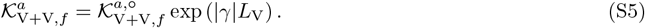

**FIG. S8.**
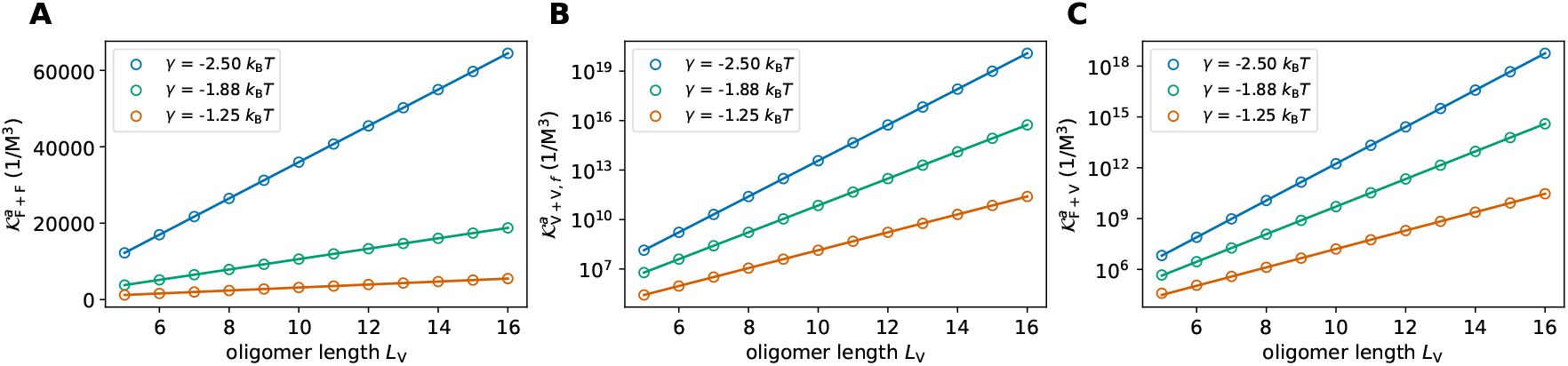
Effective association constants of complexes facilitating F+F ligations (**A**), false V+V ligations (**B**), and F+V ligations (**C**). The dots depict the effective association constants derived based on the combinatoric rules presented in S4, the solid lines represent the respective scaling laws introduced in Eq. (S4-S7). Different colors correspond to different hybridization energies per matching nearest neighbor block *γ*.

Fig. S8 shows the *L*_V_-dependence of the effective association constants. The circles represent the effective association constants derived based on the combinatoric rules discussed in Section S4, while the solid lines show the length-dependent scaling computed via Eq. (S4) and (S5).

Given the scaling of 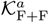 and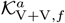, we find that the optimal ratio of equilibrium concentrations 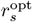 equals

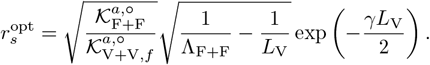

We obtain *r*^opt^ (Fig. 2**Replication performance of VCG pools containing VCG oligomers of a single length (single-length VCG pools). (A)** The pool contains a fixed concentration of monomers, 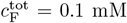, as well as VCG oligomers of a single length, *L*_V_, at variable concentration 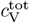(the VCG oligomers cover all possible subsequences of the genome and its complement at equal concentration). **(B)** The yield increases as a function of 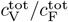, because dimerizations become increasingly unlikely for high VCG concentrations. **(C)** The ligation share of different ligation types depends on the total VCG concentration: In the low concentration limit, dimerization (F+F) dominates; for intermediate concentrations, F+V ligations reach their maximum, while, for high concentrations, a substantial fraction of reactions are V+V ligations. The panel depicts the behavior for *L*_V_ = 6 nt. **(D)** Replication efficiency is limited by the small yield for small 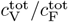. In the limit of high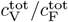, replication efficiency decreases due to the growing number of error-prone V+V ligations. Maximal replication efficiency is reached at intermediate VCG concentration. **(E)** V+V ligations are prone to the formation of incorrect products due to the short overlap between educt strand and template. In general, the probability of correct product formation, *p*_corr_, depends on the choice of circular genome and as well as its mapping to the VCG pool. The probabilities listed here refer to a VCG pool with *L*_*G*_ = 16 nt, *L*_*E*_ = 2 nt and *L*_*U*_ = 3 nt. **(F)** The optimal equilibrium concentration ratio of free VCG strands to free feedstock strands, which maximizes replication efficiency, decays as a function of length (continuous line). The analytical scaling law (dashed line, Eq. (4equation.8)) captures this behavior. The window of close-to-optimal replication, within which efficiency deviates no more than 1% from its optimum (shaded areas), increases with *L*_V_, facilitating reliable replication without fine-tuning to match the optimal concentration ratio. **(G)** Maximal replication efficiency, which is attained at the optimal VCG concentration depicted in panel E, increases as a function of *L*_V_ and approaches a plateau of 100%. For high efficiency, Eq. (5equation.9) provides a good approximation of the length-dependence of *η*_max_ (dashed lines). The oligomer length at which replication efficiency equals 95% is determined using Eq. (5equation.9) (vertical dotted lines). **(H)** The unique motif length, 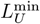, increases logarithmically with the length of the genome, *L*_*G*_. The length of VCG oligomers, *L*_V_, at which the optimal replication efficiency reaches 95% (computed using Eq. (5equation.9)) exhibits the same logarithmic dependence on *L*_*G*_.figure.caption.7) from 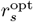 by multiplying with the appropriate combinatorial multiplicity,

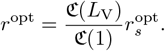

### S7. CHARACTERISTIC LENGTH-SCALE OF VCG OLIGOMERS ENABLING HIGH REPLICATION EFFICIENCY

Starting from Eq. S3, the optimal efficiency of replication can be expressed as

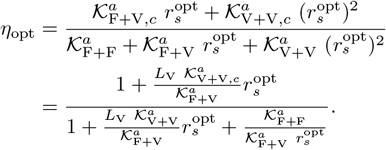

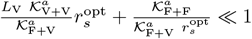 (for sufficiently high *L*_V_), we can expand in this small argument,

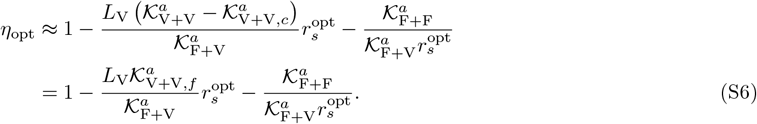

To understand the scaling of *η*_opt_ as function of *L*_V_, we need to know the scaling of 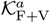 in *L*_V_. A typical complex allowing for a F+V ligation is depicted in Fig. 2**Replication performance of VCG pools containing VCG oligomers of a single length (single-length VCG pools). (A)** The pool contains a fixed concentration of monomers, 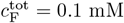, as well as VCG oligomers of a single length, *L*_V_, at variable concentration 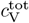 (the VCG oligomers cover all possible subsequences of the genome and its complement at equal concentration). **(B)** The yield increases as a function of 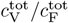, because dimerizations become increasingly unlikely for high VCG concentrations. **(C)** The ligation share of different ligation types depends on the total VCG concentration: In the low concentration limit, dimerization (F+F) dominates; for intermediate concentrations, F+V ligations reach their maximum, while, for high concentrations, a substantial fraction of reactions are V+V ligations. The panel depicts the behavior for *L*_V_ = 6 nt. **(D)** Replication efficiency is limited by the small yield for small 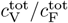. In the limit of high 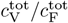, replication efficiency decreases due to the growing number of error-prone V+V ligations. Maximal replication efficiency is reached at intermediate VCG concentration. **(E)** V+V ligations are prone to the formation of incorrect products due to the short overlap between educt strand and template. In general, the probability of correct product formation, *p*_corr_, depends on the choice of circular genome and as well as its mapping to the VCG pool. The probabilities listed here refer to a VCG pool with *L*_*G*_ = 16 nt, *L*_*E*_ = 2 nt and *L*_*U*_ = 3 nt. **(F)** The optimal equilibrium concentration ratio of free VCG strands to free feedstock strands, which maximizes replication efficiency, decays as a function of length (continuous line). The analytical scaling law (dashed line, Eq. (4equation.8)) captures this behavior. The window of close-to-optimal replication, within which efficiency deviates no more than 1% from its optimum (shaded areas), increases with *L*_V_, facilitating reliable replication without fine-tuning to match the optimal concentration ratio. **(G)** Maximal replication efficiency, which is attained at the optimal VCG concentration depicted in panel E, increases as a function of *L*_V_ and approaches a plateau of 100%. For high efficiency, Eq. (5equation.9) provides a good approximation of the length-dependence of *η*_max_ (dashed lines). The oligomer length at which replication efficiency equals 95% is determined using Eq. (5equation.9) (vertical dotted lines). **(H)** The unique motif length, *L*^min^, increases logarithmically with the length of the genome, *L*_*G*_. The length of VCG oligomers, *L*_V_, at which the optimal replication efficiency reaches 95% (computed using Eq. (5equation.9)) exhibits the same logarithmic dependence on *L*_*G*_.figure.caption.7E in the main text. The length of the overlap *L*_o_ region between template and educt VCG oligomer can vary; it is at least one nucleotide, and at most *L*_V_ − 1 nucleotides (one base pairing position needs to remain available for the monomer). We obtain the effective association constant by summing the contributions of all of these configurations,

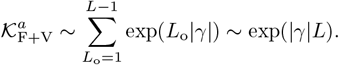

Therefore, we use the following scaling ansatz for 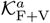,

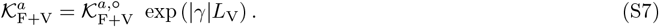

Combining the scaling laws for 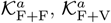, and 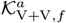 with Eq. S6, we find

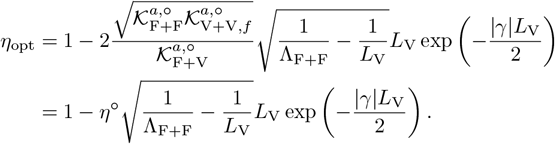

### S8. DEPENDENCE OF REPLICATION EFFICIENCY ON SUBSEQUENCE LENGTHSCALES OF THE CIRCULAR GENOME

The coarse-grained representation (Section S3) used in the adiabatic approach applies only to a specific class of genomes – those in which the exhaustive coverage length *L*_*E*_ is maximal and the unique motif length *L*_*U*_ is minimal. This corresponds to genomes that satisfy 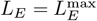 and 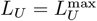 (see Section S1 for details).

To analyze replication behavior beyond this limited case, we developed a fully sequence-resolved extension of the adiabatic approach. Rather than using a coarse-grained view of oligomers, this method considers each distinct strand sequence as a separate chemical species. For example, a genome of length *L*_*G*_ = 64 nt that is encoded in a pool containing monomers and octamers, a total of 132 individual oligomers must be tracked: 4 monomers and 128 distinct octamers. This contrasts sharply with the coarse-grained scenario, which involves only two variables (total monomer and total octamer concentrations). Starting from the single-stranded oligomers, all possible complexes involving up to three strands are enumerated, and their hybridization free energies are calculated using the same energetic model applied in the coarse-grained framework. In the aforementioned example, this results in 351 200 distinct sequence-resolved complexes, compared to just 135 complexes in the coarse-grained model, highlighting the increased computational demands in terms of memory and runtime. The hybridization equilibrium is computed by solving the algebraic system defined by the law of mass action and mass conservation. The procedure mirrors that of the coarse-grained approach (Section S4), with the key difference that combinatorial prefactors are no longer required: these are inherently encoded in the full enumeration of unique sequences and their binding configurations.

For our further analysis, we fix the genome length at *L*_*G*_ = 64 nt and systematically vary the motif length scales *L*_*E*_ and *L*_*U*_. Genomes with desired values of *L*_*E*_ and *L*_*U*_ are constructed as described in Section S1: Starting from a genome with maximal motif entropy 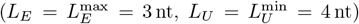, we reduce the motif entropy on intermediate length scales *L*_*E*_ *< L < L*_*U*_ to achieve the target characteristics. Table S2 provides an overview of all sampled genome sequences. We consider two limiting cases for motif distributions between *L*_*E*_ and *L*_*U*_. In the weakly biased case, the motif distribution (for motifs of length *L*_*E*_ *< L < L*_*U*_) is nearly uniform: most motifs of length *L* appear exactly once, except for a few motifs that occur twice. This leads to genomes where *L* ≠ *L*_*U*_, but the deviation from maximal entropy is minimal. In the strongly biased case, many motifs appear multiple times, yielding a much less uniform distribution.

**TABLE S2.**
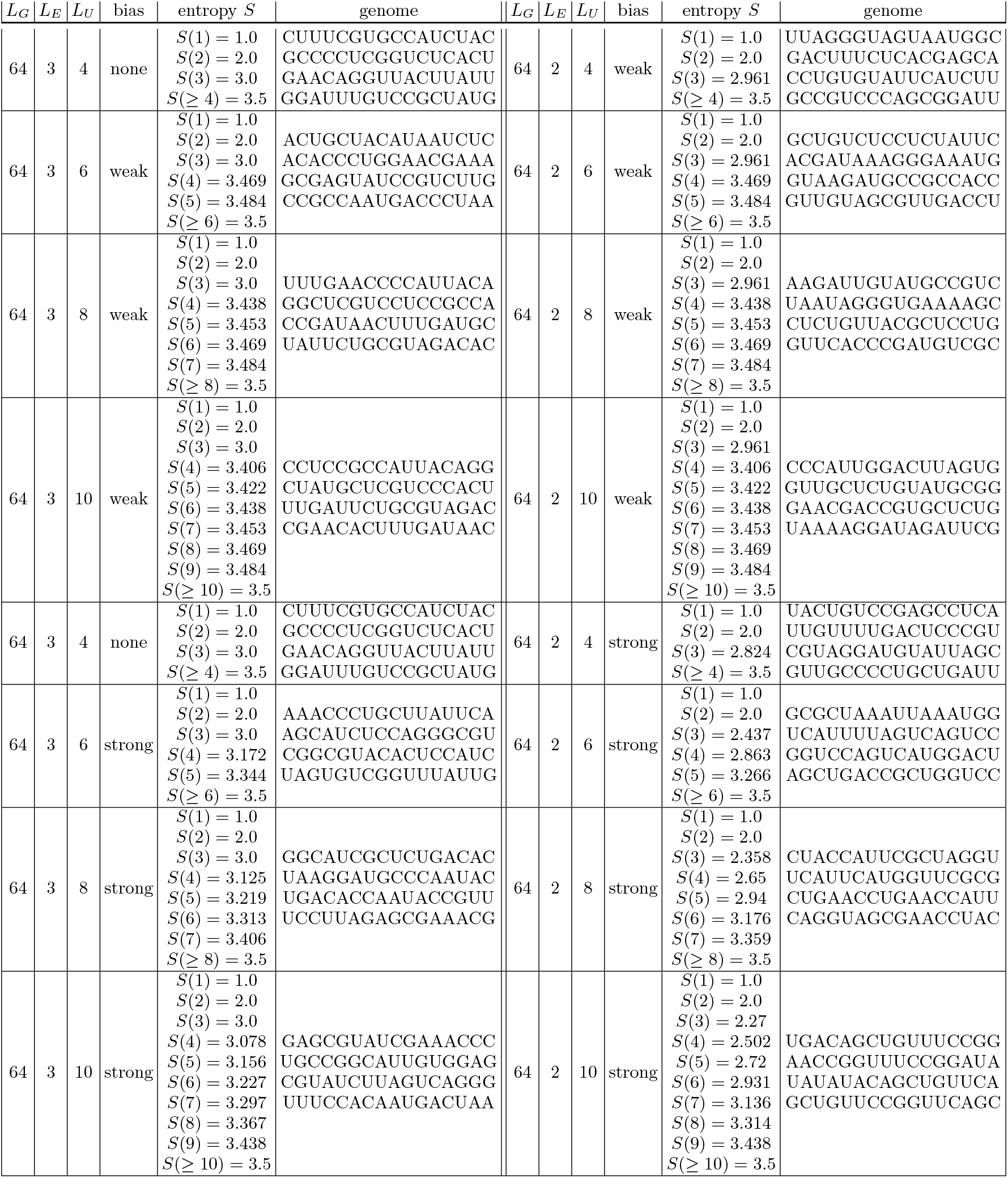
Genomes sampled via the Metropolis-Hastings algorithm using the motif entropy as “Hamiltonian”. The table summarizes the sequence of the genome, its characteristic length scales *L*_*E*_ and *L*_*U*_, as well as the motif entropy on all length scales of interest. The keyword “bias” is used to distinguish two different sampling procedures: Weakly biased genomes are designed to obey the desired length scales *L*_*E*_ and *L*_*U*_ while retaining a close-to-uniform motif distribution for subsequences of length *L*_*E*_ *< L < L*_*U*_, whereas the motif distribution is far from uniform for strongly biased genomes.

**FIG. S9.**
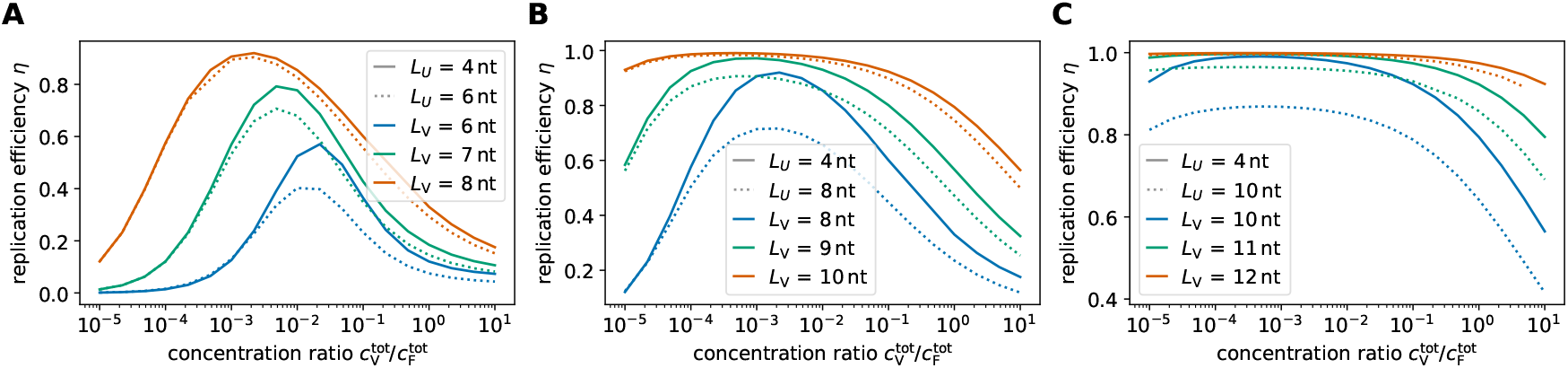
Replication efficiency as a function of the concentration of VCG oligomers in the pool for different choices of genomes and varying VCG oligomer length. All genomes are *L*_*G*_ = 64 nt long, and include all motifs up to length *L*_*E*_ = 2 nt, but differ with respect to their minimal unique subsequence length *L*_*U*_ : (A) *L*_*U*_ = 6 nt, (B) *L*_*U*_ = 8 nt, and (`C`) *L*_*U*_ = 10 nt (shown as dotted lines). For comparison, every panel shows the replication efficiency of a genome with *L*_*E*_ = 3 nt, *L*_*U*_ = 4 nt (solid line). Different colors are used to distinguish different VCG oligomer lengths. Under otherwise identical conditions (e.g., identical oligomer length), replication proceeds with lower efficiency in genomes with higher unique subsequence length *L*_*U*_.

**FIG. S10.**
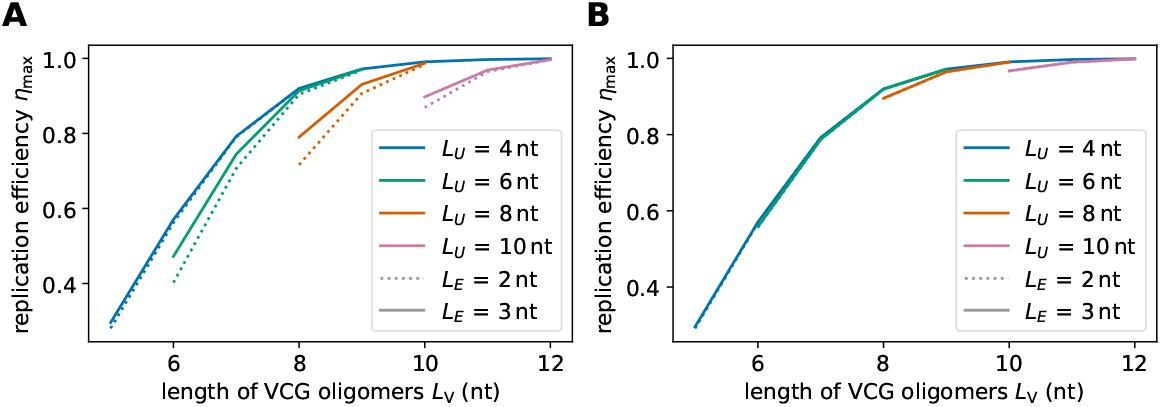
Maximal replication efficiency as a function of the oligomer length for different genomes (all *L*_*G*_ = 64 nt long). Regardlesss of the genome, the oligomer lenght needs to exceed *L*_*U*_ to enable replication with high efficiency (e.g., higher than 95%). The difference between the genome length required for high efficiency replication and the unique motif length *L*_*U*_ depends on the motif distribution on intermediate length scales (*L*_*E*_ *< L < L*_*U*_): Genomes with strong bias require longer oligomers (**A**) than genomes with weak bias (**B**) (see Table S2 for the genomes and their motif entropies).

Each genome is mapped to a Virtual Circular Genome (VCG) pool containing monomers and oligomers of variable length *L*_V_. Assuming all oligomers are chemically activated, we compute the replication efficiency of each pool as a function of the relative oligomer concentration, for different values of *L*_V_ and across genome types (Fig. S9). As expected, replication efficiency exhibits a maximum at intermediate oligomer concentrations. The maximum efficiency achieved depends on both the VCG oligomer length and the genome’s motif properties. In general, longer oligomers enable more efficient replication. However, at a fixed oligomer length, genomes with higher unique motif lengths *L*_*U*_ (or lower *L*_*E*_) replicate with lower efficiency, implying that these genomes require longer oligomers for successful replication. For example, a VCG pool with oligomers of length *L*_V_ = 9 nt can replicate a genome with *L*_*E*_ = 3 nt and *L*_*U*_ = 4 nt at 97% efficiency (see solid green curve in Fig. S9B). In contrast, replicating a genome with *L*_*E*_ = 3 nt and *L*_*U*_ = 10 nt requires oligomers of at least *L*_V_ = 11 nt to reach comparable efficiency (see dotted green curve in Fig. S9C).

Across all tested genomes, the oligomer length required to achieve *>* 95% efficiency (denoted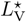) consistently exceeds the unique motif length *L*_*U*_ (Fig. S10). However, the difference 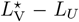 depends on the structure of the motif distribution in the intermediate length range *L*_*E*_ *< L < L*_*U*_. Genomes with strongly biased motif distributions require longer oligomers (larger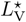, see Fig. Fig. S10A), whereas the replication behavior of weakly biased genomes closely mirrors that of the fully entropic genome (*L*_*E*_ = 3 nt, *L*_*U*_ = 4 nt) (Fig. Fig. S10B). In realistic prebiotic settings, genomes are likely to fall somewhere between these extremes, depending on the strength of biases introduced during their emergence.

### S9. ENHANCED REPLICATION EFFICIENCY IN MULTI-LENGTH VCG POOLS

We investigate if a pool containing a range of VCG oligomer lengths exhibits higher replication efficiency than single-length pools. To this end, we study a VCG pool comprised of monomers (with fixed concentration), as well as tetramers and octamers (with variable concentration). For moderate binding affinity, *γ* = −2.5 *k*_B_*T*, the pool reaches optimal replication efficiency, if the tetramer concentration is smaller than the octamer concentration (Main Text Fig. 3**Replication performance of pools containing VCG oligomers of two different lengths. (A)** The pool contains a fixed concentration of monomers, 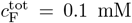, as well as tetramers and octamers at variable concentration. The hybridization energy per nearest-neighbor block is *γ* = −2.5 *k*_B_*T*. **(B)** Replication efficiency reaches its maximum for *c*^tot^(8) ≈ 0.1 µM and significantly lower tetramer concentration, *c*^tot^(4) ≈ 7.4 pM. Efficiency remains close-to-maximal on a plateau around the maximum spanning almost two orders of magnitude in tetramer and octamer concentration. In addition, efficiency exhibits a ridge of increased efficiency for high tetramer concentration and intermediate octamer concentration. **(C)** Complexes that facilitate templated ligation are grouped by the length of the template and the educts, 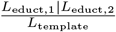. We distinguish complexes producing correct (labeled “c”) and false products (labeled “f”). For each relevant type of complex, we highlight the region in the concentration plane where it contributes most significantly, i.e., at least 20% of the total ligation flux. The plateau of high efficiency is dominated by the ligation of monomers to octamers, whereas the ridge of increased efficiency is due to the correct ligation of two tetramers templated by an octamer.figure.caption.10), with extensions of octamers by monomers (1+8) being the most dominant type of templated ligation reactions. For strong binding affinity, *γ* = −5.0 *k*_B_*T*, templated ligations of tetramers on octamer templates eventually achieve the same replication efficiency as (1+8) ligations (Fig. S11). The mechanism relies on a significant difference in the length of the template (octamers) and educt strands (tetramers). If longer educt strands (heptamers) are chosen, highest efficiency is achieved in the parameter regime, in which VCG oligomers are extended by monomers, regardless of the choice of hybridization energy per matching nearest-neighbor block, *γ* (Fig. S12 and Fig. S13).

**FIG. S11.**
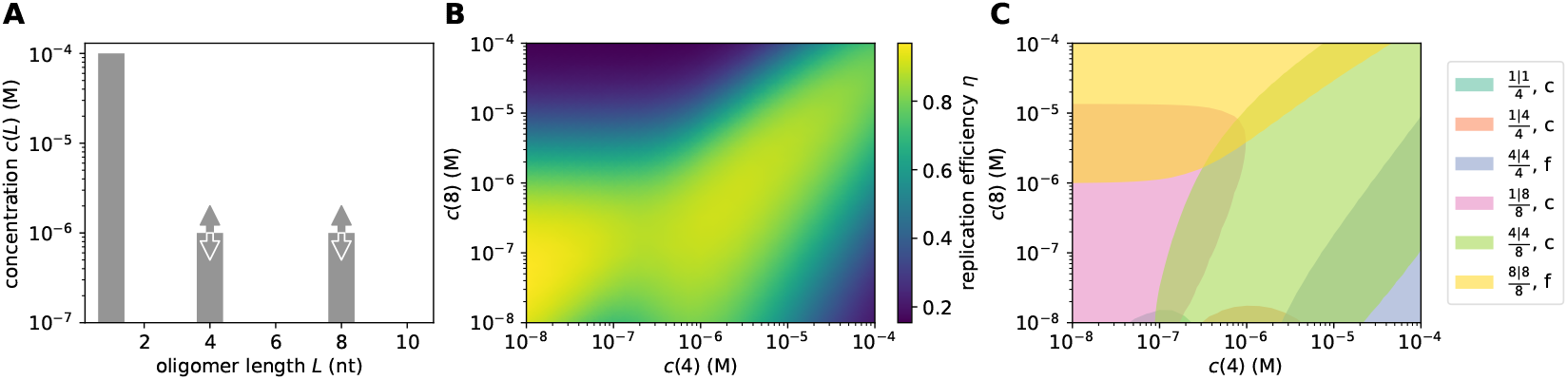
Length-modulated enhanced ligation in pools containing tetramers and octamers for strong binding affinity, *γ* = −5.0 *k*_B_*T*. The replication efficiency reaches its maximum in the concentration regime that supports templated ligation of tetramers on octamer templates.

**FIG. S12.**
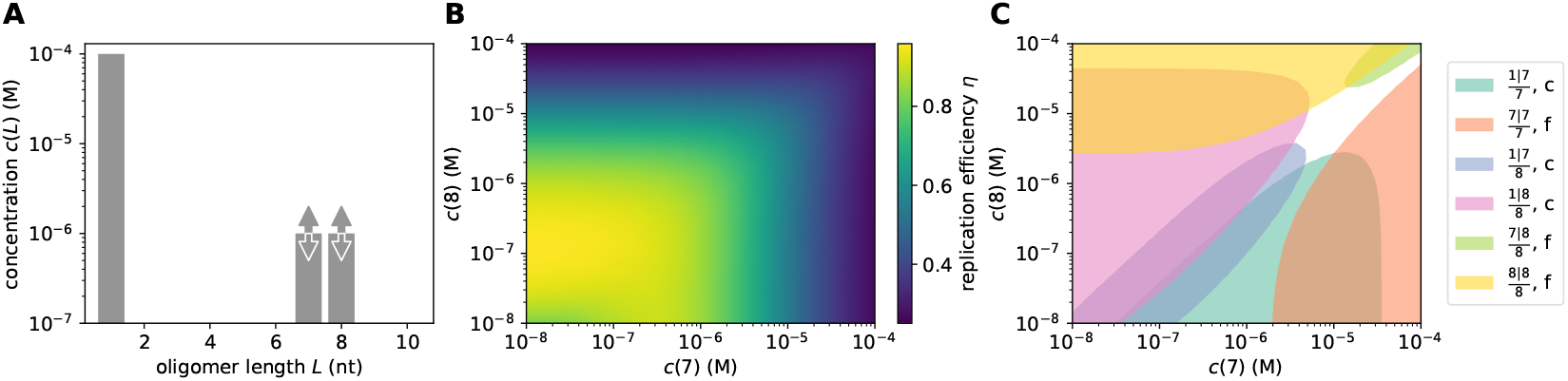
Length-modulated enhanced ligation in pools containing heptamers and octamers for weak binding affinity, *γ* = −2.5 *k*_B_*T*. The replication efficiency reaches its maximum in the concentration regime that is dominated by the addition of monomers to the VCG oligomers.

**FIG. S13.**
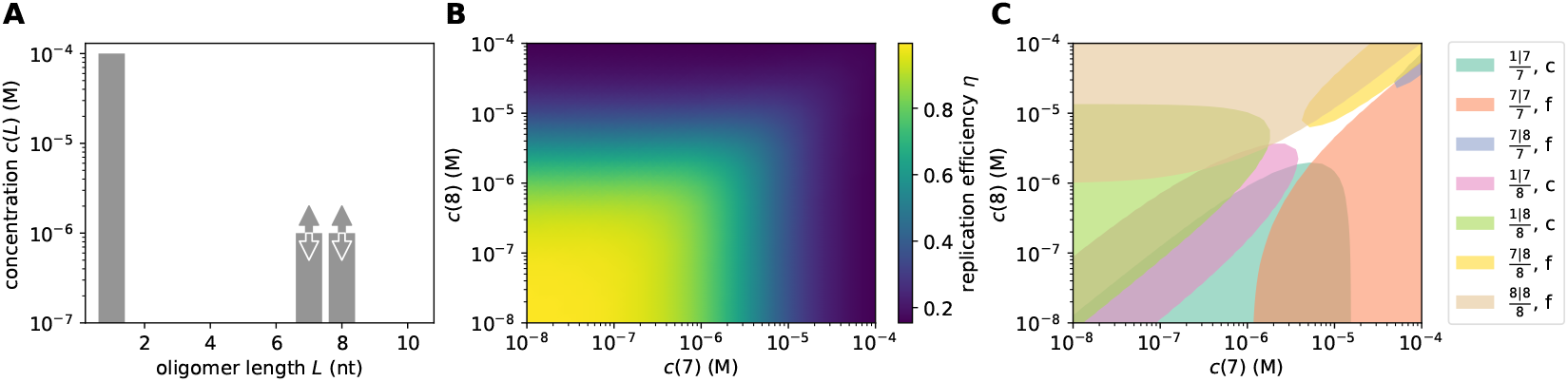
Length-modulated enhanced ligation in pools containing heptamers and octamers for strong binding affinity, *γ* = −5.0 *k*_B_*T*. The replication efficiency reaches its maximum in the concentration regime that is dominated by the addition of monomers to the VCG oligomers.

### S10. DIMER-RELATED UPPER BOUND OF REPLICATION EFFICIENCY

In a VCG pool comprised of monomers, dimers and VCG oligomers of single length, the replication efficiency can be expressed as

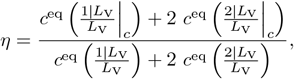

provided the pool operates in the concentration regime where all types of ligations other than F+V ligations are negligible. In this equation, 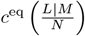 stands for the equilibrium concentration of all complexes allowing for the templated ligation of oligomers of length *L* to oligomers of length *M* if templated by oligomers of length *N*. 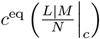 denotes the same concentration under the additional constraint that the product oligomers need to be correct.

Complexes allowing for templated ligation need to involve at least three oligomers, but may be comprised of more strands. For the following analytical derivation, we restrict ourselves to ternary complexes, while the full numerical solution (see continuous lines in Main Text Fig. 5**Replication performance of single-length VCG pools containing monomers and dimers as feedstock. (A)** The pool contains a fixed total concentration of feed-stock, 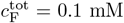, partitioned into monomers and dimers, as well as VCG oligomers of a single length, *L*_V_. The proportion of monomers and dimers can be adjusted via *κ*_F_, and the concentration of the VCG oligomers is a free parameter, 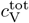. **(B)** Replication efficiency exhibits a maximum at intermediate VCG concentration in systems with (dashed blue curve) and without dimers (solid blue curve). The presence of dimers reduces replication efficiency significantly, as they enhance the ligation share of incorrect F+V ligations (dashed green curve). The panel depicts the behavior for *L*_V_ = 7 nt and *κ*_F_ = 2.3. **(C)** Optimal replication efficiency increases as a function of oligomer length, *L*_V_, and asymptotically approaches a plateau (dashed lines, Eq. (6equation.13)). The value of this plateau, 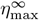, is determined by the competition between correct and false 2+V reactions, both of which grow exponentially with *L*_V_. Thus, 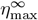 depends on the relative concentration of the dimers in the pool: the more dimers are included, the lower is 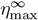. **(D)** Erroneous 1+V ligations are possible if the educt oligomer has a short overlap region with the template. The hybridization energy for such configurations is small, and independent of the length of the VCG oligomers (left). While 2+V ligations may produce incorrect products via the same mechanism (middle), they can also be caused by complexes in which two VCG oligomers hybridize perfectly to each other, but the dimer has a dangling end. The stability of these complexes increases exponentially with oligomer length (right).figure.caption.12C) also includes quaternary complexes. If only ternary complexes are accounted for, each equilibrium concentration may be written as a product of the effective association constant and the equilibrium concentrations of the involved oligomers,

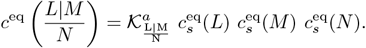

**FIG. S14.**
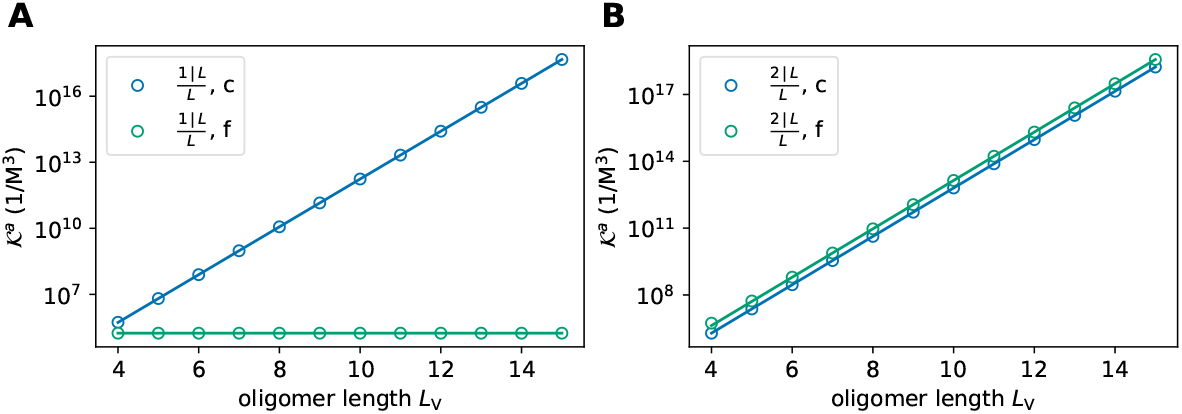
Effective association constants of complexes facilitating 1+V ligations (**A**) and 2+V ligations (**B**).

Using this expression, the equation for the replication efficiency may be simplified,

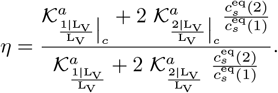

The effective association constants in the denominator may be expressed as the sum of the association constants of correct and false products,

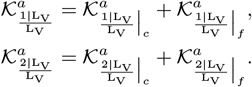

In the following, we simplify the notation by referring to 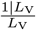 as 1+V ligations, and to 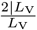 as 2+V ligations. The effective association constants differ with respect to their dependence on *L*_V_: While 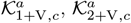 and 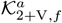 increase exponentially in 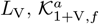 is length-independent (Fig. S14). Introducing the parametrizations,

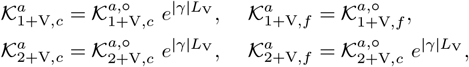

we can express *η* as

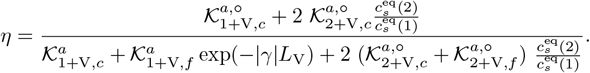

Taking the limit *L*_V_ → ∞ yields and upper bound for the replication efficiency,

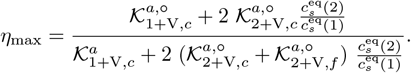

Provided that the ratio of equilibrium concentrations of dimers and monomers is comparable to the ratio of total concentration of dimers and monomers, *c*^eq^(2)*/c*^eq^(1) ≈ *c*^tot^(2)*/c*^tot^(1), we can write

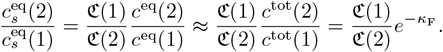

Therefore, the upper bound of the replication efficiency may be expressed as

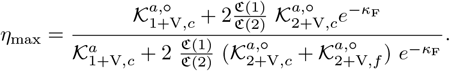

For the genome that is analyzed in the main text, 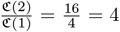. Comparison of the analytical approach (involving only ternary complexes) to the numerical solution (involving quaternary complexes) reveals that the contribution of quaternary complexes is indeed negligible (Main Text Fig. 5**Replication performance of single-length VCG pools containing monomers and dimers as feedstock. (A)** The pool contains a fixed total concentration of feedstock, 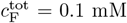, partitioned into monomers and dimers, as well as VCG oligomers of a single length, *L*_V_. The proportion of monomers and dimers can be adjusted via *κ*_F_, and the concentration of the VCG oligomers is a free parameter, 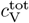. **(B)** Replication efficiency exhibits a maximum at intermediate VCG concentration in systems with (dashed blue curve) and without dimers (solid blue curve). The presence of dimers reduces replication efficiency significantly, as they enhance the ligation share of incorrect F+V ligations (dashed green curve). The panel depicts the behavior for *L*_V_ = 7 nt and *κ*_F_ = 2.3. **(C)** Optimal replication efficiency increases as a function of oligomer length, *L*_V_, and asymptotically approaches a plateau (dashed lines, Eq. (6equation.13)). The value of this plateau, 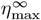, is determined by the competition between correct and false 2+V reactions, both of which grow exponentially with *L*_V_. Thus, 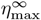 depends on the relative concentration of the dimers in the pool: the more dimers are included, the lower is 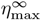. **(D)** Erroneous 1+V ligations are possible if the educt oligomer has a short overlap region with the template. The hybridization energy for such configurations is small, and independent of the length of the VCG oligomers (left). While 2+V ligations may produce incorrect products via the same mechanism (middle), they can also be caused by complexes in which two VCG oligomers hybridize perfectly to each other, but the dimer has a dangling end. The stability of these complexes increases exponentially with oligomer length (right).figure.caption.12)

### S11. ASYMPTOTIC FRACTION OF OLIGOMERS UNDERGOING EXTENSION BY MONOMERS

In pools containing only activated monomers, as well as non-activated dimers and VCG oligomers of a single length, we observe that the fraction of oligomers that can be extended by a monomer reaches an asymptotic value for high VCG concentrations. Notably, this asymptotic value is independent of the oligomer length.

The fraction of oligomers that are in a monomer extension-competent state can be computed based on the equilibrium concentration of complexes facilitating the extension of an oligomer by a monomer,

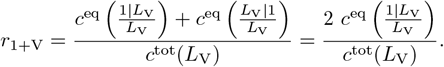

Note that, in principle, the monomer can be added to the 5^*′*^- or the 3^*′*^-end of the oligomer, leading to two contributions in the numerator, which are identical due to symmetry, 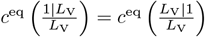.

Just like in the previous section, 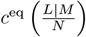 denotes the equilibrium concentration of all complexes allowing for the templated ligation of oligomers of length *L* to oligomers of length *M* if templated by oligomers of length *N*. We assume that ternary complexes are the dominant type of complexes facilitating templated ligation, and write the equilibrium concentration as,

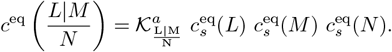

Therefore, the fraction of monomer-extendable oligomers reads,

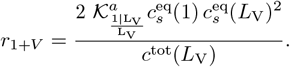

In the limit of high VCG concentration, almost all VCG oligomers form duplexes. Hence, the equilibrium concentration of free oligomers can be approximated by,

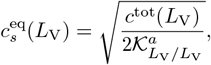

implying,

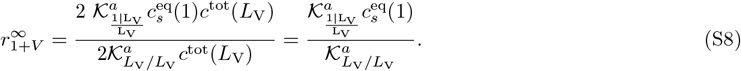

**FIG. S15.**
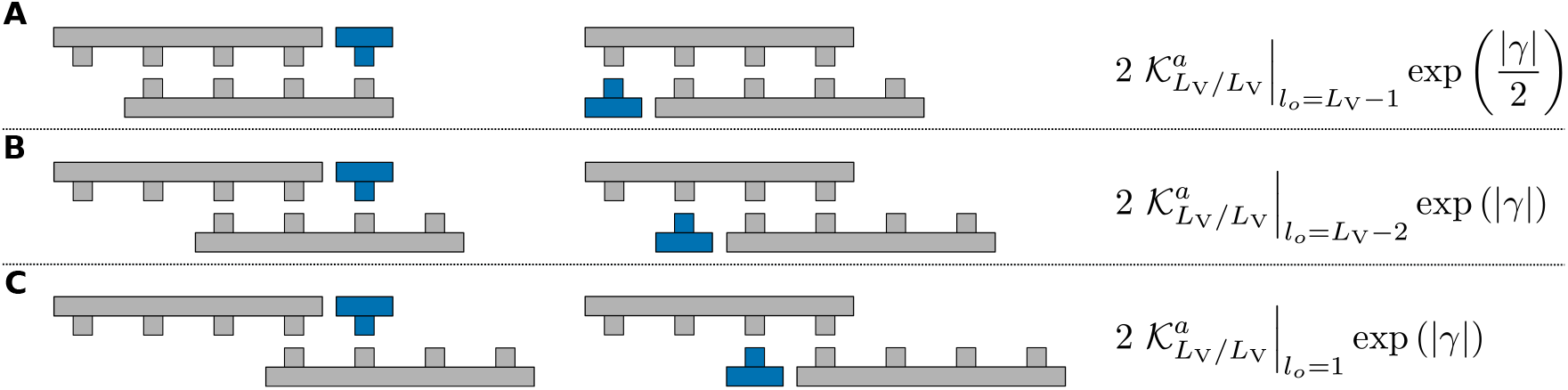
The effective association constants of reactive ternary complexes (complexes comprising three strands) can be computed based on the effective association constant of the duplexes. **A** If the hybridization region of the two VCG oligomers is *L*_V_ − 1 nucleotides long, the monomer hybridizes to the end (start) of the template strand. As the template has no dangling end, the energy contribution of the hybridizing monomer is *γ/*2. **B** and **C** For hybridization regions that are shorter than *L*_V_ − 1, but at least 1 nucleotide long, the energy contribution due to the added nucleotide is *γ*. In all cases, the prefactor 2 accounts for the two possible positions at which a monomer might be added.

As almost all monomers are free in solution, their equilibrium concentration may be approximated by the total concentration of monomers, 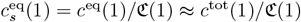.

From this representation of 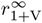, it is not clear yet why 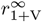 should be independent of *L*_V_. In order to derive an *L*_V_-independent expression for 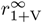, we need to express the association constant of ternary complexes, 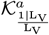, in terms of the duplex association constant, 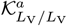. As illustrated in Fig. S15, the effective association constant of the triplex may be obtained by multiplying the effective association constant of the duplex by the binding affinity of the monomer. We need to distinguish the contributions of duplexes based on the length of their hybridization region: If the two strands in the duplex form a hybridization region that is as long as the oligomers, there is no free base pair for the monomer to hybridize; such duplexes must be excluded in the computation of the triplex association constant. Similarly, monomers that hybridize to the end of a strand have a different binding affinity than monomers hybridizing to the center of an oligomer (Fig. S15A vs. Fig. S15B-C). All in all, the effective triplex association constant reads,

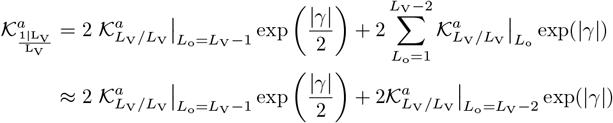

The notation 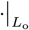 implies that only complex configurations with a specific overlap length *L*_o_ are included. The full effective duplex association constant may be expressed as

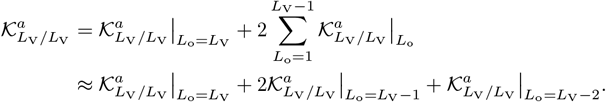

We write the duplex association constants in terms of the binding energy due to their overlap length,

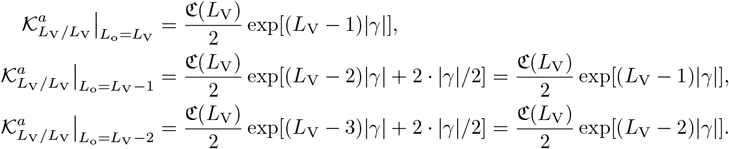

Therefore, the asymptotic ratio of monomer-extended oligomers reads,

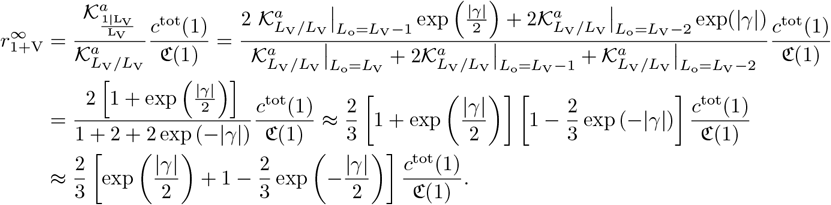

Addition of a single base pair will decrease the energy by the energy of a half nearest neighbor block, i.e., by *γ/*2. Thus, we assign *K*_*d*_(1) = exp (−|*γ*|*/*2), and find,

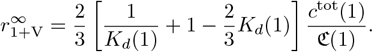

### S12. ANALYTICAL SOLUTION FOR EQUILIBRIUM CONCENTRATIONS IN MULTI-LENGTH VCG POOLS

We consider pools comprised of monomers, dimers and VCG oligomers of multiple lengths. For simplicity, we restrict ourselves to profiles with a uniform distribution of VCG oligomers, such that all VCG oligomers have the same concentration. Assuming that almost all oligomer mass is contained in single strands and duplexes, the mass conservation equations read

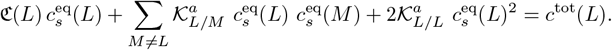

Note that there is a mass conservation equation for each oligomer length individually, i.e., we are dealing with a set of multiple coupled quadratic equations. We can make the system of equations dimensionless by introducing a dimensionless concentration 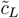,

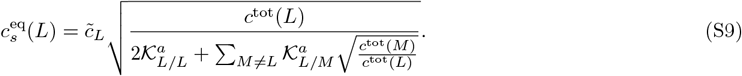

This rescaling is chosen as it is the solution to the (approximative) mass conservation equation in the high concentration limit,

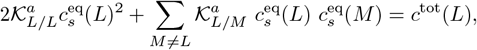

under the assumption that 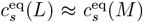. The latter assumption is reasonable given the concentration profile is uniform. We expect 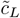 to be of order 1 in the high concentration limit.

Writing the full mass conservation equation in terms of the dimensionless concentration yields,

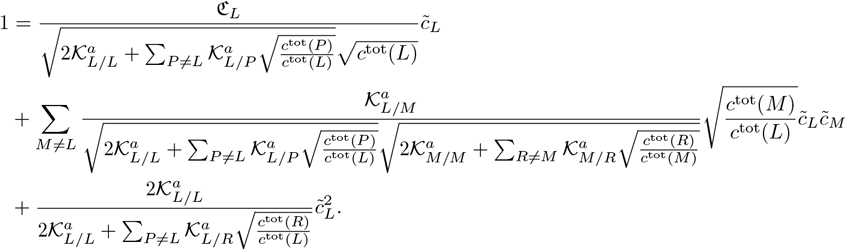

We drop all ratios of total concentrations, as we assume that the total concentration is the same for each oligomer length of VCG oligomer (uniform concentration profile),

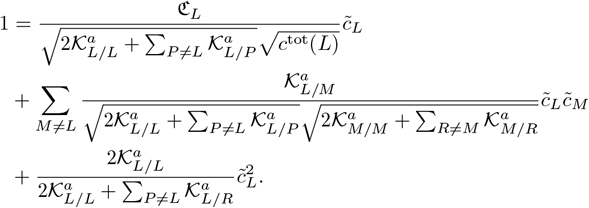

By introducing the dimensionless prefactors,

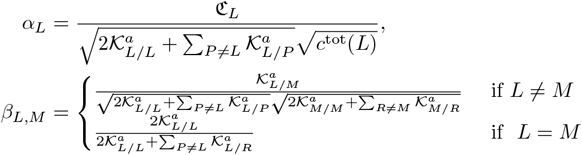

we can rewrite the mass conservation equation,

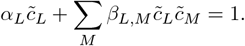

To solve for 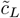, we make the assumption that the dimensionless equilibrium concentration is close to 1 for any length *L*. For this reason, we may assume that 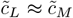. This assumption is crucial, as it allows us to de-couple the quadratic equations. Therefore, the mass conservation reads,

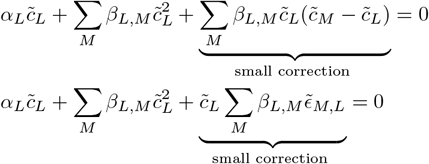

We can use this representation to compute the equilibrium concentrations recursively: We start the recursion with the assumption 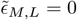, and compute 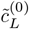 with this assumption,

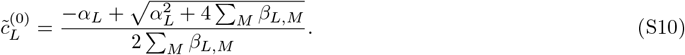

In the first recursion step, we compute 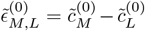, and solve the mass conservation equation for the equilibrium concentration,

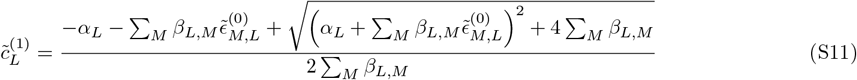

This scheme can be applied until the approximated values of 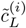 match the true values (obtained via numerical root finding) sufficiently well,

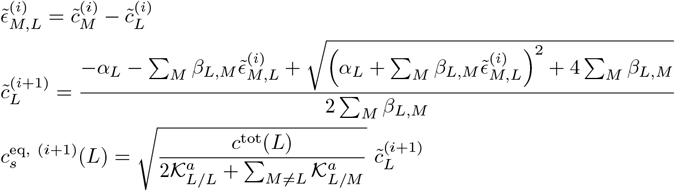

Note that the equilibrium concentration obtained after the (*i* + 1)-th iteration step 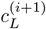 depends on the total concentration *c*^tot^(*L*) via the rescaling prefactor as well as via *α*_*L*_. It turns out that the approximation converges after about five iteration steps; the relative error between the true (numerically computed) equilibrium concentration and the approximation is already below 1% in the first iteration step.

**FIG. S16.**
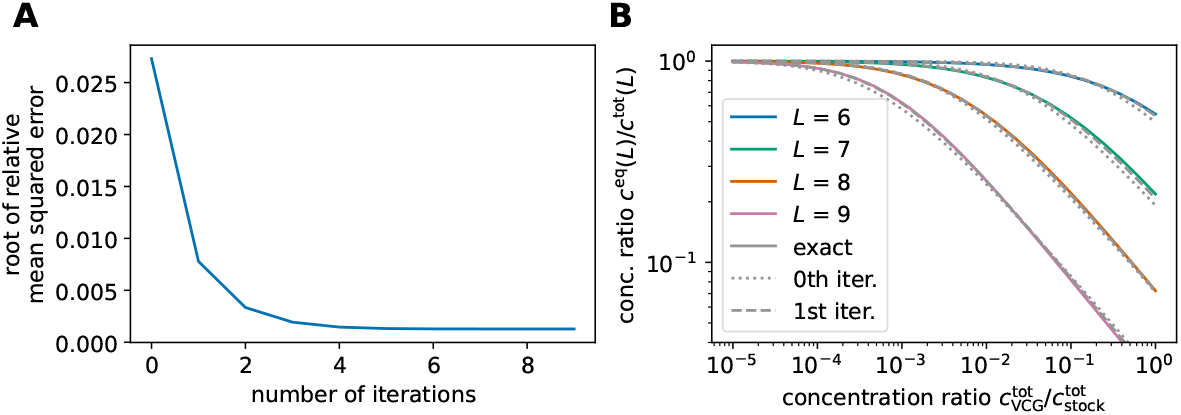
Comparison between approximate (analytical) and true (numerical) solution of the (de)hybridization equilibrium in multi-length VCG pools. **A** The approximation converges after roughly five iteration steps. The relative error drops below 1% in the first iteration step already. **B** Equilibrium concentrations as a function of total VCG concentration. Continuous lines show the numerical solution, while the dotted and dashed lines depict the approximation obtained in the zeroth or first iteration step respectively. The feedstock concentration is fixed, 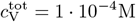.

### S13. THRESHOLD CONCENTRATION FOR THE INVERSION OF PRODUCTIVITY

In VCG pools that contain VCG oligomers of multiple lengths as well as activated monomers, we observe “inversion of productivity”: The fraction of short VCG oligomers in a monomer extension-capable complex configuration exceeds that of long oligomers. However, this is only the case for sufficiently high concentration of VCG oligomers. The threshold concentration that is necessary for oligomers of length *M* to exceed the monomer-extension fraction of oligomers of length *L* (assuming *M < L*) is set by the condition *r*_1+V_(*L*) = *r*_1+V_(*M*). To compute *r*_1+V_(*L*), we need to account for all possible complexes, in which an oligomer *L* can be extended by a monomers,

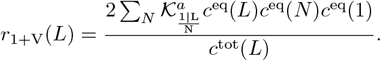

As we are considering a uniform concentration profile, all VCG oligomers have the same total concentration, i.e., *c*^tot^(*L*) = *c*^tot^(*M*). Thus, we can express the condition for the threshold concentration as

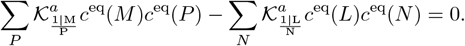

We combine this condition with the analytical approximation of the equilibrium concentration up to the first iteration step derived in Section S12, *cf*. Eqs. S9-S11. As the equilibrium concentrations 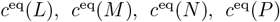 depend on the total concentration of VCG oligomers 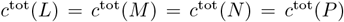, this yields an semi-analytical criterion for the threshold concentration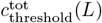, at which the fraction of monomer-extended *M*-mers exceeds the fraction of monomer-extended *L*-mers.

### S14. INVERSION OF PRODUCTIVITY USING PARAMETERS OF SYSTEM STUDIED BY DING *ET AL*

In their experimental study, Ding *et al*. focus on a genome of length *L*_*G*_ = 12 nt. In the VCG pool encoding the genome, Ding *et al*. include (Tab. S1 in [9])

- dimers with 11 different sequences,
- trimers with 20 different sequences, and
- tetramers up to 12-mers with 24 different sequences each.

Every oligomer sequence is included with a concentration of 1 µM (so-called 1x profile), adding up to a total concentration of 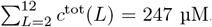. The feedstock for the replication is comprised of activated imidazolium-bridged homo-dinucleotides with a total concentration of 20 mM. Over a timescale of around 4 h, these homo-dinucleotides react with each other to form 6 additional imidazolium-bridged hetero-dinucleotides as well as activated mono-nucleotides (Fig. S1 in [9]).

We construct a genome that mimics the properties of the genome used by Ding *et al*., but obeys our genome construction principles outlined in the method section of the main text. To this end, we consider a genome of length *L*_*G*_ = 12 nt and a minimal unique subsequence length of *L*_*S*_ = 3 nt. This implies that our VCG pool contains

- dimers with 16 different sequences, and
- trimers up to 12-mers with 24 different sequences.

Our model does not include imidazolium-bridged dinucleotides explicitely. Instead, we assume that the concentration of activated mononucleotides in our model equals the total concentration of activated homo-dinucleotides (20 mM) used in the experimental study. The total concentration of non-activated oligomers is treated as a free parameters.

We find that the system exhibits inversion of productivity: For the entire considered range of concentration of non-activated oligomers, 10-mers are more likely to be extended by a monomer than 12-mers (Fig. S17). Moreover, 8-mers are more productive than 12-mers (provided the concentration of non-activated oligomers exceeds roughly 0.3 µM), and more productive than 10-mers (provided 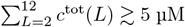). Thus, for the experimentally used concentration of non-activated oligomers (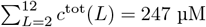, see vertical dashed line in Fig. S17), 8-mers are more productive than 10-mers, which are in turn more productive than 12-mers. Unlike in the experimental system, 6-mers are less likely than 8-mers and 10-mers to be extended by a monomer. We suppose that this difference can be attributed to the differences between the experimental setup and our theoretical model: (i) We choose a (slightly) different genome than Ding *et al*.. (ii) We model imidazolium-bridged dinucleotides as mononucleotides, as imidazolium-bridged dinucleotides only incorporate one mononucleotide at a time, just like activated mononucleotides. However, dinucleotides bind more stably to an existing complex than mononucleotides, which will affect the fraction of monomer-extended oligomers predicted.

**FIG. S17.**
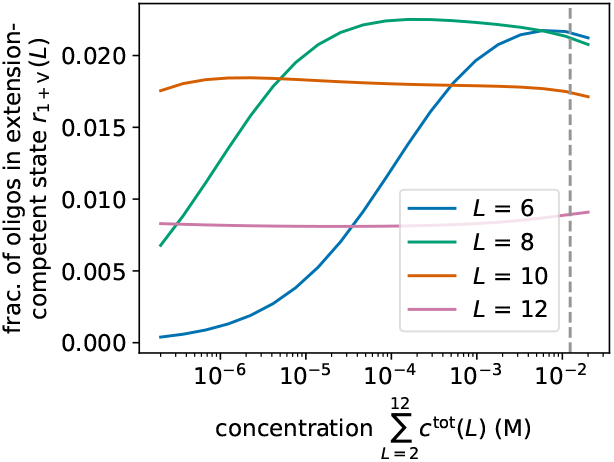
Replication performance of multi-length VCG pools, in which only the monomers are activated. The pool includes activated monomers (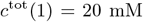, as well as non-activated oligomers of length *L* = 2 nt up to *L* = 12 nt with variable total concentration 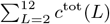. The system exhibits inversion of productivity: 10-mers are more likely to be in a monomer-extension competent state than 12-mers, 8-mers are more likely to be extended by monomers than 10-mers (for provided 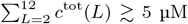). For the experimentally used concentration (vertical dashed line), 8-mers are more productive than 10-mers, and those are more productive than 12-mers. However, unlike in the experimental system, 6-mers are less productive than 8-mers and 10-mers.

## References

Ashwood B, Jones MS, Radakovic A, Khanna S, Lee Y, Sachleben JR, Szostak JW, Ferguson AL, Tokmakoff A. Thermodynamics and kinetics of DNA and RNA dinucleotide hybridization to gaps and overhangs. Biophys J. 2023; 122(16):3323–3339. 10.1016/j.bpj.2023.07.009.

Attwater J, Raguram A, Morgunov AS, Gianni E, Holliger P. Ribozyme-catalysed RNA synthesis using triplet building blocks. eLife. 2018; 7:e35255. 10.7554/eLife.35255.

Attwater J, Wochner A, Holliger P. In-ice evolution of RNA polymerase ribozyme activity. Nat Chem. 2013; 5(12):1011–1018. 10.1038/nchem.1781.

Becker S, Thoma I, Deutsch A, Gehrke T, Mayer P, Zipse H, Carell T. A high-yielding, strictly regioselective prebiotic purine nucleoside formation pathway. Science. 2016; 352(6287):833–836. 10.1126/science.aad2808.

Benner SA, Kim HJ, Carrigan MA. Asphalt, Water, and the Prebiotic Synthesis of Ribose, Ribonucleosides, and RNA. Acc Chem Res. 2012; 45(12):2025–2034. 10.1021/ar200332w.

Braunlin WH, Bloomfield VA. Proton NMR study of the base-pairing reactions of d(GGAATTCC): salt effects on the equilibria and kinetics of strand association. Biochemistry. 1991; 30(3):754–758. 10.1021/bi00217a026.

Chamanian P, Higgs PG. Computer simulations of Template-Directed RNA Synthesis driven by temperature cycling in diverse sequence mixtures. PLoS Comput Biol. 2022; 18(8):e1010458. 10.1371/journal.pcbi.1010458.

Ding D, Zhou L, Giurgiu C, Szostak JW. Kinetic explanations for the sequence biases observed in the nonenzy-matic copying of RNA templates. Nucleic Acids Res. 2021; 50(1):35–45. 10.1093/nar/gkab1202.

Ding D, Zhou L, Mittal S, Szostak JW. Experimental Tests of the Virtual Circular Genome Model for Nonenzymatic RNA Replication. J Am Chem Soc. 2023; 145(13):7504–7515. 10.1021/jacs.3c00612.

Duzdevich D, Carr CE, Ding D, Zhang SJ, Walton TS, Szostak JW. Competition between bridged dinucleotides and activated mononucleotides determines the error frequency of nonenzymatic RNA primer extension. Nucleic Acids Res. 2021; 49(7):3681–3691. 10.1093/nar/gkab173.

Göppel T, Obermayer B, Chen IA, Gerland U. A kinetic error filtering mechanism for enzyme-free copying of nucleic acid sequences. biorXiv. 2021; 10.1101/2021.08.06.455386.

Göppel T, Rosenberger JH, Altaner B, Gerland U. Thermodynamic and Kinetic Sequence Selection in Enzyme-Free Polymer Self-Assembly inside a Non-equilibrium RNA Reactor. Life. 2022; 12(4):567. 10.3390/life12040567.

Higgs PG, Lehman N. The RNA World: molecular cooperation at the origins of life. Nat Rev Genet. 2015; 16(1):7–17. 10.1038/nrg3841.

Ianeselli A, Salditt A, Mast C, Ercolano B, Kufner CL, Scheu B, Braun D. Physical non-equilibria for prebiotic nucleic acid chemistry. Nat Rev Phys. 2023; 5(3):185–195. 10.1038/s42254-022-00550-3.

Johnston WK, Unrau PJ, Lawrence MS, Glasner ME, Bartel DP. RNA-Catalyzed RNA Polymerization: Accurate and General RNA-Templated Primer Extension. Science. 2001; 292(5520):1319–1325. 10.1126/science.1060786.

Joyce GF. RNA evolution and the origins of life. Nature. 1989; 338(6212):217–224. 10.1038/338217a0.

Kervio E, Hochgesand A, Steiner UE, Richert C. Templating efficiency of naked DNA. Proc Natl Acad Sci USA. 2010; 107(27):12074––12079. 10.1073/pnas.0914872107.

Kervio E, Sosson M, Richert C. The effect of leaving groups on binding and reactivity in enzyme-free copying of DNA and RNA. Nucleic Acids Res. 2016; 44(12):5504–5514. 10.1093/nar/gkw476.

Kim HJ, Ricardo A, Illangkoon HI, Kim MJ, Carrigan MA, Frye F, Benner SA. Synthesis of Carbohydrates in Mineral-Guided Prebiotic Cycles. J Am Chem Soc. 2011; 133(24):9457–9468. 10.1021/ja201769f.

Kriebisch C, Burger L, Zozulia O, Stasi M, Floroni A, Braun D, Gerland U, Boekhoven J. Template-based information transfer in chemically fueled dynamic combinatorial libraries. Nat Chem. 2024; 16:1240–1249. 10.1038/s41557-024-01570-5.

Laurent G, Göppel T, Lacoste D, Gerland U. Emergence of Homochirality via Template-Directed Ligation in an RNA Reactor. PRX Life. 2024; 2(1):013015. 10.1103/PRXLife.2.013015.

Leu K, Kervio E, Obermayer B, Turk-MacLeod RM, Yuan C, Luevano JMJ, Chen E, Gerland U, Richert C, Chen IA. Cascade of Reduced Speed and Accuracy after Errors in Enzyme-Free Copying of Nucleic Acid Sequences. J Am Chem Soc. 2013; 135(1):354–366. 10.1021/ja3095558.

Leu K, Obermayer B, Rajamani S, Gerland U, Chen IA. The prebiotic evolutionary advantage of transferring genetic information from RNA to DNA. Nucleic Acids Res. 2011; 39(18):8135–8147. 10.1093/nar/gkr525.

Leveau G, Pfeffer D, Altaner B, Kervio E, Welsch F, Gerland U, Richert C. Enzyme-Free Copying of 12 Bases of RNA with Dinucleotides. Angew Chem Int Ed. 2022; 61(29):e202203067. 10.1002/anie.202203067.

Mathews DH, Disney MD, Childs JL, Schroeder SJ, Zuker M, Turner DH. Incorporating chemical modification constraints into a dynamic programming algorithm for prediction of RNA secondary structure. Proc Natl Acad Sci USA. 2004; 101(19):7287–7292. 10.1073/pnas.0401799101.

Mutschler H, Wochner A, Holliger P. Freeze–thaw cycles as drivers of complex ribozyme assembly. Nat Chem. 2015; 7(6):502–508. 10.1038/nchem.2251.

Müller UF, Bartel DP. Improved polymerase ribozyme efficiency on hydrophobic assemblies. RNA. 2008; 14(3):552–562. 10.1261/rna.494508.

Powner MW, Gerland B, Sutherland JD. Synthesis of activated pyrimidine ribonucleotides in prebiotically plausible conditions. Nature. 2009; 459(7244):239–242. 10.1038/nature08013.

Pressman AD, Liu Z, Janzen E, Blanco C, Müller UF, Joyce GF, Pascal R, Chen IA. Mapping a Systematic Ribozyme Fitness Landscape Reveals a Frustrated Evolutionary Network for Self-Aminoacylating RNA. J Am Chem Soc. 2019; 141(15):6213–6223. 10.1021/jacs.8b13298.

Prywes N, Blain JC, Del Frate F, Szostak JW. Nonenzymatic copying of RNA templates containing all four letters is catalyzed by activated oligonucleotides. eLife. 2016; 5:e17756. 10.7554/elife.17756.

Rajamani S, Ichida JK, Antal T, Treco DA, Leu K, Nowak MA, Szostak JW, Chen IA. Effect of Stalling after Mismatches on the Error Catastrophe in Nonenzymatic Nucleic Acid Replication. J Am Chem Soc. 2010; 132(16):5880–5885. 10.1021/ja100780p.

Robertson MP, Joyce GF. The Origins of the RNA World. Cold Spring Harb Perspect Biol. 2012; 4(5):a003608–a003608. 10.1101/cshperspect.a003608.

Rosenberger JH, Göppel T, Kudella PW, Braun D, Gerland U, Altaner B. Self-Assembly of Informational Polymers by Templated Ligation. Phys Rev X. 2021; 11(3):031055. 10.1103/PhysRevX.11.031055.

Scott WG, Horan LH, Martick M. The Hammerhead Ribozyme. Prog Mol Biol Transl Sci. 2013; 120:1–23. 10.1016/B978-0-12-381286-5.00001-9.

Sosson M, Pfeffer D, Richert C. Enzyme-free ligation of dimers and trimers to RNA primers. Nucleic Acids Res. 2019; 47(8):3836–3845. 10.1093/nar/gkz160.

Szostak JW. An optimal degree of physical and chemical heterogeneity for the origin of life? Philos Trans R Soc B. 2011; 366(1580):2894–2901. 10.1098/rstb.2011.0140.

Tjhung KF, Shokhirev MN, Horning DP, Joyce GF. An RNA polymerase ribozyme that synthesizes its own ancestor. Proc Natl Acad Sci USA. 2020; 117(6):2906–2913. 10.1073/pnas.1914282117.

Todisco M, Ding D, Szostak JW. Transient states during the annealing of mismatched and bulged oligonu-cleotides. Nucleic Acids Res. 2024; 52(5):2174–2187. 10.1093/nar/gkae091.

Todisco M, Radakovic A, Szostak JW. RNA Complexes with Nicks and Gaps: Thermodynamic and Kinetic Effects of Coaxial Stacking and Dangling Ends. J Am Chem Soc. 2024; 146(26):18083–18094. 10.1021/jacs.4c05115.

Toparlak OD, Sebastianelli L, Egas Ortuno V, Karki M, Xing Y, Szostak JW, Krishnamurthy R, Mansy SS. Cyclophos-pholipids Enable a Protocellular Life Cycle. ACS Nano. 2023; 17(23):23772–23783. 10.1021/acsnano.3c07706.

Walton T, Szostak JW. A Highly Reactive Imidazolium-Bridged Dinucleotide Intermediate in Nonenzymatic RNA Primer Extension. J Am Chem Soc. 2016; 138(36):11996–12002. 10.1021/jacs.6b07977.

Walton T, Szostak JW. A Kinetic Model of Nonenzymatic RNA Polymerization by Cytidine-5’-phosphoro-2-aminoimidazolide. Biochemistry. 2017; 56(43):5739–5747. 10.1021/acs.biochem.7b00792.

Welsch F, Kervio E, Tremmel P, Richert C. Prolinyl Nucleotides Drive Enzyme-Free Genetic Copying of RNA. Angew Chem Int Ed. 2023; 62(41):e202307591. 10.1002/anie.202307591.

Wetmur JG, Davidson N. Kinetics of renaturation of DNA. J Mol Biol. 1968; 31(3):349–370. 10.1016/0022-2836(68)90414-2.

Wochner A, Attwater J, Coulson A, Holliger P. Ribozyme-Catalyzed Transcription of an Active Ribozyme. Science. 2011; 332(6026):209–212. 10.1126/science.1200752.

Zhou L, Ding D, Szostak JW. The virtual circular genome model for primordial RNA replication. RNA. 2021; 27(1):1–11. 10.1261/rna.077693.120.

